# Injury improves tolerance to acute heat stress through induction of heat shock protein expression in the annelid *Pristina leidyi*

**DOI:** 10.64898/2026.08.30.748154

**Authors:** Corey W. Rennolds, James D. Nowotny, Alexandra E. Bely

## Abstract

Evolutionary patterns of regeneration ability in animals may reflect the relative costs of injury versus regeneration on organismal function. While the cost of injury seems obvious, the investment of resources and other physiological effects produced by regeneration may be considerable, and the effects of both injury and regeneration may depend upon myriad factors intrinsic to organisms or within their environments. Such effects and how they vary are not well understood. We investigated the physiological effects of amputation injury and subsequent regeneration of tissue in the annelid *Pristina leidyi*, using environmental stress tolerance as a relevant measure of performance. Unexpectedly, injury improved survival under heat stress, but improved heat tolerance was unrelated to changes in metabolic rate. We performed TagSeq to investigate the transcriptional basis of injury-induced heat tolerance and found that injured worms produced an exaggerated response to subsequent heat stress compared to either injury or heat stress alone. Both injury and heat stress commonly induced the expression of just two heat shock family proteins. Pharmacological inhibition of one of these proteins, mortalin, drastically reduced heat tolerance in both uninjured and injured worms and substantially impaired anterior tissue regeneration. We posit that injury-induced expression of mortalin and other cellular stress response factors briefly confers improved resistance to broad forms of stress. Our work demonstrates that injury may have unpredictable effects and points to a shared ancestral response to various forms of biological damage with a critical role in animal function during the regeneration process.

## Introduction

Mechanical injury is a threat to animal survival and function in nature. Injury damages tissue and organs, severing physical links and communication channels between cells, and disrupts function in myriad ways; injury breaks the barrier between internal and external environments, allowing circulatory fluids to escape and pathogens to enter; lost tissue may include critical biological materials and energy reserves; and severe, unmitigated injury may result in an animal’s death (Archie, 2013; Niethammer, 2016; Rennolds and Bely, 2023). Injury can also lead to lost function associated with a damaged tissue or appendage alongside a variety of secondary effects (Bernardo and Agosta, 2005; Juanes and Smith, 1995). The homeostatic challenge that injury imposes on animals at cellular or higher levels of biological organization is comparable in many ways to environmental stressors such as temperature, radiation, or hypoxia (Kassahn et al., 2009; Kultz, 2020a; Sulmon et al., 2015; Todgham and Stillman, 2013). The effects of stress may shape the structure of entire biological communities and influence the course of evolution (Kassahn et al., 2009; Kultz, 2020b; Sulmon et al., 2015), and injury may similarly affect biological processes beyond the scope of individual organisms if sufficiently prevalent in nature. Injury is quite common in natural populations (Lindsay, 2010), whether the source is predatory attacks, nonpredatory biotic interactions, or environmental accidents. Due to its potential critical consequences, injury is a significant stressor that animals are likely to avoid or mitigate whenever possible (Rennolds and Bely, 2023).

As with other forms of stress, animals have evolved a range of responses to injury that serve to reduce its detrimental effects. Among the most dramatic of these responses is regeneration, the process by which damaged or lost tissue is restored, either partially or totally, to its previous form and function. Regeneration involves a range of physiological and genetic mechanisms which may be conserved, convergent, or highly distinct between species and contexts (Bely and Nyberg, 2010; Lai and Aboobaker, 2018). All forms of regeneration are united, however, by requiring energy, materials, and time (Mack and Bely, 2025). Since these variables are limited in organisms, regeneration often proceeds at direct or indirect cost to other biological functions. The effects of regeneration on reproduction and non-regenerative growth have perhaps been best described in animals broadly (Archie, 2013; Bernardo and Agosta, 2005; Henry and Hart, 2005; Maginnis, 2006; Sepulveda et al., 2008). However, there are more species that can regenerate than those in which the costs of regeneration have been studied. As such costs might hinder animal function and subsequently reduce fitness, the limited knowledge of these costs and their variation in highly diverse species and contexts hinders our ability to understand the broader ecological and evolutionary relevance of regeneration and to predict which mechanisms underpinning regeneration are potential targets of selection.

A major challenge in understanding the effects of regeneration on organisms is distinguishing such effects from those caused directly by injury itself. The costs and other effects of injury are rarely straightforward or consistent: myriad factors such as life history, nutritional status, or environmental conditions can modulate the response to injury, such as regeneration speed or quality, or trade-offs with other biological processes (Maginnis, 2006; Rennolds and Bely, 2023; Rennolds and Bely, 2024). The nature of the injury itself may also influence the response, particularly if the damaged structure is functionally critical or contains energy reserves, as in lizard tails (Bernardo and Agosta, 2005; Smyth, 1974). Injury occurs under variable conditions in nature, and thus testing the effects of injury alone without introducing variation, such as in the abiotic environment or the exact location of injury, can obscure the complexity in injury responses and thus distort our understanding of its role in shaping biological processes. Without adequately distinguishing the effects of injury from the effects of resource consumption or other changes induced by regeneration or recovery more broadly, we will be unable to fully understand why animals evolved such diverse responses to injury, including diversity in regenerative ability.

In this study, we investigated how mechanical injury and regeneration affect organismal physiology in the annelid *Pristina leidyi* (Smith, 1896). We made an effort to distinguish the effects of initial injury versus subsequent regeneration by measuring physiological endpoints at different times in the post-injury period. *P. leidyi* is a suitable organism for studying regeneration due to its rapid whole-body regeneration, iterative anatomy, transparency, and experimental tractability (Bely, 2022). *P. leidyi* typically regenerates large portions of its body following transverse amputation within five days, including at both the anterior and posterior ends (Zattara and Bely, 2011). Posterior regeneration is always epimorphic, while loss of more than four anterior segments induces epimorphic regeneration of up to four segments coupled with morphallactic remodeling of existing segments into new segmental identities (Ozpolat and Bely, 2016; Zattara and Bely, 2011). Anterior and posterior ends possess different complements of organs and associated functions; therefore, the physiological effects of losing these tissues and the costs of restoring them are likely to differ from one another, and we investigated these impacts separately in this study.

We first assessed the effects of injury and regeneration in *P. leidyi* on the level of whole organismal physiology. *P. leidyi* usually survive amputations of significant body portions and regenerate without difficulty. Therefore, we decided to assess how injury and regeneration affect environmental stress tolerance, as this is expected to be a physiologically demanding challenge that at extreme levels or extended exposure is lethal and thus can reveal otherwise subtle changes in organismal condition, by measuring survival and oxygen consumption. While stress tolerance was reduced after regeneration, we unexpectedly observed a mild improvement in stress tolerance shortly after injury, and neither of these findings were related to changes in aerobic respiration. We followed up on the finding that injury improves stress tolerance by measuring gene expression following injury and heat stress, both separately and in combination, via 3’ tag-based bulk RNA-seq (TagSeq). Common to all injury and heat stress conditions was the upregulation of two heat shock family proteins (HSPs), homologs of *hsp10* and *mortalin*. Through targeted pharmacological inhibition, we show that mortalin function is critical for both regeneration success and acute heat tolerance, providing evidence of a mechanistic connection between the response to injury and environmental stress.

## Methods

### Culturing and obtaining experimental animals

*Pristina leidyi* originally purchased from Carolina Biological Supply (sold as *Stylaria)* were cultured at room temperature (23 ⁰C) in glass bowls (12 cm diameter) filled approximately halfway (∼150 mL) with artificial spring water (1% artificial seawater) (ASpW). Strips of brown paper towels were provided as substrate. Cultures were fed once weekly with 10 mg powdered Spirulina. Half-volume water changes were administered weekly.

To obtain animals for experiments, worms that appeared healthy were washed several times in clean ASpW and held together in a glass bowl for three days, allowing worms to clear their gut contents. Worms that appeared damaged, showed signs of aging or stress (e.g., shrunken, dense chloragogenous pigmentation), or had visible fission zones were removed after three days. Worms were randomly selected from this remaining pool of animals for use in the following experiments.

### Amputation

Individuals were first anesthetized in 0.05 mM nicotine and then pipetted onto glass slides, where either the anterior or posterior end was removed with a scalpel. Uninjured worms were anesthetized for approximately equivalent times. Worms were amputated between segments 6 and 7 (anterior injury), between segments 16 and 17 (posterior injury), or not injured (controls). Amputated fragments (the shorter piece following amputation) were discarded.

*P. leidyi* forms a wound epithelium over the site of injury and begins construction of the blastema, the undifferentiated cell mass that develops into regenerated tissue, by approximately 24 hours post-injury (Zattara and Bely, 2011). At roughly five days, regeneration of either anterior or posterior tissue is typically complete. We therefore used one and five days post amputation (1 dpa and 5 dpa) as measurement time points throughout the study.

### Thermal stress experiments – determining acute thermal tolerance range

To choose suitably stressful temperatures, we first determined the thermal tolerance range of *P. leidyi*. After amputation, worms were placed singly in 0.5 mL microcentrifuge tubes filled completely with fresh ASpW. Tubes were placed in low light at room temperature (23°C) for 1 dpa or 5 dpa without disturbance. The 1 dpa treatment was intended to measure changes in thermal tolerance resulting from injury and the loss of tissue prior to any significant regeneration. The 5 dpa treatment was intended to measure the impact of regeneration on thermal tolerance. Uninjured controls in the 5 dpa treatment group allowed us to determine the relative impact of fasting on thermal tolerance, as worms had no access to food during any point of the experiment.

At either 1 or 5 days, worms were checked for mortality (i.e., resulting from injury) (>95% of worms survived initial injury). Their tubes were then randomly arranged in a water bath inside an adjustable low temperature incubator (for temperatures below 23°C) (146E, Fisher Scientific, Waltham, MA) or in flooded wells of a thermal incubator (for temperatures above 23°C) (Isotemp 125D, Fisher Scientific, Waltham, MA). A range of temperatures (5, 7, 15, 23, 35, 39, 40°C) were tested to determine *P. leidyi*’s acute tolerance range. Tubes were also placed in a water bath on a benchtop in low light at room temperature (23°C) as a thermal control. Temperatures were regularly verified by an analog thermometer to be ±0.5°C from the set point. Tubes remained undisturbed in these conditions for 48 hours.

At 48 hours, tubes were removed from experimental temperature and left to sit at room temperature for approximately twenty minutes. Mortality was then assessed by looking for body movement or gut peristalsis within fifteen seconds of observation. Worms that exhibited neither or showed clear signs of decomposition were considered dead. Most dead worms were in various degrees of decomposition at the time of assessment. Survival was scored as a percentage of the number of worms alive at this check point (e.g., 7 of 10 worms alive = 70% survival). This experiment was replicated *N* = 5 times per treatment combination, using *n* = 10 worms per treatment combination per replicate.

### Thermal stress experiments – survival assay at thermal extremes

Worms were amputated, placed in tubes, and allowed to sit for one (1 dpa) or five days (5 dpa) as described previously. All other procedures up to and including moving tubes to incubators were also identical. Only two temperatures were tested, representing the lowest observed high (39°C) and highest observed low (5°C) temperatures at which death was significant (at or near 0% survival) at 48 hours on average across all injury treatments. Mortality of all worms was scored approximately every 3 hours until all worms in each run (*n* = 15 worms per treatment combination) were dead.

### Salinity stress experiments – acute salinity tolerance range-finding and survival assay

Salinity tolerance was used to assess whether effects of injury and regeneration on thermal tolerance extended to the broader organismal stress response. Amputations and pre-stress incubation periods were conducted as described previously, but worms were placed singly in filled (∼1 mL fresh ASpW) wells of 24-well tissue culture polystyrene plates. At 1 dpa or 5 dpa, worms were moved to wells filled with water of varying salinities (1-10 ppt (parts per thousand), in increments of 1 ppt) and placed in low light. Survival was scored at 48 hours. Survival was 100% at 9 ppt and 0% at 10 ppt, so we repeated this assay with a narrower range of salinities (9.1-9.9 ppt, in increments of 0.1 ppt).

The lowest observed salinity at which mortality was substantial (<50% survival) at 48 hours across all injury treatments was determined to be 9.5 ppt. To determine the difference in survival between injury treatments at a finer temporal scale, an identical experimental design was used as described above for survival at thermal extremes, but at this elevated salinity rather than extreme temperatures, and scoring of survival was identical (*n* = 15 worms per treatment combination).

### Respirometry

Worms were amputated, placed singly in filled wells of 24-well plates, and allowed to sit for 1 (1 dpa) or 5 days (5 dpa) as described previously.

To measure how injury at different body sites at different stages of recovery affected the sublethal response to thermal stress, we measured oxygen uptake (assumed hereafter to be equivalent to consumption) under exposure to 35°C, the highest temperature at which no mortality occurred in our range-finding experiment, and 10°C, which we incidentally found to be nonlethal (approximately 100% survival) to worms for at least several days when testing culture temperatures for an unrelated study (C.W. Rennolds, unpublished observations). We used a complete optical microplate system with an 80 μL well volume (Loligo Systems, Viborg, Denmark). The microplate was prepared for each experimental run by first immersing the wells in fresh ASpW for >20 minutes. The ASpW was then removed, and the system was calibrated using fresh ASpW that was first allowed to equilibrate to the relevant trial temperature and bubbled with air for >20 min (100% O_2_) or a solution of 20 g L^-1^ Na_2_SO_3_ in fresh ASpW (0% O_2_), according to the manufacturer’s instructions. The plate was then rinsed, and wells were filled with fresh ASpW. Worms were randomly assigned to wells and transferred, removing any air bubbles formed during the transfer. Each run with the respirometer included eighteen worms (six worms per injury location (anterior, posterior, or uninjured)) and six wells used as blanks, and only included worms from the same recovery stage and processed batch (i.e., injured at the same time). After removal of bubbles, the plate was sealed with a sheet of translucent adhesive PCR film (Thermo Fisher Scientific, Waltham, MA) and placed on the microplate reader. A silicone gasket and compression block were placed on top of the plate to ensure a continuous seal. The microplate apparatus was then moved to either a low-temperature incubator at 10°C (146E, Fisher Scientific, Waltham, MA), an oven at 35°C (6241, Fisher Scientific, Waltham, MA), or in low light at room temperature (23°C) as a thermal control. Recording was then initiated, and oxygen measurements were taken every minute until all samples reached 70% air saturation. Data were recorded in the included MicroResp™ software (v1). Worms were then removed and checked for survival; runs with more than one worm death were not used in analysis. Surviving worms were removed and imaged for body volume measurements, as described below.

Respirometry of worms under differing salinities (0.35 “ASpW” versus 6 ppt) was identical to the procedure described above with the following exceptions: all respirometry was conducted exclusively at room temperature; calibration used either ASpW or 6 ppt water as appropriate to the trial; and worms were first rinsed in 6 ppt before transfer to the respirometer to minimize dilution for all 6 ppt trials.

### Imaging and calculations of mass and oxygen consumption rate

Due to the logistical constraints of directly measuring individual body mass in *P. leidyi*, mass was approximated through measurements of individual volume. Individual worms were pipetted onto glass slides and anesthetized with 0.05 mM nicotine until movements ceased, and then worms were imaged under a 2.5x objective with a Zeiss Axioplan 2 microscope and AxioVision 4.8 image processing software. Individual length and width were measured in Fiji (Schindelin et al., 2012). Body length (*L,* mm) spanned from the posterior tip to just prior to the beginning of the anterior proboscis (Fig. S1). Body width (*W,* mm) was determined by averaging three width measurements, drawn just prior to where the anterior and posterior ends began to taper, and across the approximate center of the worm. Volume was calculated using the equation of a cylinder (*V=π_r_*^2^*h*), substituting *W*/2 for *r* and *L* for *h*. Body mass was then calculated by multiplying volume (*V*) by the density of water (*ρ*) at room temperature (*ρ* = 0.997773 kg L^-1^) (Colt, 2012).

Calculation of absolute oxygen consumption rates was performed in the MicroResp™ software. Phase values in each well were converted to oxygen concentration in mmol L^-1^. Data from the first 30 minutes of each run were not included in calculation of oxygen consumption rate. Only oxygen data between 90% and 70% air saturation were used in the calculation of the slope of oxygen concentration, deemed the metabolic oxygen consumption rate (MO_2_) (nmol hr^-1^). Linear regression of oxygen concentration within these limits was performed for each worm, and those with R^2^ <0.95 (23°C and 35°C; ASpW and 6 ppt) or <0.90 (10°C) were not used in analysis. MO_2_ of the blank well means were subtracted from those of the experimental wells for each run in order to account for background factors, yielding absolute MO_2_ for each worm. These values were divided by calculated body mass of each worm to obtain the relative metabolic rate (nmol hr^-1^ mg^-1^). *Q*_10_ coefficients for each recovery stage (1 dpa, 5 dpa) and injury condition (anterior, posterior, uninjured) combination across the range of temperatures (10, 23, 35°C) or salinities tested (0.35, 6 ppt) were calculated using 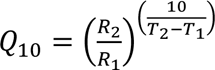 where *R*_x_ is MO_2_ at temperature or salinity *T*_x_ as appropriate.

### TagSeq procedure

We performed a gene expression study using TagSeq to assess the transcriptomic response to injury and thermal stress at 1 dpa following anterior or posterior amputation. TagSeq differs from standard RNA-seq by focusing sequencing on the 3’ end of each transcript, producing one read per transcript. This serves as a cost-effective methodology for differential gene expression studies when a reference genome or transcriptome is available (Lohman et al., 2016). Our experiment included four injury treatments, including two amputation conditions and two tissue controls (see below), replicated across two temperature conditions (23, 35°C) for a total of eight treatment combinations. This matrix was replicated *n* = 3 times.

Worms were amputated and placed in tubes as described previously. At approximately 24 hours, worms were either left at room temperature (23°C) or moved to a thermal incubator at 35°C for six hours. Tubes were then removed, worms were checked visually to confirm survival, and then worms were pooled in a single tube in groups of 10-12 of the same injury type per tube. Uninjured worms were amputated as described previously in groups of 10-12 per injury location (anterior or posterior) immediately following removal from their individual tubes and then quickly (< 5 min) collected in single tubes. These worms served as tissue loss controls, to account for potential differential gene expression in body segments not present in amputated worms and unrelated to injury or thermal stress, such that each injury group had its own control group. After transfer of worms to single tubes, these were snap frozen in liquid nitrogen and moved to -80°C until RNA extraction.

Tubes containing worms were administered 50 µL TRIzol (Thermo Fisher Scientific, Waltham, MA), vortexed, frozen at -80°C for approximately 15-30 minutes, and thawed. 1 µL polyacryl carrier and 150 µL TRIzol was added to each sample, which was then mixed and incubated a few minutes at room temperature. Then, 40 µL chloroform was added, tubes were incubated 15 minutes at room temperature, and samples were then centrifuged at maximum speed at 4°C for 15 minutes. The upper aqueous phase of each tube was carefully removed and mixed with 100 µL isopropanol, incubated for 30 minutes, and centrifuged at maximum speed at 4°C for 15 minutes. Supernatant was removed and precipitated RNA was washed with 75% ethanol and centrifuged at room temperature for 5 minutes. RNA pellets were air dried and then resuspended in 30 µL of RNase-free water. Total RNA was analyzed on a NanoDrop 8000 (Thermo Fisher) spectrophotometer for yield and purity. Yield was confirmed and integrity assessed by running RNA on a BioAnalyzer 2100 (Agilent Technologies, Santa Clara, CA). RNA was kept frozen at -80°C until library preparation.

Library construction and sequencing was carried out at the Genomic Sequencing and Analysis Facility at the University of Texas at Austin. TagSeq libraries were constructed from total RNA according to published methods (Lohman et al., 2016). In brief, RNA was heat-fragmented at 95°C for 2.5 minutes and transcribed into first-strand cDNA using SMARTScribe™ reverse transcriptase (Takara Bio Inc., Mountain View, CA) and template switching oligos (Integrated DNA Technologies, Coralville, IA). cDNA was purified using AMPure XP beads (Beckman Coulter, Brea, CA) and PCR amplified for 18 cycles. PCR products were further purified, indexed (Integrated DNA Technologies), purified once more, quantified using PicoGreen (Thermo Fisher), pooled equally, and size-selected to 350-550 base pairs (bp) with a BluePippin electrophoresis system (Sage Science, Beverly, MA). Libraries were sequenced on a NovaSeq 6000 SR100 lane (Illumina, San Diego, CA).

### TagSeq analysis

Sequenced libraries ranged from approximately 4.7 to 22.4 million raw reads per sample. Custom scripts (https://github.com/z0on/tag-based_RNAseq) were used to collapse duplicate reads and trim Illumina TagSeq adapter contaminants (one of four possible adapters + 24 downstream bp) and 3’ poly(A) tails (8 or more bp). Reads less than 20 bases long after trimming were discarded. After deduplication and trimming, libraries ranged from approximately 1.1 to 5.5 million reads per sample. Trimmed and filtered reads were mapped and quantified using ‘Salmon’ v1.1 (Patro et al., 2017) with a *k*-mer size of 19. Reads were mapped to the *P. leidyi* reference transcriptome prepared by Alvarez-Campos et al. (NCBI GEO accession GSM7225503; Alvarez-Campos et al., 2024). Mapping rates ranged between approximately 83- 89%. Full transcript accession IDs are formatted in this transcriptome as “UnnamedSample_HQ_transcript/####”, but for simplicity throughout we refer to these accession IDs only by their identifying number (e.g., UnnamedSample_HQ_transcript/79043 referenced as 79043).

Differential expression analyses were performed using ‘edgeR’ v3.34.1 (Robinson et al., 2010) in the R computing environment (Team, 2024). Low abundance transcripts were filtered using the filterByExpr function to maximize DET discovery, leaving 34,351 transcripts included in subsequent analysis, and library sizes were normalized using the calcNormFactors function. Sample read quantifications were fitted to a negative binomial generalized linear model with quasi-likelihood test using a combined injury and temperature condition factor with the glmQLFit function and robust dispersion estimation, and DETs were identified from pairwise treatment comparisons with the glmQLFTest function.

### Functional annotation

The Trinotate v3.2.2 pipeline (Bryant et al., 2017) was used to functionally annotate the *P. leidyi* IsoSeq transcriptome. TransDecoder v5.5.0 (https://github.com/TransDecoder) was first used to predict open reading frames (ORFs) and protein coding regions using the default minimum length of 100 amino acids. Homology to known proteins was used as ORF retention criteria by performing a BLASTp search against the UniRef90 database (Suzek et al., 2015) with BLAST v2.12.0 (Altschul et al., 1990). Assembled transcripts and predicted proteins were then used to perform homology searches against the UniProtKB/Swiss-Prot database (UniProt, 2019) with BLASTx and BLASTp, respectively, using the default *E*-value cutoff of 0.001. Predicted proteins were also used to search for protein domains with HMMER v3.3.2 (Finn et al., 2011) against the Pfam database (Mistry et al., 2021). Results were loaded into a Trinotate SQLite Database, and all hits that did not meet the E-value threshold of 0.001 were removed. For transcripts with multiple valid hits, only the top hit was retained for annotation of gene expression data. If there were discrepancies between BLASTx and BLASTp hits, the one with the lower E-value was used for identification in included tables. Pfam hits, if available, were used if none were returned by BLASTx or BLASTp.

### qPCR

The experimental procedure for quantitative polymerase chain reaction (qPCR) was similar to that for TagSeq. There were eight treatment groups in total: anteriorly amputated worms at 6 and 24 hpa; their corresponding tissue controls, as discussed above; uninjured worms exposed to 35°C in an incubator, as previously described, for either three or six hours; and their corresponding controls at each time point, maintained at room temperature (23°C). Amputations and thermal exposure were administered in similar manner as described previously, as were the preparation of anterior tissue controls just prior to snap freezing. Each replicate consisted of 4-5 worms pooled in a single tube, which were snap frozen and stored at -80°C until RNA extraction, which was carried out in identical fashion as above. Each experimental condition was replicated five times. Resuspended RNA was treated with TURBO DNase (Invitrogen, Thermo Fisher) according to the manufacturer’s protocol. Reverse transcription into cDNA was done with the Superscript III First-Strand Synthesis Kit (Invitrogen) with the same volume of template RNA (7 μL) per sample per reaction and random hexamers. cDNA was stored at -20°C for up to several weeks until used for qPCR.

Primer pairs (Table S20) were designed for three transcripts in our qPCR experiment: one housekeeping gene, *GAPDH*, and two transcripts found to be commonly upregulated under all treatment conditions of the TagSeq experiment: the 10 kDa HSP *hsp10* and the 70 kDa HSP family member *mthsp70*, also known and hereafter referred to as *mortalin*. Homology was verified with NCBI’s online BLASTX search tool. Primers were designed with the NCBI Primer-BLAST tool with 55% GC content for amplicons between 100-140 bp. Primers were synthesized by GeneWiz (Azenta Life Sciences, South Plainfield, NJ) and resuspended to a stock concentration of 100 μM in Milli-Q water, from which 10 μM working solutions of combined forward and reverse primers were prepared.

qPCR reactions were prepared as: 1 μL cDNA, 1 μL of 10 μM primers mix, 3 μL RNase-free water, and 5 μL 2X SYBR Green I Master Mix (Roche Diagnostics, Basel, CH). Reactions were prepared in 96-well plates, sealed with transparent PCR film, and run on a Roche LC480 with the cycling parameters: 95°C for 5 min, then 40 cycles of 95°C for 15 s and 60°C for 75 s. The uninjured or 23°C condition was used as the control at each time point for the anteriorly injured and 35°C conditions, respectively. Relative expression (log_2_ fold change) was calculated via ΔΔCt per replicate (Livak and Schmittgen, 2001). Expression of *hsp10* and *mortalin* were normalized to *GAPDH* expression. One outlier replicate in the 23°C, 3 h group was identified in the post-reaction diagnostic report and removed from subsequent analysis.

### Drug trials

1-Ethyl-2-[[3-ethyl-5-(3-methyl-2(3H)-benzothiazolylidene)-4-oxo-2- thiazolidinylidene]methyl]-pyridinium chloride (MKT-077; Sigma-Aldrich, M5449) powder was diluted in anhydrous dimethyl sulfoxide (DMSO) to a stock concentration of 10 mM. The stock solution was further diluted in ASpW to a range of working concentrations (1-20 μM), which were used initially to determine an optimal concentration that would produce a clear regeneration phenotype without rapid death. That concentration was determined to be 4 μM (J.D. Nowotny, data not shown), resulting in a final DMSO concentration of 0.04%. Control worms were kept in ASpW with 0.04% DMSO throughout the experiment. Control and MKT-077-treated worms were kept individually in single wells of a 96-well plate in 200 μL volumes. Regenerating worms were placed in their respective treatment within 30 s after amputation, which was performed anteriorly in identical fashion as previously described, as were exposures to temperature (23°C or 37°C). Medium was replaced every two days, during which < 20% evaporation was observed in the wells. On day 6 of the experiment, regenerating worms were anesthetized and imaged as described above. Length of regenerated tissue was measured as the distance between the anterior-most end of the pigmented gut (the approximate site of amputation in all cases) and the base of the proboscis (or to the end of the head in worms without a proboscis). The drug treatment experiment was performed in two blocks, and only worms in the second block were imaged. Final survival (of worms in both blocks) was scored on day 7.

### Statistical analysis

We performed all statistical analysis in R. We tested differences in thermal tolerance and salinity tolerance after fixed 48-hour exposure across a range of temperatures or salinities, between injury recovery stages and injury location via multiple factorial ANOVA and Tukey’s-adjusted post-hoc comparisons. A QQ plot of survival was symmetrical with minor heteroscedasticity in the residuals, and these were not corrected by transformation and did not affect analysis. To compare survival within these groups at thermal extremes or high salinity at higher temporal resolution, we used Kaplan-Meier survival analysis using Cox proportional hazards models (Cox, 1972) on interval-censored data with the ‘icenReg’ package (Anderson-Bergman, 2024) and specifying 100 bootstrapped samples. We used Type III SS ANCOVA to test the effects of injury location, recovery stage, and temperature or salinity on MO_2_, setting calculated worm mass as the covariate, using the ‘car’ package (Fox and Weisberg, 2019). We detected one extreme outlier in MO_2_, but its removal made no difference in the statistical outcomes, so we did not exclude it from analysis. As the raw temperature experiment MO_2_ data violated normality (Shapiro-Wilk test *P* < 0.001) and heteroscedasticity (increasing residuals, Levene’s test *P* < 0.001) assumptions, we initially log-transformed MO_2_, which improved each. This led to a significant (*P* < 0.05) interaction between injury and timepoint. However, we detected no significant differences in Tukey’s post-hoc comparisons between injury × timepoint treatments, and all other ANCOVA terms remained significant or nonsignificant as prior to transformation. Because of this, we concluded that transformation did not substantially alter our analysis, and therefore the following results are discussed in reference to the raw MO_2_ data. Raw salinity experiment MO_2_ data also violated normality and heteroscedasticity checks, and so we log-transformed these values. We then detected two extreme outliers in MO_2_ and removed these from further analysis, which satisfied the statistical assumptions. This led to two additional significant post-hoc comparisons based on the significant three-way interaction versus when data were not log-transformed. Due to this, all discussion of respirometry results in the salinity experiments are in reference to log-transformed MO_2_ data, although readers should note that these results are not considerably different overall than analysis of the raw data. The effect of recovery stage and injury location on worm mass was tested via Type II SS ANOVA. qPCR was analyzed via two-way ANOVA with ΔCt as the response variable and treatment (injury condition or temperature) and time point as the explanatory variables. Treatment and time point did not interact significantly in these models. Length of regenerated tissue in the drug experiments was analyzed via Type III SS ANOVA with temperature and drug condition as explanatory variables. Survival in the drug experiments was analyzed via logistic regression with Firth’s modification (Heinze and Schemper, 2002) using the ‘logistf’ package (Heinze et al., 2025), setting survival at day 7 as a binary variable as affected by injury condition, drug treatment, and temperature, with block as a random effect. To perform Tukey’s-adjusted post-hoc comparisons in all cases, we used the ‘lsmeans’ package (Lenth, 2016), except for the drug experiments, for which we used the ‘emmeans’ package (Lenth and Piaskowski, 2026).

## Results

### Injury improves survival under acute heat stress

We first investigated the effects of injury and regeneration on thermal tolerance. We assayed the effects of injury at 1 day post amputation (dpa), a timepoint when worms have sealed the wound but have not yet formed a visible blastema, and assayed the effects of regeneration at 5 dpa, a timepoint when worms have completed regeneration. We performed an initial range-finding experiment, exposing worms for 48 h to a range of temperatures, to determine the acute thermal tolerance range of *P. leidyi* and whether it differed following anterior or posterior amputation. We found that uninjured, recently injured (1 dpa), and regenerated worms (5 dpa) show high (>96%) survival following short term exposure to a broad thermal range between 15 and 35°C. However, survival declines sharply outside of this range, with no survival at 5 or 40°C (Fig. S2). In this assay, we detected no effect of amputation site (anterior vs. posterior) or recovery timepoint (1 vs. 5 dpa) on survival at any temperature (multiple ANOVA).

We then performed a continuous exposure experiment at three temperatures (5, 23, and 39°C) to reveal potential subtle effects of injury and regeneration on thermal tolerance. In this experiment, we tested whether survival under continuous exposure to extreme temperature declined at different rates. We chose to run these trials at 5 and 39°C as these were the lowest and highest temperatures at which mortality in our range determination experiment was substantial (<50% in all groups: 0% at 5°C, 0-32% at 39°C), and again tested both 1 and 5 dpa (to evaluate the effects of injury and regeneration, respectively) following anterior or posterior amputation, as well as uninjured controls (Fig. 1). We predicted that injury (1 dpa) would decrease thermal tolerance at both high and low temperature extremes, as a consequence of physical and physiological stress of injury, and that regeneration (5 dpa) would also decrease thermal tolerance, as a consequence of the costs of regeneration. Unexpectedly, we found that at 1 dpa, worms were more heat-tolerant than controls in both anteriorly (*P* < 0.001) and posteriorly injured (*P* < 0.01) worms (Fig. 2A). For cold tolerance, however, there was no effect of injury at 1 dpa (Fig. 2B). At 5 dpa, thermal tolerance of regenerated worms was mixed: only anteriorly regenerated worms were less heat-tolerant than controls (*P* < 0.05) (Fig. 2C), and there was again no difference in cold tolerance between groups (Fig. 2D). Overall, worms did not live as long at 5°C as they did at 39°C, with all worms being dead by roughly half the time under cold at both 1 dpa and 5 dpa as under heat. Interestingly, for control worms, the median time to 50% mortality was approximately the same (∼50 h) between the 1 and 5 dpa timepoints at 39°C, suggesting that food restriction over this duration does not affect heat tolerance. To confirm the surprising effect of injury-induced heat tolerance, we repeated this experiment focusing specifically on the 1 dpa timepoint and found again that anteriorly injured worms survived longer at 39°C (though posteriorly injured worms did not differ from controls) (Fig. S3A). There was again no difference between groups at 5°C (Fig. S3B).

**Figure 1.**
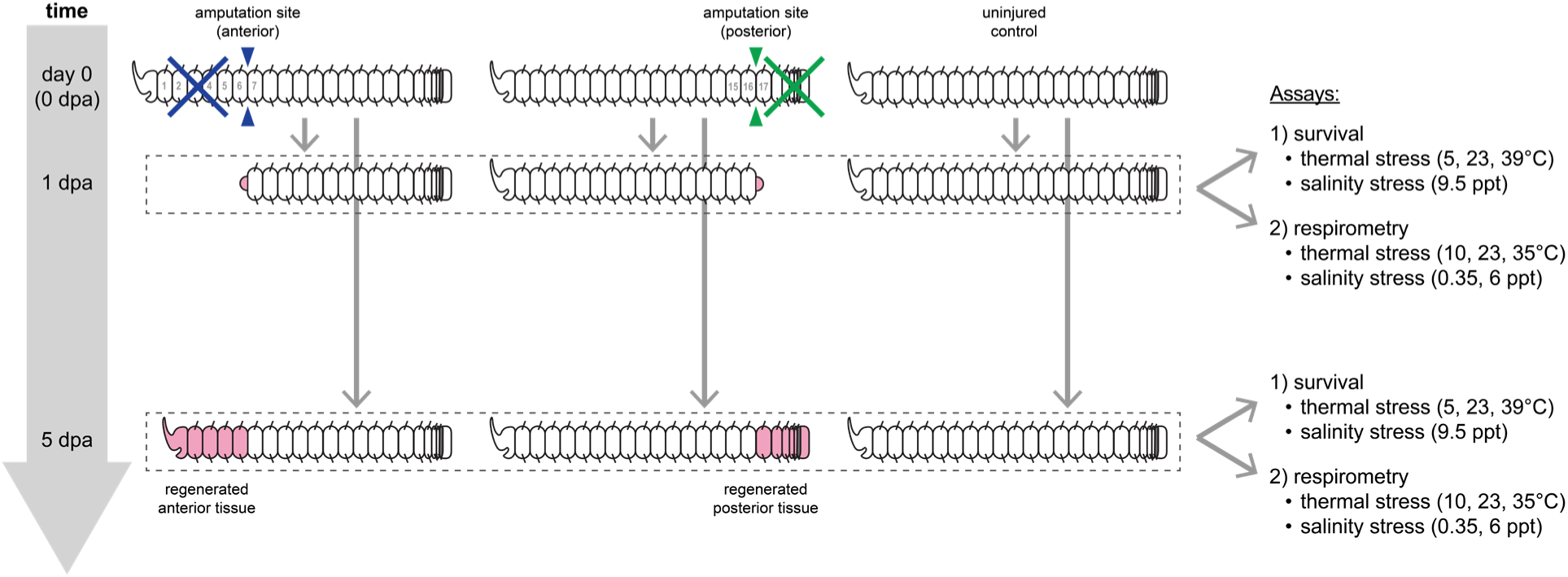
Experimental design for survival and respirometry experiments. Anterior is to the left in this and all subsequent figures.

**Figure 2.**
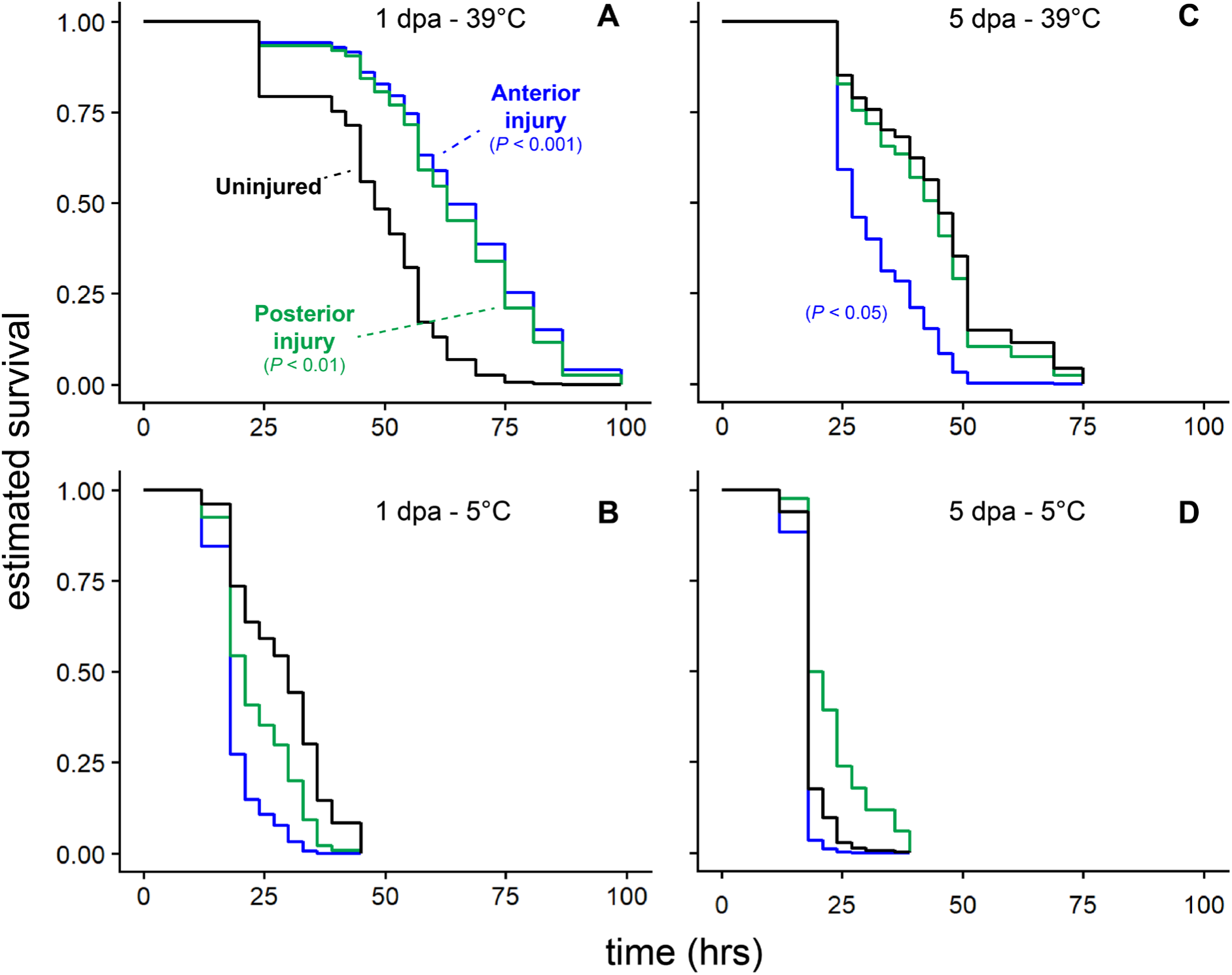
Estimated interval-censored survival of worms under continuous exposure to thermal stress. Worms were exposed to temperature (A,C: 39°C; B,D: 5°C) at 1 dpa (A,B) or 5 dpa (C,D). Color indicates injury condition (black = uninjured, blue = anterior amputation, green = posterior amputation). Each curve represents *n* = 15 worms. Significant differences between both injury conditions and control were detected in (A) (both *P* < 0.01) and between anteriorly regenerated worms and control in (C) (*P* < 0.05) via Kaplan-Meier survival analysis with Cox proportional hazards models.

Our finding that thermal tolerance was improved in recently injured worms led us to hypothesize that injury induces a broad stress response that improves tolerance to a variety of environmental stressors. To test this hypothesis, we performed a similar experiment to that described above but with salinity stress. *P. leidyi* is a freshwater species that is reared in the lab at ∼0.35 ppt. We performed an initial range determination experiment and found that, following a 48 hour exposure at a range of salinities, survival was substantially reduced (< 50%) at salinities at or above 9.5 ppt (data not shown). We then performed an experiment in which we exposed injured (1 dpa) or regenerated (5 dpa) worms to continuous 9.5 ppt salinity and monitored survival over time. The results of this experiment were inconsistent across repeated trials. In a first trial, there was no clear effect of injury on salinity tolerance at 1 dpa (Fig. S4A) but rapid death (within 4 hours) of all anteriorly regenerated worms at 5 dpa (*P* < 0.001), although uninjured and posteriorly regenerated worms all died shortly after (Fig. S4B). However, a second trial showed improved salinity tolerance in posteriorly (*P* < 0.001) and anteriorly injured worms, although the latter was not statistically significant (*P* ∼ 0.13) (Fig. S4C), and rapid death of all groups at 5 dpa, although posteriorly regenerated worms survived slightly longer than the rest (*P* < 0.001) (Fig. S4D). Additional follow-up trials (C.W. Rennolds, data not shown) were unable to consistently replicate any improved tolerance effect, perhaps pointing to the high sensitivity of *P. leidyi* to salinity conditions or the subtlety of the injury effect.

### Injury and regeneration affect oxygen consumption minimally

We speculated that injury- and regeneration-related changes in thermal tolerance may derive in part from altered metabolism. Prior work in various invertebrates has shown that aerobic respiration increases following injury or regeneration (Christensen et al., 2023; Collier, 1947; Hu et al., 2014; Needham, 1955, 1958; Stoner, 1970), and some theoretical frameworks implicate changes in standard metabolic rate (SMR) as underlying organismal performance under environmental stress (Schulte, 2015). Therefore, we investigated whether SMR is affected by injury and regeneration, and we predicted that SMR would be suppressed by injury and elevated by regeneration.

We used microplate respirometry to assess the rate of oxygen consumption, MO_2,_ in individuals at rest, a common proxy for SMR (Chabot et al., 2016; Tomlinson et al., 2018). Measurements were made on individual worms using a similar but sublethal set of experimental conditions as compared to our previous experiments (temperature: 10, 23, 35°C; salinity: 0.35, 6 ppt) (Fig. 1). For temperature, contrary to our predictions, there was no effect of injury (1 dpa) (Fig. 3A) or regeneration (5 dpa) (Fig. 3B) at either end of the body on MO_2_ under a range of temperatures. MO_2_ increased from colder to hotter temperatures, as expected, but injury condition did not significantly affect the rate of change in MO_2_. The interaction between timepoint and temperature was significant (ANCOVA *P* < 0.05), but no significant differences were detected from post-hoc comparisons between timepoints at any temperature. *Q*_10_, a measure of the sensitivity of biological processes to changes in environmental parameters including temperature (Mundim et al., 2020), was slightly higher in both injury groups versus controls (by ∼1.8 - 2.1) from 10 to 23°C at 1 dpa and in all groups between 1 and 5 dpa over the same temperature range, with controls increasing (by ∼1.1) and regenerated worms decreasing (by ∼1.6 – 1.9), but all other within- and between-timepoint *Q*_10_ comparisons differed minimally (by < 1) (Table S1). Estimated mass of worms used in these respirometry experiments was lower at 5 dpa than at 1 dpa for each of the three groups, presumably reflecting fasting, and importantly that reduction was comparable in magnitude for each group (uninjured, -5.4 µg; anteriorly regenerated, -4.7 µg; posteriorly regenerated, -6.7 µg) (all *P* < 0.05) (Fig. 3C). There is therefore no evidence that additional mass was consumed by regeneration-associated metabolic changes. Allometric scaling of MO_2_ was generally similar between injury conditions, temperatures, and recovery timepoints (Fig. S5).

**Figure 3.**
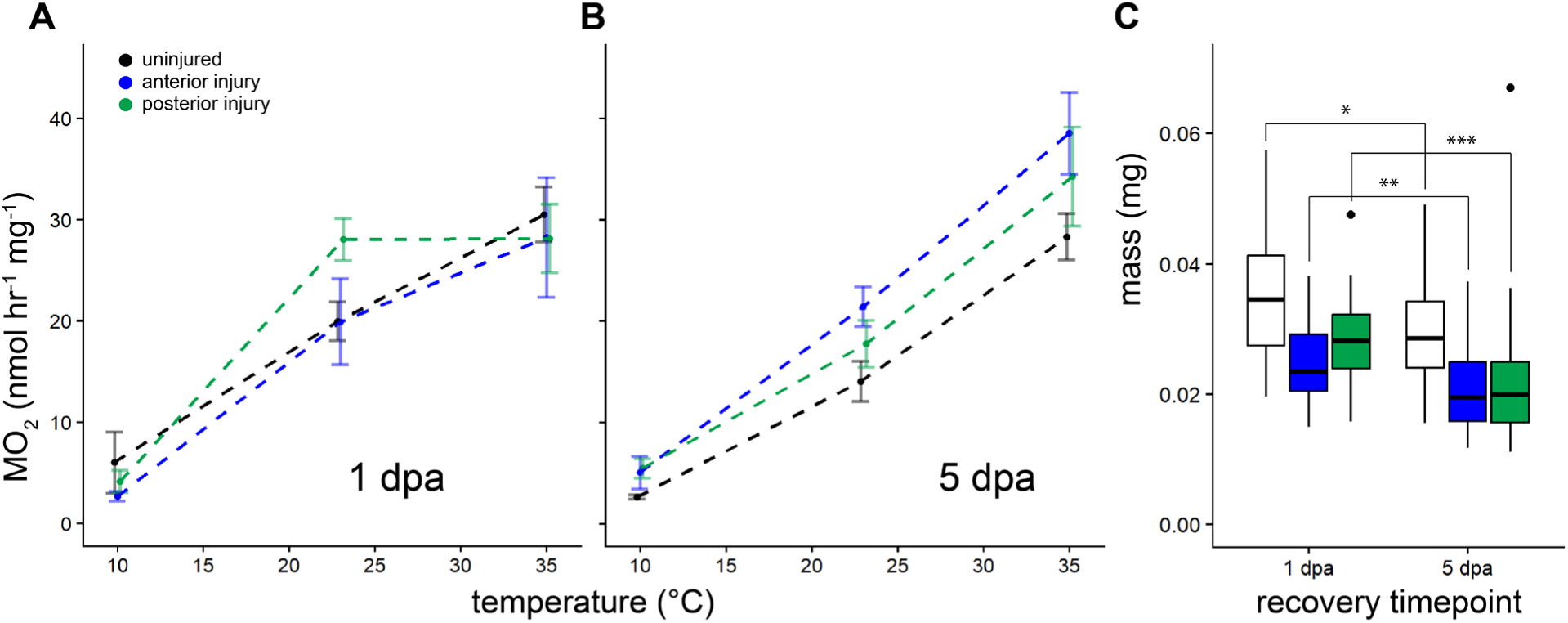
The effects of injury on physiological condition. (A-B) Oxygen consumption of worms under each injury condition across a range of temperatures was measured beginning at 1 dpa (A) or 5 dpa (B). Points at the same temperature are slightly offset for visibility. Error bars = s.e.m. (C) Worm mass by recovery timepoint and injury condition. * = *P* < 0.05, ** = *P* < 0.01, *** = *P* < 0.001. *n* = 18 worms pooled from three separate 6-worm replicate trials for all treatment combinations.

The effect of elevated salinity on respiration was somewhat more distinct, with MO2 (log transformed) being slightly affected by both salinity and post-injury timepoint. At 1 dpa, MO_2_ of control worms increased between 0 and 6 ppt, while MO_2_ decreased over the same interval in injured worms, but unlike temperature, the difference in MO_2_ at high and low salinity was only significantly different in posteriorly injured worms (*P* < 0.05) (Fig. S6A). In posteriorly injured worms, MO_2_ was significantly lower than in controls at 6 ppt only (*P* < 0.05). The difference in direction of change is reflected in the salinity *Q*_10_, with anteriorly and posteriorly injured worms having values of less than 1 (Table S2). Posterior regeneration (5 dpa) led to an increase in MO_2_ (*P* < 0.05), while other groups did not differ versus at 1 dpa (Fig. S6B). All groups had a positive *Q*_10_ at 5 dpa.

Ultimately, we found that the effects of injury and regeneration on metabolic rate under a range of temperature and salinity were slightly variable but minimal and with no obvious connection to patterns of survival.

### Shared elements in the transcriptional responses to injury and heat stress

The absence of a relationship between survival and SMR led us to hypothesize that injury may improve stress tolerance at least in part by inducing a general stress response at the molecular level. Induction of gene expression programs that are shared between injury and other stress responses could effectively “frontload” worms to better tolerate additional subsequent stressors by increasing the levels of active stress-protective factors, thus priming worms to exhibit a pronounced defensive response to further stress (e.g., Barshis et al., 2013; Collins et al., 2021). To assess whether transcriptional overlap exists between the responses to injury and to heat stress, whether their combination induces a more pronounced transcriptional response, and to identify candidate differentially expressed transcripts (DETs) that may confer cross-tolerance, we a conducted a 3’ TagSeq experiment (Fig. 4A). We extracted RNA and generated pooled cDNA libraries from worms injured anteriorly (A) or posteriorly (P) at 1 dpa, along with tissue controls to account for localized gene expression differences within each body region. Worms were subjected to nonlethal heat stress (35°C) or held at room temperature (23°C) within each injury group. For the remainder of this manuscript, we refer to the treatment combinations in this experiment using the following shorthand: A23, P23, CA23, CP23, A35, P35, CA35, and CP35 (Fig. 4B).

**Figure 4.**
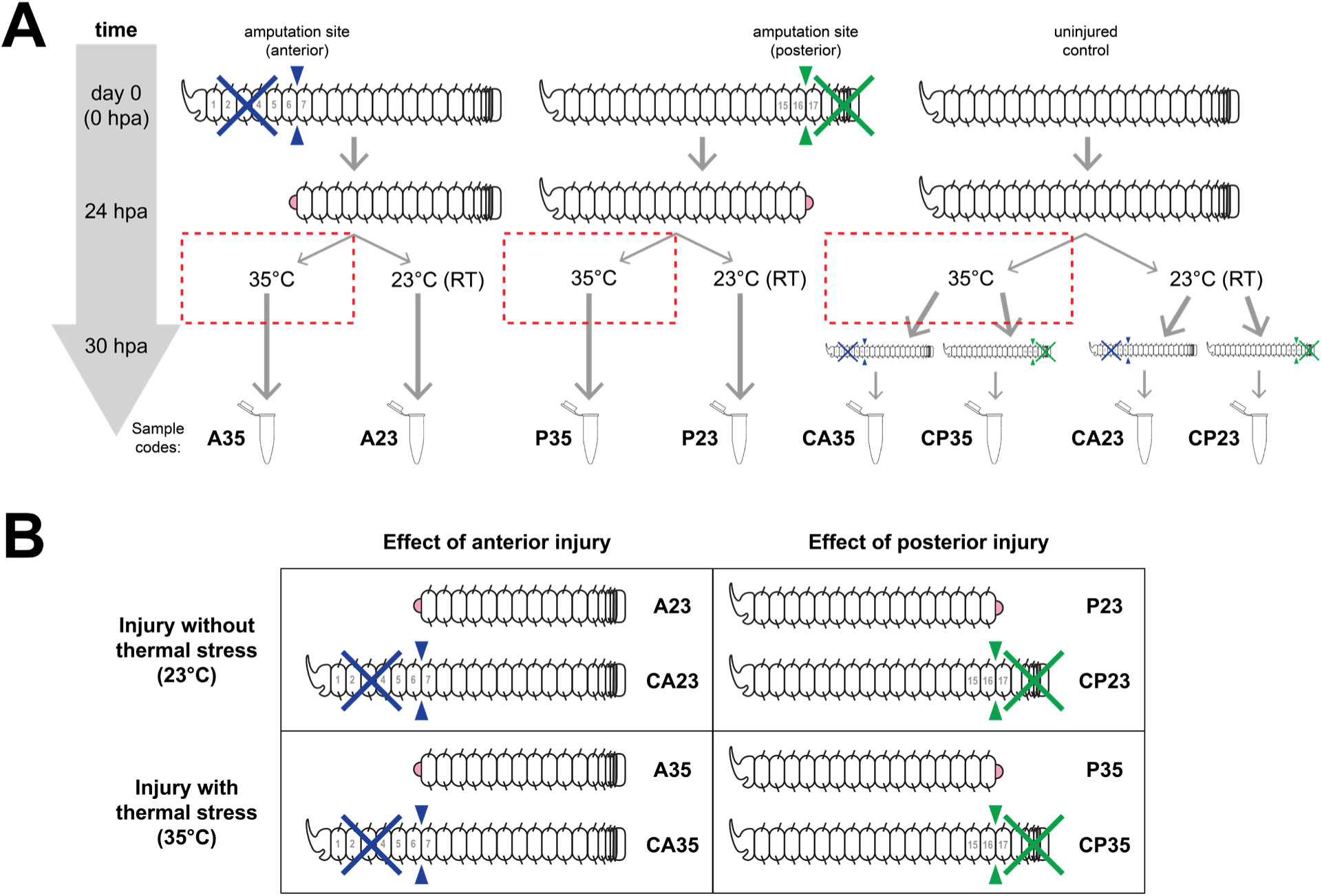
Experimental design of TagSeq experiment. (A) Injury and thermal stress scheme, with groups exposed to thermal stress and time period of exposure indicated by dashed red boxes. Thicker arrows indicate the passage of time between events according to the left axis, and thin arrows indicate the allocation of worms between events without respect to time. Each sample consisted of RNA pooled from 10-12 worms, and each treatment was replicated *N*=3 times. Codes used to refer to treatment groups are indicated along the bottom. (B) Matrix grouping each injury treatment with its corresponding tissue control under each thermal condition to indicate how main effects are determined.

TagSeq samples clustered fairly well by treatment group overall (Fig. S7). Mean-difference plots show that comparisons of interest are similarly distributed in transcript expression and that most DETs are relatively lowly expressed (Fig. S8), although the effect of posterior injury appears to disproportionately upregulate highly expressed transcripts (Fig. S8E,H), and unstressed anterior and posterior fragments (CA23 x CP23) are unsurprisingly distinguished by a relatively high proportion of transcripts with large differences in expression level (Fig. S8I). We identified a diverse assortment of both upregulated and downregulated DETs in pairwise comparisons of interest, with high representation of HSPs, metabolic enzymes, transporters, structural components of extracellular matrix, growth factors, ATPases, signaling molecules, and transcription factors (Tables S3-S13). A relatively large proportion of DETs distinguishing the anterior and posterior worm body (CA23 x CP23) were unannotated (Table S13), suggesting that much of the constitutive molecular profile that characterizes *P. leidyi* anatomy remains to be described.

The number of DETs detected in our comparisons of interest indicate striking differences in the magnitude of the responses to injury, heat stress, and their combination (Table S14). Injury alone at either location affected the expression of approximately 200 transcripts each (A23 x CA23: 227; P23 x CP23: 205), while heat stress alone roughly doubled that number (CA35 x CA23: 403; CP35 x CP23: 362). The addition of heat stress elicited more than twice as many DETs within anteriorly injured worms than within posteriorly injured worms (A35 x A23: 463; P35 x P23: 191), and a similar but more modest degree of expression distinguished the added effect of injury at each location within heat-stressed worms (A35 x CA35: 276; P35 x CP35: 96). However, the greatest magnitude of a transcriptional response in both body fragments was to the combination of injury and heat stress (A35 x CA23: 1650; P35 x CP23: 1296), suggesting a synergistic effect of injury on the subsequent response to a different source of physiological stress.

If injury-induced molecular frontloading confers broad stress tolerance, then this should be detectable as a molecular signature. Specifically, we would expect to see the same transcripts, or transcripts with similar function, differentially expressed by both injury and thermal stress separately. To evaluate this prediction, we identified overlapping DETs between selected combinations of treatment comparisons to identify candidate transcripts that may either confer a stress-protective effect or that may distinguish responses to stress between posterior and anterior tissue (Table S15). Lists of these overlapping DETs (those differentially expressed in the same direction) for those combinations of comparisons that are of greatest interest are presented in Tables S16-S19. Top DETs broadly include molecular chaperones, metabolic enzymes, extracellular matrix components, transcripts with immune function, transporters, and regulators of cell proliferation and differentiation. Heat stress alone consistently upregulated members of the HSP family, including *α-crystallin B*, *heat shock cognate 71*, and several co-chaperones.

Injury notably upregulated several histone proteins, suggestive of elevated DNA replication during wound healing and early regeneration, and downregulated a number of membrane transporters, suggesting shifts in metabolic activity and cellular homeostasis. We noted that heat stress elicited a larger shared response between tissue fragments (160 total DETs) than injury elicited between fragments (37 total DETs), suggesting that localized expression may be a relatively greater proportion of the transcriptomic response to injury than to heat stress, the latter of which is applied to the entire animal rather than to a specific body site.

The site of injury appears to have a considerable effect on the relative strength of the response to injury versus temperature in the composition DETs as well as their total number. An examination of the most variably expressed transcripts in each treatment group shows that these groups cluster primarily by tissue fragment (anterior or posterior), but the next level of clustering is contrasted between fragments: posterior worm fragments cluster primarily by temperature condition, while anterior fragments cluster primarily by injury condition (Fig. 5A). This suggests that the presence or absence of certain structures, such as the cerebral ganglion (brain), in each fragment may be critical for regulating the organismal response to environmental stress.

**Figure 5.**
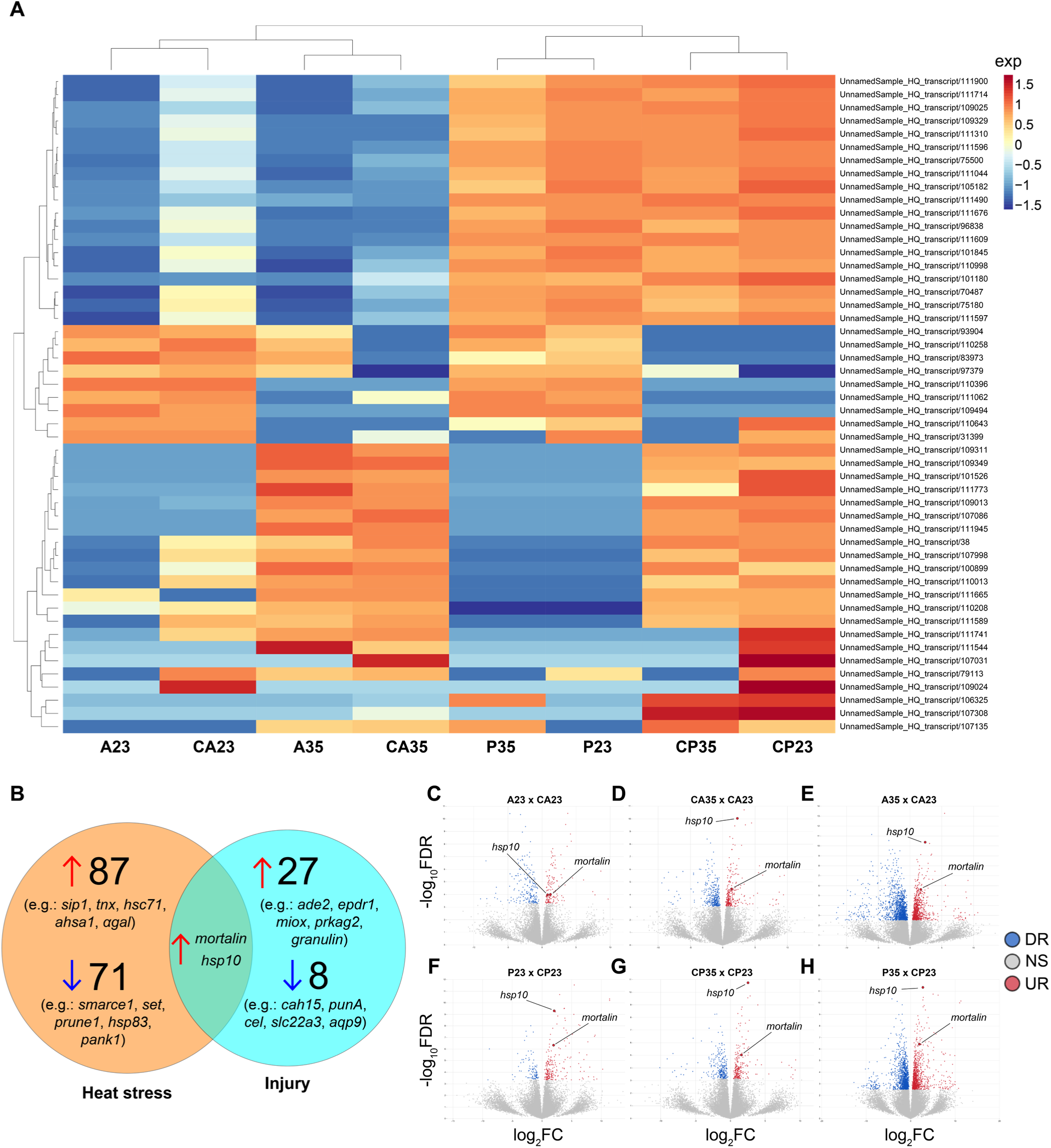
Output of differential expression analysis. (A) Heatmap of the top fifty most variable transcripts. Expression is determined as the variance of average log_2_CPM (counts per million). (B) Venn diagram showing the number of upregulated and downregulated transcripts induced by heat stress in both body fragments (CA35 x CA23 / CP35 x CP23), induced by injury at both sites of amputation (A23 x CA23 / P23 x CP23), and the two transcripts shared between them. (C-H) Volcano plots showing upregulated (UR), downregulated (DR), and nonsignificant (NS) transcripts in anterior-less (C-E) and posterior-less (F-H) fragments following injury (C,F), heat stress (D,G), and combined injury and heat stress (E,H). The transcripts *mortalin* and *hsp10* are indicated on each plot.

We then looked for overlap in the lists of shared DETs to identify common elements of the response to both injury and heat stress irrespective of body region (anterior or posterior). From this comparison, we identified only two transcripts, both coding for HSPs: mitochondrial *hsp10* (accession: 111384) and mitochondrial *hsp70*, often called *mortalin* (and referred to as such hereafter) (accession: 79043) (Fig. 5B). Both of these transcripts were upregulated by injury and heat stress alone and in combination in both body regions (Fig. 5C-H), with *hsp10* generally but variably more highly expressed than *mortalin* with the exception of in anteriorly injured worms at control temperature (Fig. 5C). We validated our TagSeq findings by performing qPCR for these two transcripts in separate groups of anteriorly injured and heat stressed worms (Fig. 6A; primer sequences in Table S20). We measured relative expression of these transcripts at different time points in each treatment condition to capture the potential window of upregulation induced by each treatment. We found that *mortalin* expression was elevated by both injury (*P* < 0.05) and heat stress (*P* < 0.01) at later time points (24 hpa and 6 h) (Fig. 6B), roughly corresponding to the TagSeq design, while *hsp10* was significantly elevated only by heat stress but at both 3 and 6 h (both *P* < 0.05) (Fig. 6C). However, mean *hsp10* expression was elevated at 24 hpa with high variance, suggesting that a significant effect could be identified with additional replication.

**Figure 6.**
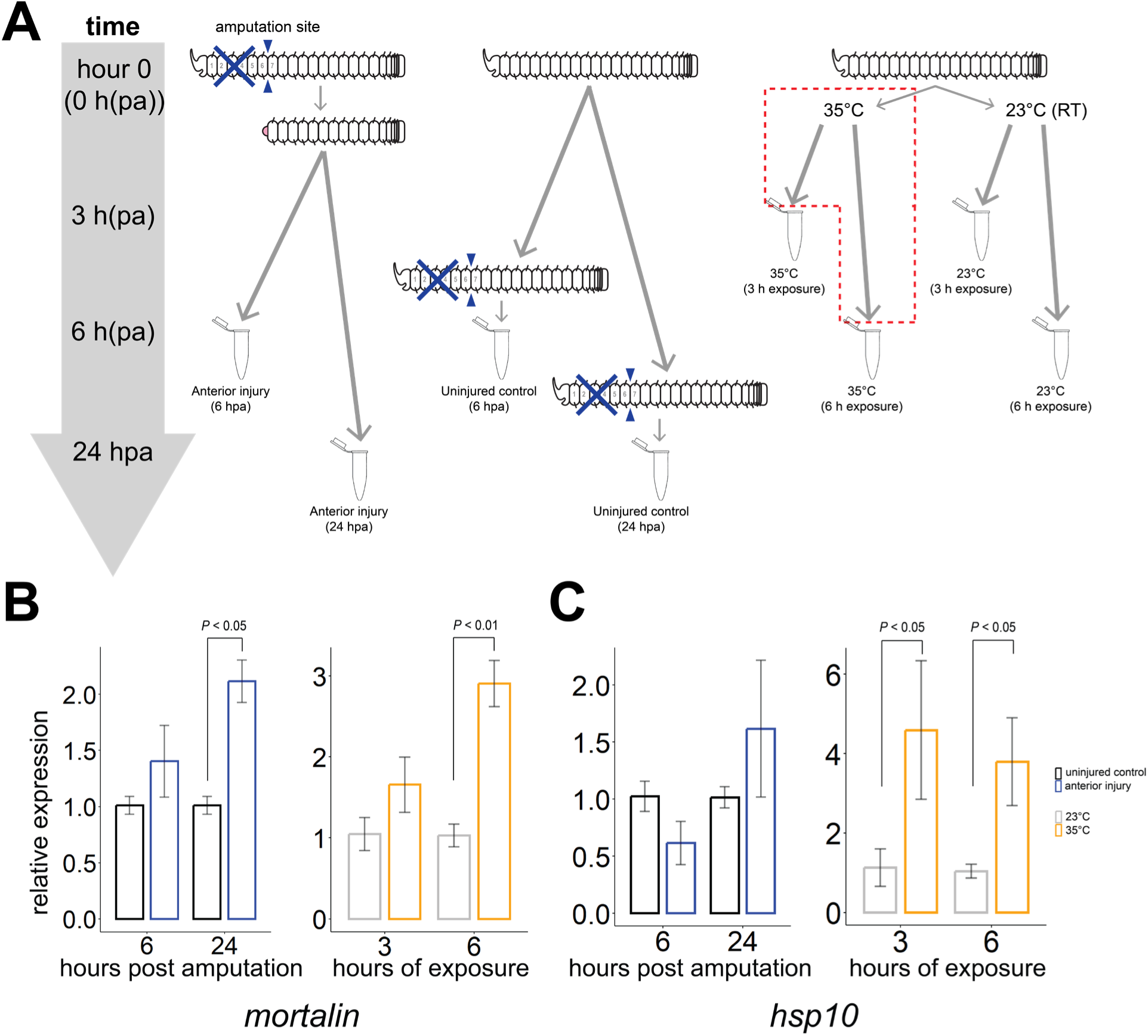
Expression of *mortalin* and *hsp10* following anterior amputation injury or thermal stress. (A) Experimental design, with groups exposed to thermal stress and time period of exposure indicated by a dashed red box. Thicker arrows indicate the passage of time between events according to the left axis, and thin arrows indicate the allocation of worms between events without respect to time. Each sample consisted of RNA pooled from 4-5 worms. Bottom panels show relative expression of target transcripts *mortalin* (B) and *hsp10* (C) in injured or thermally stressed worms versus control groups (set at 1) measured by qPCR. Expression was measured in *N=*5 replicates per treatment group (with one exception, see Methods). Comparisons showing statistically significant upregulation versus respective controls are indicated, with all others nonsignificant. Error bars = s.e.m.

### Mortalin inhibitor MKT-077 impairs both regeneration and heat tolerance

Since *hsp10* and *mortalin* were both upregulated by injury and heat stress and represent the entirety of the body-wide shared response to these stressors, we hypothesized that the induced expression of one or both of these transcripts by amputation may be at least partly responsible for improved heat tolerance in injured worms. We chose to focus on *mortalin*, since our qPCR data confirmed its induction by both injury and heat stress, and *mortalin* has known roles in development and thermoprotection in various organisms (Ben-Hamo et al., 2018; Conte et al., 2009; He et al., 2010).

The rhodacyanine dye MKT-077 is a compound that inhibits cancer cell proliferation by specifically binding to the nucleotide-binding domain of the mortalin protein (Li et al., 2013). We treated worms with 4 μM MKT-077 and determined whether it affected their survival and regeneration under control temperature and heat stress. Uninjured worms held at 23°C suffered no apparent effects from MKT-077 exposure, but all uninjured worms held at 37°C and treated with MKT-077 were dead by seven days (*P* < 0.01) (Fig. 7A). No worms treated with an equivalent percentage of DMSO solvent died during this period. A similar pattern applied to anteriorly injured worms, with the only death after seven days occurring in worms held at 37°C and treated with MKT-077 (*P* < 0.01) (Fig. 7B). However, unlike uninjured worms, several anteriorly injured worms were still alive in this group after seven days, perhaps reflecting a counteracting effect of injury-induced *mortalin* expression not fully inhibited by MKT-077 treatment. We also measured the length and fidelity of regenerated anterior tissue in surviving worms under the same treatment combinations. Regenerated tissue was shorter in MKT-077- treated worms versus DMSO controls when held at 23°C and at 37°C (both *P* < 0.0001), with one worm in the latter having no measurable regenerated tissue (Fig. 7C). Within MKT-077- treated worms, regenerated length was shorter in worms held at 37°C (*P* < 0.01). The fidelity and completeness of regeneration was clearly affected by MKT-077, as worms at both temperatures failed to form a proboscis, while those held at 37°C also failed to regenerate any other clear structural landmarks besides a partial mouth (Fig. 7D-G). Altogether, we show clearly that pharmacological inhibition of mortalin function reduces heat tolerance and severely impairs anterior regeneration.

**Figure 7.**
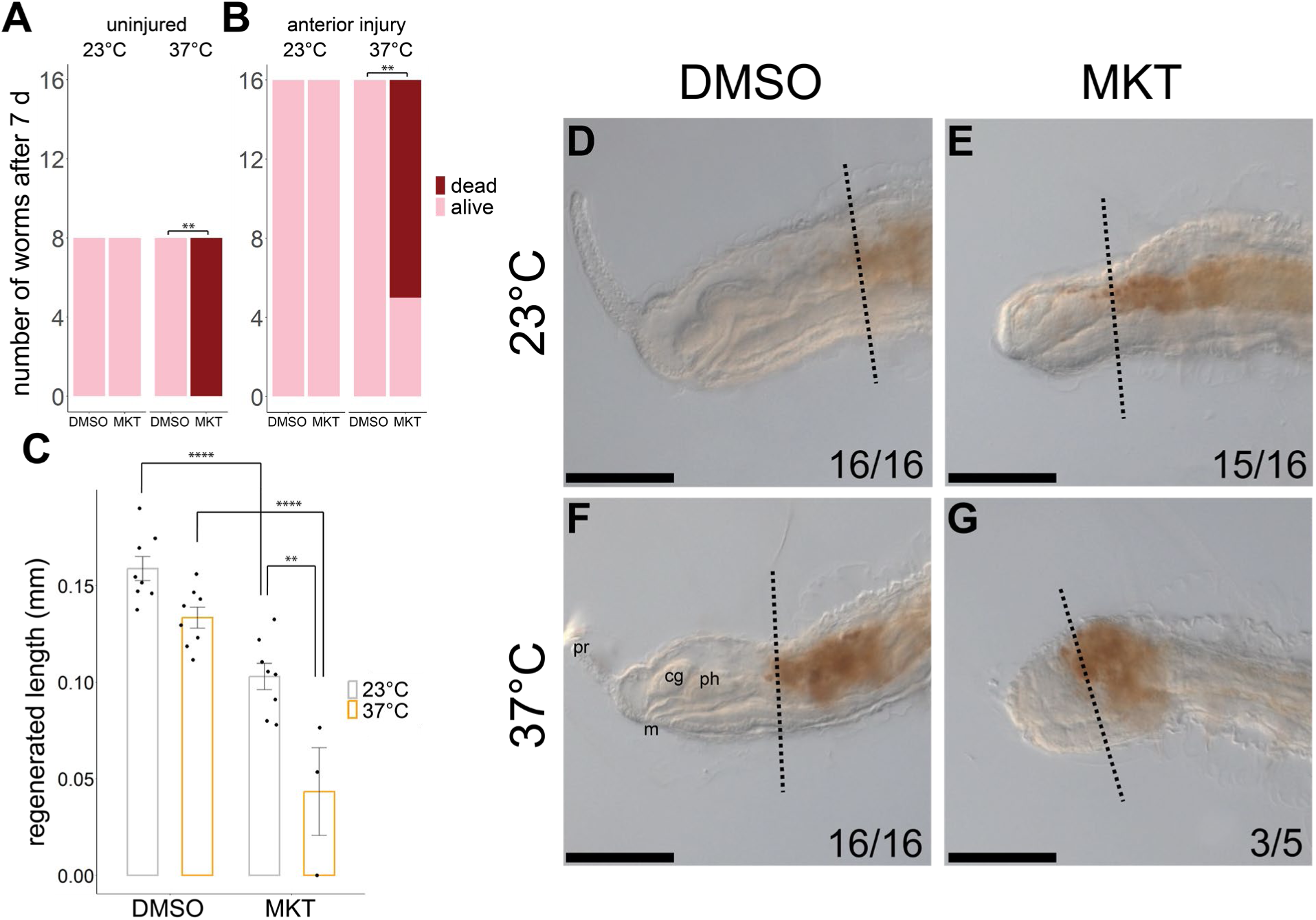
Effects of the mortalin inhibitor MKT-077 on survival and regeneration. (A-B) Number of surviving (pink) and dead (red) uninjured (A) and anteriorly amputated (B) worms after six days of exposure to either 0.04% DMSO or 4μM MKT-077 (MKT) under 23°C or 37°C. (C) Length of regenerated anterior tissue at 6 dpa under the same experimental conditions. Only statistically significant within-group pairwise comparisons are indicated. ** = *P* < 0.01, **** = *P* < 0.0001. (D-G) Representative images of regenerated anterior tissue (lateral view) at 6 dpa under the same experimental conditions. Vertical dashed lines represent the original amputation plane with regenerated tissue to its left. Scale bar = 100 µm. cg = cerebral ganglia, m = mouth, ph = pharynx, pr = proboscis.

## Discussion

In this study, we used a combination of techniques to characterize how injury and regeneration affect the ability of *P. leidyi* to tolerate environmental stress, a relevant measure of organismal function. We observed a mild improvement in survival time under heat (and, less consistently, salinity) stress in worms recently injured by amputation, whereas there was no clear effect of regeneration on worms’ tolerance to various forms of stress. Worms did not show improved tolerance to cold stress. We saw no evidence that injury or regeneration affect aerobic metabolic rate under a range of temperatures, leading us to investigate whether injury-induced heat tolerance has a basis in gene expression. TagSeq data show that injury induces a synergistic subsequent response to heat stress, and both injury and heat stress commonly induce the expression of two highly conserved HSP homologs, *hsp10* and *mortalin*. Pharmacological inhibition of *mortalin* led to a near-total loss of heat tolerance and considerably impaired regeneration of anterior tissue, indicating that this gene is a critical component of *P. leidyi*’s molecular response disparate forms of stress. We posit that injury induces the expression of *mortalin* and perhaps other genes that confer short-term resistance to broad forms of stress, which may be a significant aspect of *P. leidyi*’s ability to endure environmental disturbances.

Stress responses are theorized to manifest hierarchically, with systemic adjustments at the organismal level preceding responses at the cellular level (Kassahn et al., 2009; Pörtner, 2002). In this manner, potentially expensive changes at the cellular level, including the synthesis of stress-protective proteins, may be avoided when stress is brief or mild by first modifying higher-order physiological processes. With this perspective, we hypothesized that improved heat tolerance following injury could be explained by respiratory adjustments. The oxygen- and capacity-limited thermal tolerance (OCLTT) theory predicts that thermal tolerance is reduced when an animal’s demand for energy (ATP) exceeds its capacity to supply energy through aerobic respiration (Kassahn et al., 2009; Pörtner, 2010; Schulte, 2015; Sokolova, 2013; Sokolova et al., 2012). Conversely, anything that reduces energy demand at extreme levels of temperature or other stressors could hypothetically improve thermal tolerance. Cell proliferation is reduced body-wide for several days following amputation in *P. leidyi* (Zattara and Bely, 2013), which may reduce total metabolic expenditure, but we saw no evidence that injury depressed aerobic respiration. Cell proliferation may not contribute significantly to SMR in *P. leidyi* or its effects on SMR may be offset by increased ATP demand for other processes like the cellular stress response. Injury-induced changes in metabolic rate are inconsistent across animals: metabolic rate is briefly and minimally decreased after injury, followed by a prolonged increase during recovery, in various animals (Hu et al., 2014; Needham, 1955, 1958; Stoner, 1970), while oxygen uptake spikes just hours after amputation, followed by a sustained depression throughout regeneration, in the planarian *Schmidtea mediterranea*, although this depression is contingent on sexual mode and body fragment (Lewallen and Burggren, 2022). We also found no effect of regeneration on aerobic respiration, which contrasts with studies in *Tubifex*, members of the same annelid family as *P. leidyi*, and other annelids that exhibit an increased oxygen demand during regeneration (Anderson, 1956; Collier, 1947; Needham, 1958). Metabolic rate is significantly higher in regenerating versus intact brittlestars at elevated temperature, corresponding to increased regeneration rate but also higher mortality, but the authors did not detect a difference until the eighth week of regeneration (Christensen et al., 2023). In much more rapidly regenerating animal like *P. leidyi*, regeneration-induced metabolic shifts may be minimal or brief and thus difficult to detect.

The metabolic pathways involved during regeneration may also be primarily anaerobic and thus not detectable by respirometry. Early wound healing and regeneration initiation appears to be insensitive to oxygen in *Tubifex* (Anderson, 1956), and low oxygen may even promote healing in *Tubifex* worms and mammals, but not amphibians (Tsissios et al., 2026). Regeneration appears to be supplied predominantly through the anaerobic pentose phosphate pathway in *Xenopus* tadpoles (Patel et al., 2022) and possibly *Lumbriculus* annelids (Frank et al., 2025). In fireworms, lipid stores are depleted by regeneration, but the routes by which lipids are converted into ATP for supplying regeneration are unknown (Yáñez-Rivera and Méndez, 2014). Ultimately, the underlying mechanisms relating metabolism to phases of injury recovery remain obscure (Mack and Bely, 2025; Rennolds and Bely, 2023). For the present study, it is possible that OCLTT predictions may simply not apply to *P. leidyi* due in part to its small size and presumably diffusion-dependent oxygen uptake from the environment if aerobic respiration is even involved in its regeneration to any significant degree. Future work should assay anaerobic metabolism to understand how regeneration is fueled in *P. leidyi*, but we conclude that major changes in metabolic rate do not explain improved stress tolerance following injury.

We discovered two transcripts, *hsp10* and *mortalin*, consistently upregulated by injury and heat stress in our TagSeq analysis and found that inhibition of mortalin protein function with the compound MKT-077 led to drastically reduced heat tolerance and failure to regenerate properly. These findings suggest that *P. leidyi*’s primary response to environmental stress occurs at the cellular level rather than through systemic adjustments and that the heat shock response is a candidate cause of the injury-induced heat tolerance that we observed. HSPs are upregulated following injury in various animals (Husmann et al., 2014; Li et al., 2014; Matranga et al., 2000; Pinsino et al., 2007; Sánchez Navarro et al., 2009; Stewart et al., 2017; Sveen et al., 2019; Vazzana et al., 2015; Wenger et al., 2014), including the annelid *Lumbriculus variegatus* (Tellez- Garcia et al., 2021), and some, such as *hsp60*, which was commonly upregulated in our own treatment comparisons, function in regulating the regeneration process (Li et al., 2014; Ma et al., 2022; Makino et al., 2005; Patruno et al., 2001; Pei et al., 2016; Sun et al., 2025). While they were initially named due to being elicited by heat stress, various stressors may induce the expression of HSPs, which are broadly involved in maintaining cellular integrity by stabilizing proteins or facilitating their turnover (Kultz, 2020b; Richter et al., 2010; Sørensen et al., 2003). Members of this protein family are found throughout our DET lists, but only one other study to our knowledge, in anemones, has investigated their expression during simultaneous injury and other stress (e.g., temperature) (La Corte et al., 2025). Mortalin is specifically known to regulate developmental processes in a variety of animal systems and is particularly relevant for cancer research due to the link between mortalin dysfunction and uncontrolled cell proliferation (Ben-Hamo et al., 2018; Conte et al., 2009; Esfahanian et al., 2023; Kaul et al., 2007; Xu et al., 2020). Its contribution to heat tolerance appears mixed: while a mortalin-like protein from the fluke *Schistosoma japonicum* confers heat tolerance and is inducible (He et al., 2010), *mortalin* is unresponsive to heat and found at high levels of constitutive expression in planarians, specifically in neoblasts (Conte et al., 2009). Although components of the heat shock response are conserved and widespread in animals due to their critical role in supporting protein function, the heat shock family has likely diversified and expanded to support many different aspects of organismal function in various animal lineages, including in development (Tomanek, 2010). Some annelid HSPs have been described in multiple species, but their specific functions are not yet well understood (de la Fuente and Novo, 2022). Their role in regeneration merits further investigation.

Mechanical injury and environmental stress are expected to induce an ancestral, conserved molecular program. Responses to different forms of stress at higher levels of biological organization are likely to be highly diverse, as they involve great complexity dependent on the physiological traits and evolutionary history of different species (Sulmon et al., 2015). However, at lower levels of biological organization, various forms of stress disrupt function in many similar ways, including damage to cell membranes and extracellular matrix or the generation of radical oxygen species (Cossins and Prosser, 1978; Hazel and Williams, 1990; Somero, 2020). At the cellular level, the damage caused by amputation is likely to be comparable to that caused by heat stress; therefore, the molecular response to these distinct stressors ought to overlap considerably. The cellular stress response is common and likely plays a role in wound healing and regeneration in diverse animals (Feder and Hofmann, 1999; Kultz, 2020b; Somero, 2020), but how it may differ including in its interactions with the developmental programs regulating regeneration, thereby contributing to variation in regeneration ability, will require more work in a greater number of species.

The specific mechanism by which injury improves heat tolerance may be by inducing the expression of components of the cellular stress response, such as *mortalin*, that have broad function in repairing or preventing cellular damage (Fig. 8). In animals, the phenomenon by which one stressor upregulates the expression of protective proteins that improve tolerance to subsequent stress is referred to as molecular or transcriptional “frontloading”, particularly in relation to phenotypic plasticity in constitutive gene expression, whereas a comparable phenomenon may be called “priming” in plants. When one stressor, such as heat, improves tolerance to a subsequent stressor, such as hypoxia, the organism is said to exhibit “cross-tolerance”, which has been shown to occur for various stressors (Gotcha et al., 2018; Gunderson et al., 2016; Sinclair et al., 2013; Todgham and Stillman, 2013). The process by which small amounts of stress or damage improve organismal condition by inducing repair pathways is also known as “hormesis” (Calabrese and Baldwin, 1998; Costantini et al., 2010). Amputation injury may thus frontload the expression of *mortalin* and other genes, including *hsp10* and perhaps other genes with spatial or temporal specificity in *P. leidyi*, conferring enough cross-tolerance to moderately extend survival under heat stress. Additional work is required to verify the contribution of frontloading to the phenomenon we observed, including measures of protein to connect with gene expression data or overexpressing *mortalin* and other genes without injury to determine if they improve heat tolerance in a similar manner as injury.

**Figure 8.**
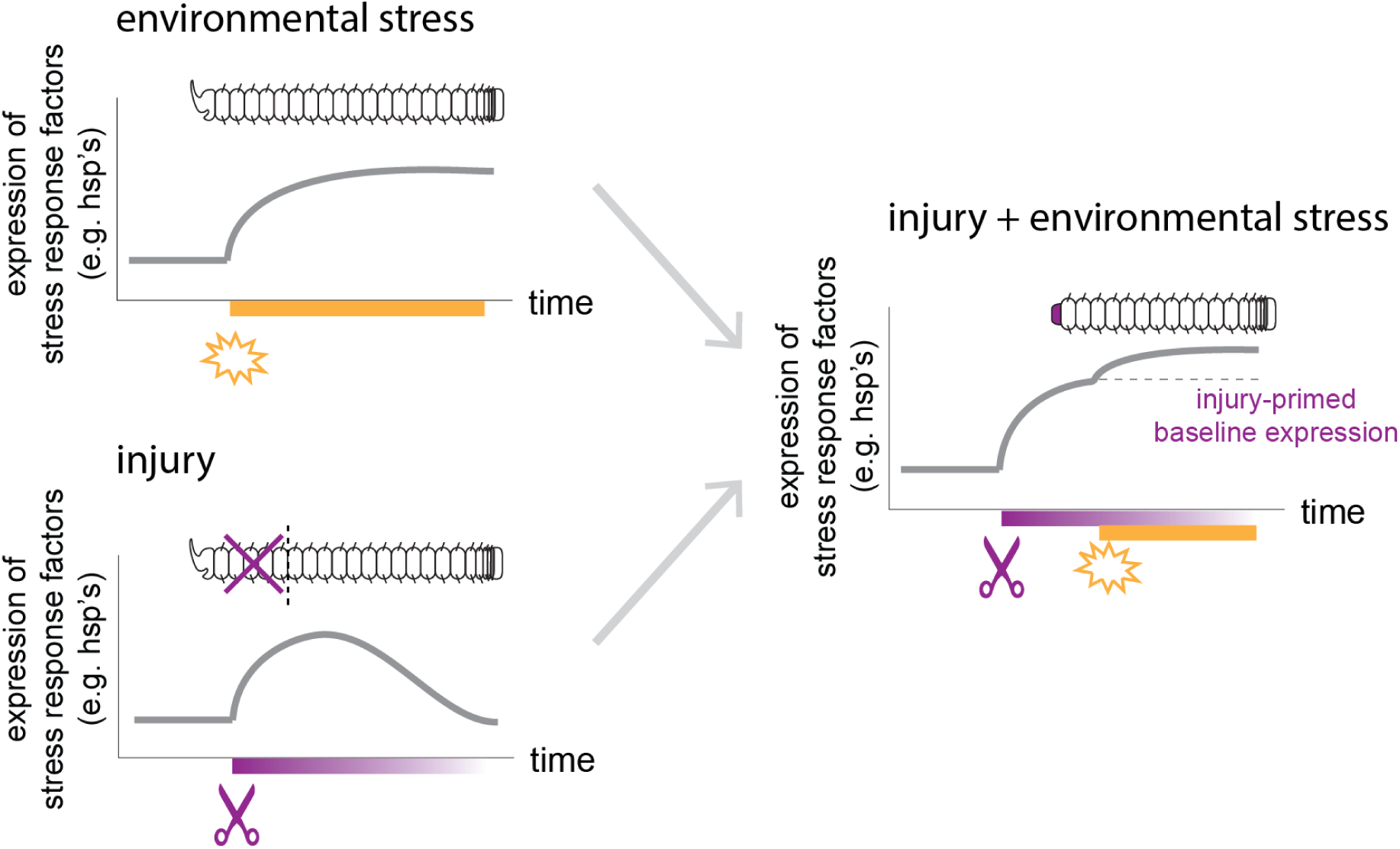
Working model of injury-induced priming of stress tolerance. Upon exposure to an acute stress, expression of the stress response program starts from the minimum baseline. Sublethal injury induces the expression of stress-responsive genes. If exposed to a separate acute stressor during the period of injury-induced expression, levels of shared stress response factors begin at an elevated baseline, allowing the maximum stress response to be reached more quickly and improving tolerance. The magnitude and sensitivity of the overall stress response may also be synergistically multiplied.

We found that injury and heat stress increase overall gene expression to a degree greater than that predicted by summing the individual effects of each stressor. Combinations of stressors often elicit transcriptomic and physiological responses of a magnitude not predictable from those elicited separately (Crain et al., 2008; DeBiasse and Kelly, 2016; Gunderson et al., 2016; Holmstrup et al., 2010; Sokolova, 2021; Todgham and Stillman, 2013). In addition to frontloading expression of specific protective genes, injury may also prime worms to exhibit a much larger overall response to additional near-future stress through an unknown mechanism, perhaps involving *mortalin* regulation directly. Studies of multiple-stressor interactions are often focused on stressors associated with either climate change (e.g., temperature, hypoxia, pH) or anthropogenic pollution (e.g., heavy metals, endocrine disruptors), whilst interactions between injury and any of these factors receive far less research attention. *P. leidyi* and other annelids inhabit freshwater environments like small streams where multiple forms of stress including physical damage may affect them simultaneously, such as during storms or droughts. Studying how injury and regeneration affect organismal function under a range of environmental conditions is therefore important for understanding the significance of these processes in the ecology of annelids and other regenerating animals.

Despite the potential overlap between the transcriptional responses to injury and heat stress, we found that overall gene expression was largely distinct between the two. Of the 89 transcripts commonly upregulated by heat stress, only two (2.25%) were also upregulated by injury. However, many of the distinct transcripts encode for proteins with similar broad predicted functions, including in the regulation of cellular metabolism, transcription, and DNA repair, suggesting that different molecular players are enlisted in similar suites of processes. We also find many similarities between our results and those from other studies of gene expression during regeneration in annelids. An earlier RNA-seq study of *P. leidyi* reported the upregulation of *serine/threonine kinase*, a regulator of apoptosis, *multifunctional protein ADE2*, and *glutamate dehydrogenase* during anterior regeneration (del Olmo et al., 2022), all of which were upregulated following injury (the latter in posteriorly injured worms only) in our study. Those authors reported over a thousand differentially expressed genes during anterior regeneration, with most of those being downregulated, in contrast to our study which reports far fewer DETs (97 upregulated and 130 downregulated). However, our study used worms at 1 dpa, while the prior study used worms at 3 dpa, indicating that the peak of the transcriptional program of regeneration occurs later in the regeneration process than what we investigated. RNA-seq studies in other annelid species report many other shared components of the regeneration program at various stages (Pare et al., 2023; Ribeiro et al., 2019; Tellez-Garcia et al., 2021), which we will not elaborate on here. Much remains to be done to characterize the specific functions of most of these genes in regeneration and elucidate the molecular pathways common to regeneration within annelids and across animals broadly.

Although we did not see a clear and consistent difference in the physiological response of *P. leidyi* to stress between injury and regeneration of anterior versus posterior tissues, we expect that these pose different potential costs for the animal. Posterior regeneration occurs entirely through epimorphosis, while anterior regeneration of more than four segments, as in our study, also includes morphallactic remodeling of intact tissue (Zattara and Bely, 2011). Whether these two types of regeneration differ in their resource costs or the metabolic pathways to fuel them are unknown but could affect organismal function in ways we failed to measure or detect and has been suggested to impact metabolism in other annelids (Needham, 1958). Anterior tissues contain the brain, an organ that contributes a large amount of energy consumption in various animals (Hand and Hardewig, 1996) and may be important in providing neural or endocrine inputs controlling growth and regeneration of posterior tissues, as shown in the annelid *Platynereis dumerilii* (Alvarez-Campos et al., 2023; Planques et al., 2019). In other annelids, posterior structures contribute to oxygen uptake, although the species for which this is documented are much larger and less able to meet their oxygen demand through diffusion across the external body surface (Glasby et al., 2021; Julian et al., 1996). These and other anatomical features may have different effects on the ability of the remaining fragment to function following their removal, particularly in feeding, but understanding these effects requires further investigation in *P. leidyi*. In contrast, the consequences of head removal are obvious for the many annelids that cannot regenerate anteriorly (Zattara and Bely, 2016).

Our TagSeq results, meanwhile, suggest that the early injury response and regeneration program differs considerably between anterior and posterior regeneration, with only 37 of a combined 396 DETs (9.34%) shared between the two in either direction. The transcriptomic profiles between early anterior and posterior regeneration are also quite distinct in syllid annelids, with posterior regeneration more closely resembling patterns of gene expression in normal posterior growth than they resemble anterior regeneration (Ribeiro et al., 2019). Posterior regeneration morphologically resembles normal growth in many ways in *P. leidyi* as well, and the timing and nature of developmental events during regeneration vary with injury location (Zattara and Bely, 2011, 2013). The presence or absence of structures like the brain may also have an effect on the subsequent response to heat, as in the nematode *Caenorhabditis elegans*, which coordinates the heat shock response body-wide through anterior neural inputs, although cells also respond autonomously (Prahlad et al., 2008). Of 535 combined DETs induced by heat within injured *P. leidyi* worms, 119 (22.24%) were shared, and 160 (26.45%) of 605 DETs induced by heat alone were shared between uninjured body fragments, indicating a major difference in the composition or magnitude of expression of the heat shock response in different regions of the worm body. A general difference in the relative response to heat versus injury is also apparent in how body fragments cluster primarily by different experimental factors in the TagSeq data of most variably expressed transcripts. Further investigation of how specific regions of the body respond to physiological challenges may provide some insight for understanding the relative consequences of losing anterior versus posterior structures and subsequently regenerating them, or not, as in anterior regeneration-incompetent naids like *Paranais* and *Chaetogaster* spp. (Bely and Sikes, 2010). Such insights could help identify candidate physiological or molecular targets of selection driving the modification or loss of anterior regeneration ability in annelids.

Our overall findings and multilevel approach allow us to make several inferences regarding the effects of injury and regeneration on organismal function. Although many animals, including annelids like *P. leidyi*, can regenerate an indefinite number of times, it is not well understood what the short- or long-term impacts of regeneration are in most species. We found that *P. leidyi* is physiologically resilient during and shortly after regeneration and may even benefit in a small way under harsh conditions by being physically damaged, perhaps incidentally but possibly through an adaptive feature of the organismal response to stress. *P. leidyi*’s metabolic rate is either unaltered by regeneration or only during a window of time we did not examine, suggesting that this species meets its energy requirements without major shifts in metabolic expenditure. Clitellate annelids like *P. leidyi* store energy in the chloragogue, an organ that polychaete annelids lack and that may supply regeneration without leading to major shifts in physiological condition (Pandian, 2019). Animals without sufficient energy stores may suffer greater functional effects during regeneration if they are unable to compensate with feeding, a hypothesis that must be tested with further comparative studies. *P. leidyi*’s infaunal mode of life may make any given body part temporarily expendable, but other animals with more complex or specialized structures may not survive their loss even if their regeneration is technically possible. The functional costs of regenerating such structures may also vary based on environmental conditions and an animal’s physiological traits, including the existence of trade-offs with other processes, and genetic toolkit. While many factors will determine how injury and recovery impact animals (Mack and Bely, 2025; Rennolds and Bely, 2023), these impacts may ultimately shape the fitness value of regeneration itself such that it is modified or lost entirely over evolutionary time scales (Goss, 1969). Patterns of regeneration across animals may therefore reflect complex relationships between their intrinsic properties themselves and involving the conditions of the environments in which they evolved.

## Acknowledgements

We would like to thank the following people for their help on various aspects of this project: Spencer Brodsky, for spearheading early attempts at respirometry experiments in naids; Todd Kana, Christopher Rowe, and members of the Nye Lab at Stony Brook University for their guidance and hands-on assistance with respirometry; Jordan Mohondro for helping with survival experiments and worm cultures; past and present members of the Bely Lab for their feedback on experiments and the manuscript; members of the Haag Lab at the University of Maryland for assistance with RNA extractions; Amy Beaven at the University of Maryland Genomics Core for training and assistance with qPCR and RNA analysis; Duygu Özpolat, Nathan Kenny, and Jordi Solana for sharing the *P. leidyi* IsoSeq reference transcriptome prior to its publication; the Genomic Sequencing and Analysis Facility at the University of Texas for all library preparation and sequencing; and Tsetso Bachvaroff and Mark Newton for bioinformatics pipeline assistance. We also thank the Institute of Marine and Environmental Technology in Baltimore, MD for the use of their computing resources to perform sequencing analyses.

