## Supplemental Materials for "Injury improves tolerance to acute heat stress through induction of heat shock protein expression in the annelid *Pristina leidyi*"

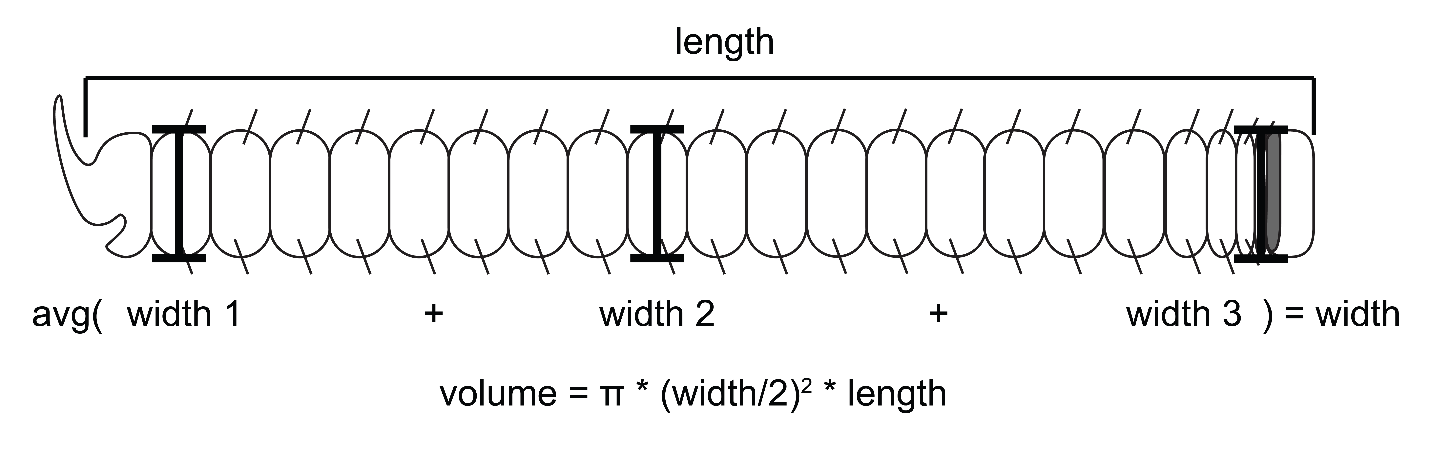
**Figure S1. Measurement and calculation of worm physical dimensions.** Cartoon is oriented with the anterior (head) to the left and posterior to the right.

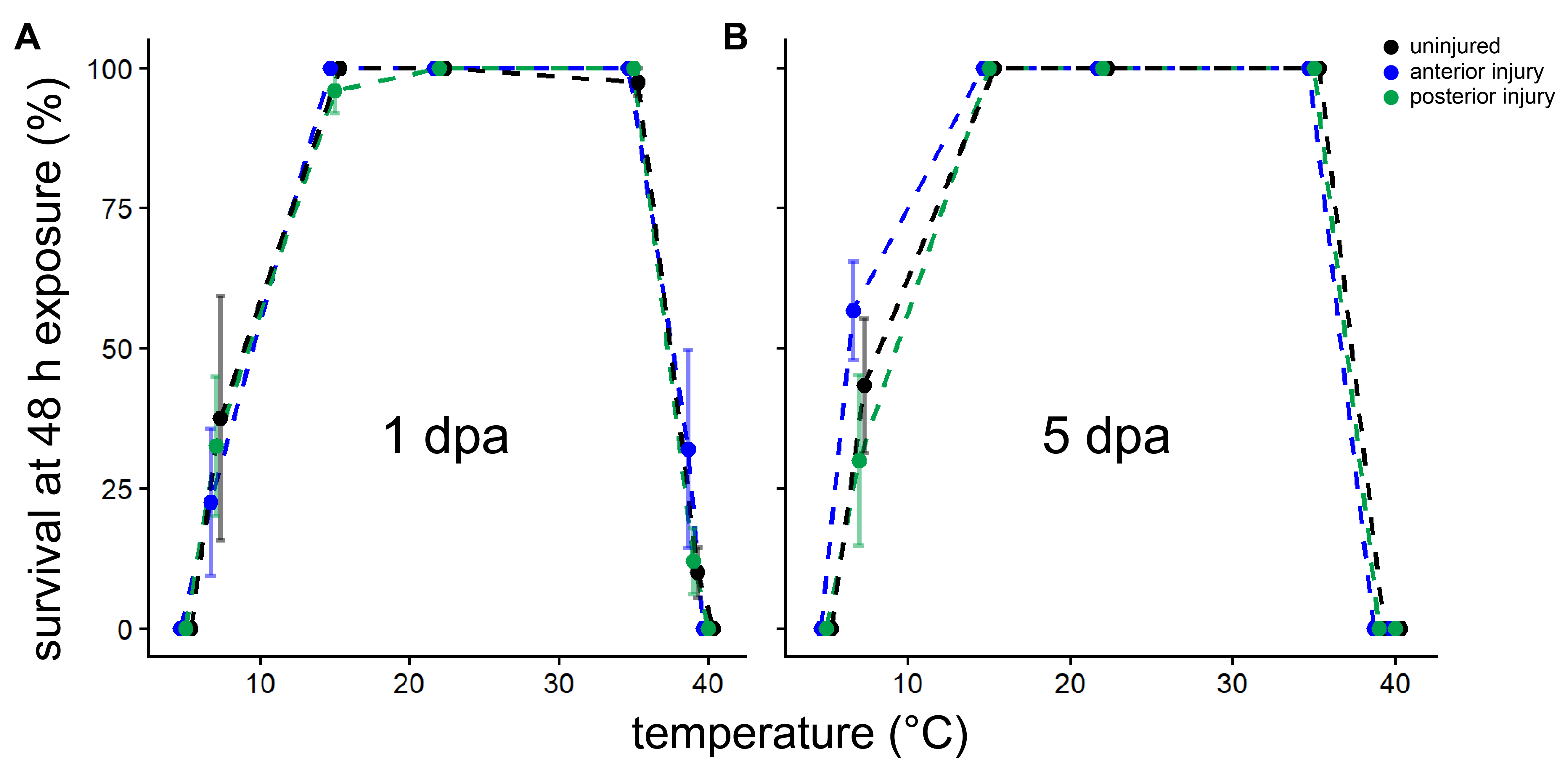

**Figure S2. *Pristina leidyi* acute thermal tolerance range.** Survival percentage of worms following 48-hour exposure to temperature (5, 7, 15, 22, 35, 39, and 40°C) at 1 dpa (i.e., thermal exposure 1-3 dpa) (A) and 5 dpa (i.e., thermal exposure 5-7 dpa) (B). *N*=5 for each data point. Error bars = s.e.m.

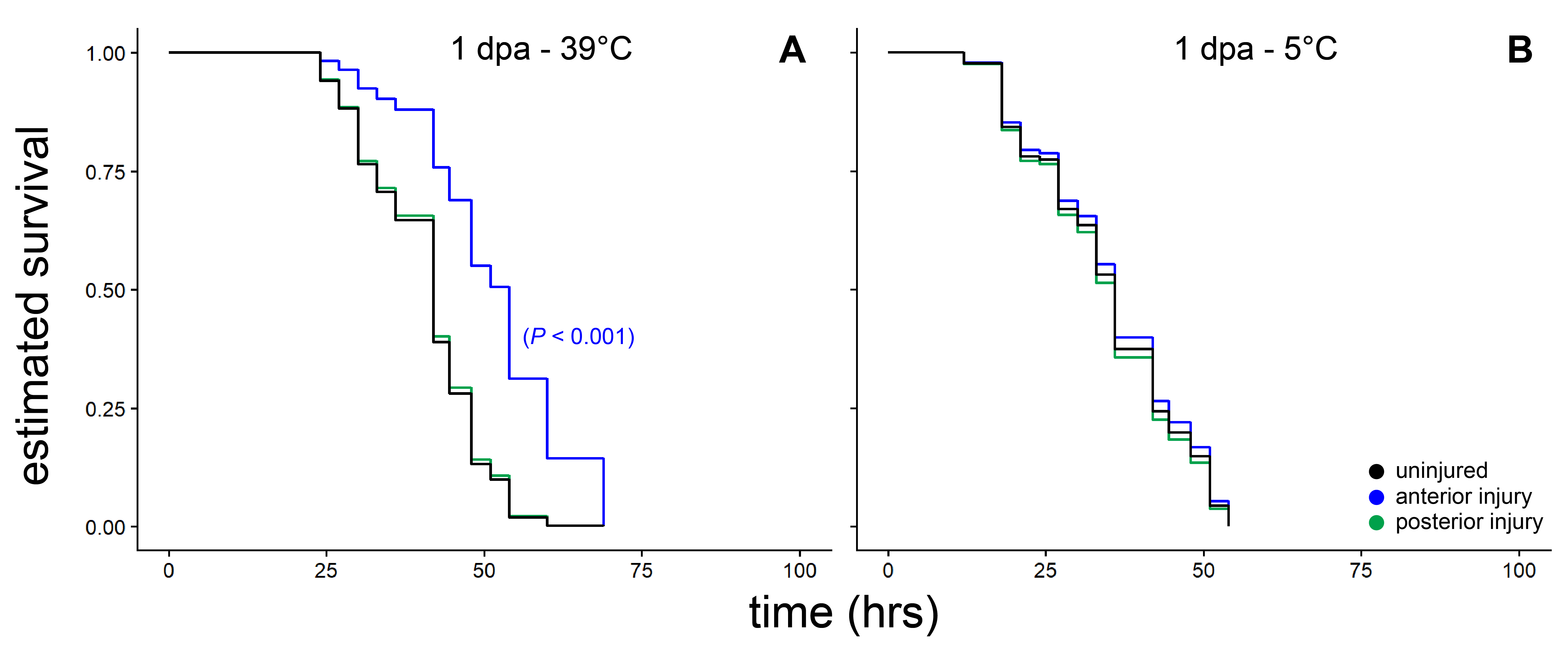

**Figure S3. Estimated interval-censored survival of worms under continuous exposure to thermal stress, second replicate.** Worms were exposed to 39°C (A) or 5°C (B) at 1 dpa. Each curve represents *n* = 15 worms. A significant difference between anteriorly injured and control worms was detected at 39°C (*P* < 0.001) via Kaplan-Meier survival analysis with Cox proportional hazards models.

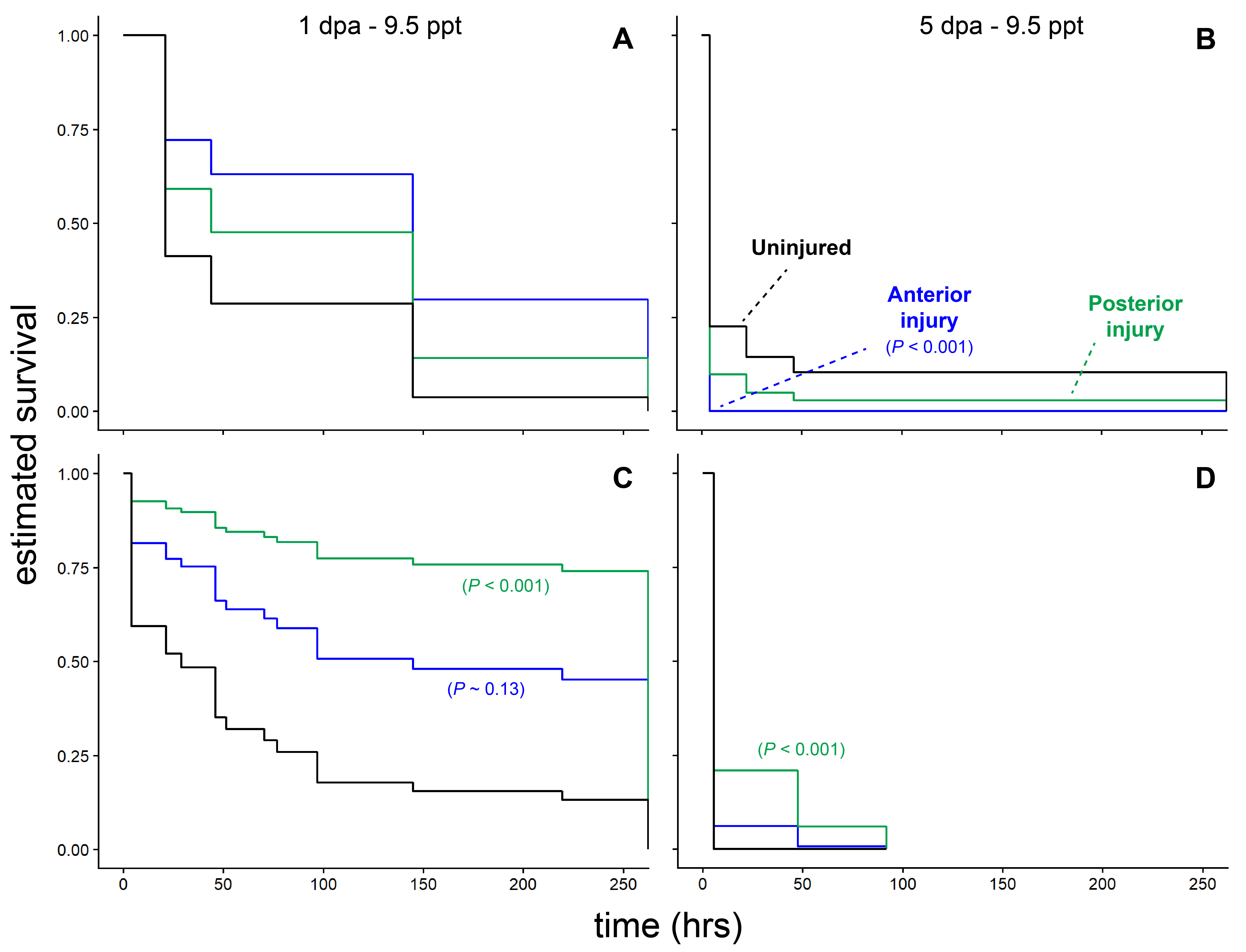

**Figure S4. Estimated interval-censored survival of worms under continuous exposure to salinity stress.** Worms were exposed to high salinity (9.5 ppt) at 1 dpa (A,C) or 5 dpa (B,D) across two replicate experiments (A-B & C-D). Color indicates injury condition (black = uninjured, blue = anterior amputation, green = posterior amputation). Each curve represents *n* = 15 worms. Significant differences indicated between injury conditions and controls were detected via Kaplan-Meier survival analysis with Cox proportional hazards models.

**Table S1.** Calculated Q_10_ of each injury condition at each recovery time point across experimental temperature intervals (from data shown in Figure 3)

|  | **Q_10_: 10→23°C** | | **Q_10_: 23→35°C** | | **Q_10_: 10→35°C** | |
| --- | --- | --- | --- | --- | --- | --- |
| **Injury treatment** | *1 dpa* | *5 dpa* | *1 dpa* | *5 dpa* | *1 dpa* | *5 dpa* |
| Uninjured | 2.5182 | 3.6051 | 1.4236 | 1.7938 | 1.9151 | 2.5787 |
| Anterior | 4.6397 | 3.0413 | 1.3379 | 1.6311 | 2.5542 | 2.2552 |
| Posterior | 4.3385 | 2.4742 | 1.0015 | 1.7312 | 2.1465 | 2.0844 |

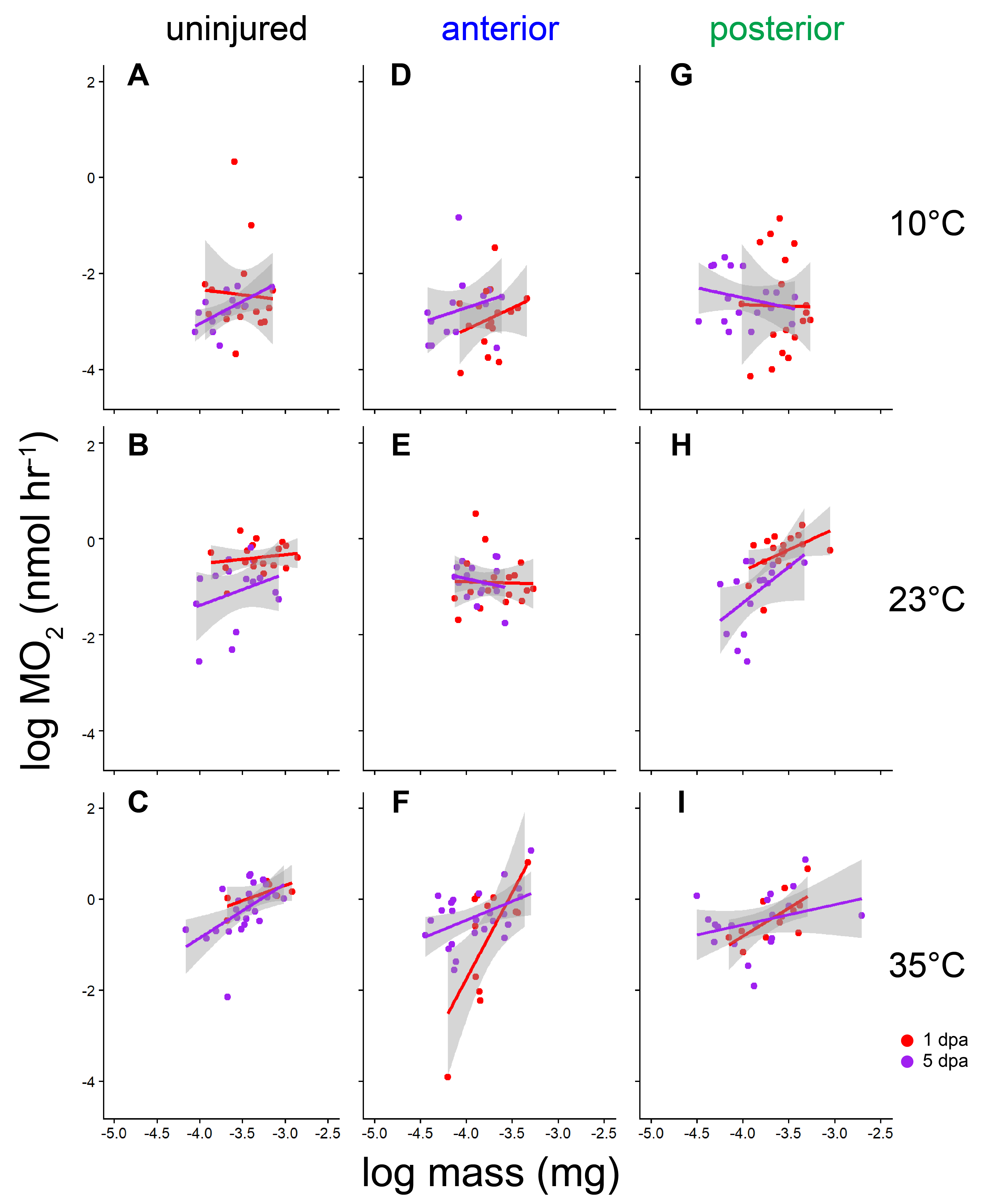

**Figure S5. Allometric scaling of aerobic respiration.** Log of oxygen consumption rate (MO_2_) as a function of estimated worm mass between 1 dpa (red) and 5 dpa (purple) uninjured (A-C), anteriorly injured (D-F), and posteriorly injured (G-I) worms at 10 (A,D,G), 23 (B,E,H), and 35°C (C,F,I). Fitted lines are constructed using a linear method. Shading represents 95% confidence intervals.

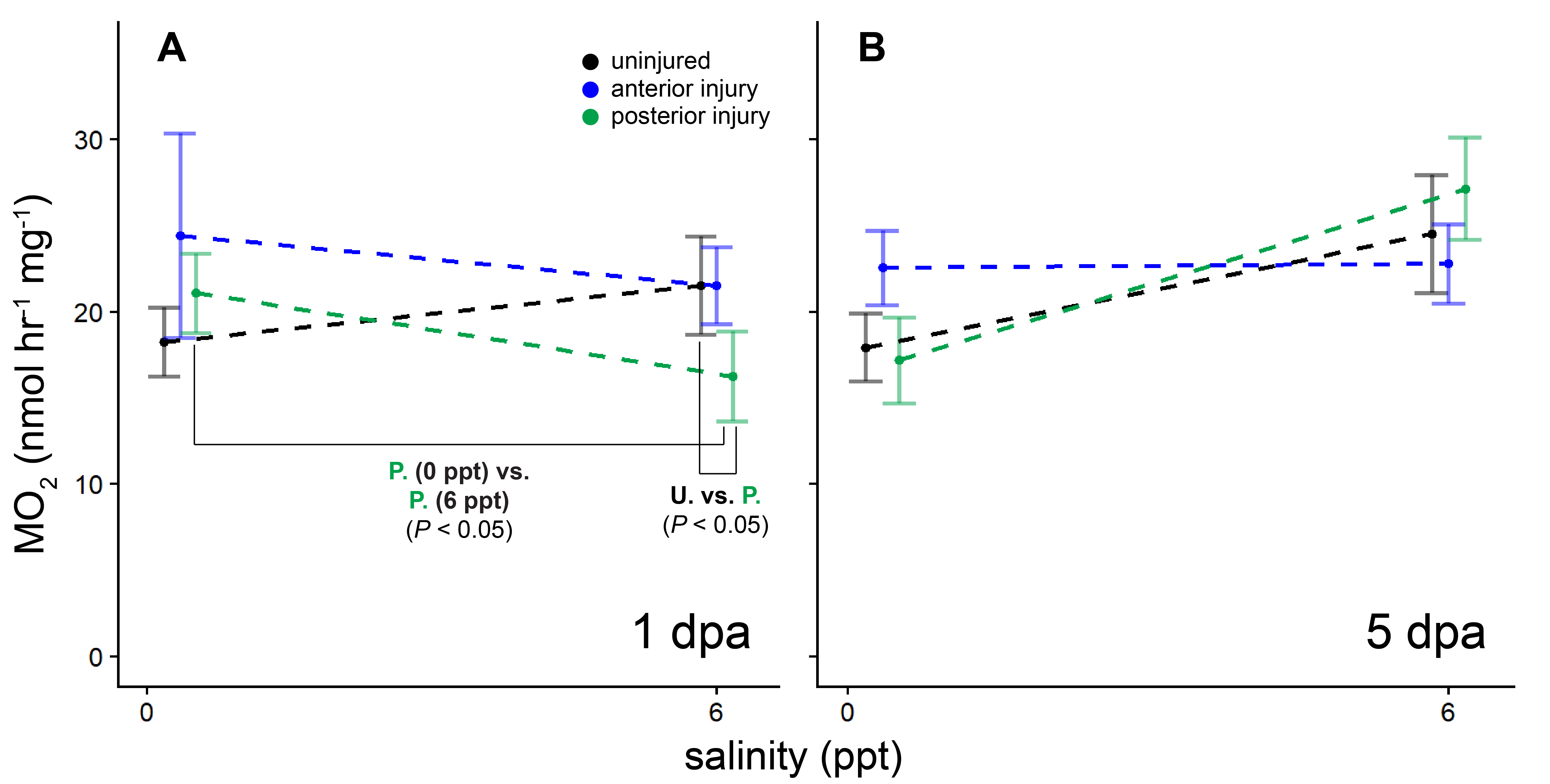

**Figure S6. The effects of injury on the metabolic response to elevated salinity.** Plots of salinity versus mass-specific (raw) MO_2_ for 1 dpa (A) and 5 dpa (B) worms under each injury condition. Each point represents *n* = 17-18 worms pooled from three separate 6-worm trials (with two outliers removed, see Methods). Points at the same salinity level are slightly offset for visibility. Significant Tukey’s-adjusted post-hoc comparisons of log-transformed MO_2_ between experimental groups are indicated, excepting those between 1 and 5 dpa for simplicity (see Results for details). Error bars = s.e.m.

**Table S2.** Calculated Q_10_ of each injury condition at each recovery time point across experimental salinity range (from data shown in Figure S5)

|  | **Q_10_: 0.35→6 ppt** | |
| --- | --- | --- |
| **Injury treatment** | *1 dpa* | *5 dpa* |
| Uninjured | 1.3374 | 1.7387 |
| Anterior | 0.7989 | 1.019 |
| Posterior | 0.6303 | 2.2457 |

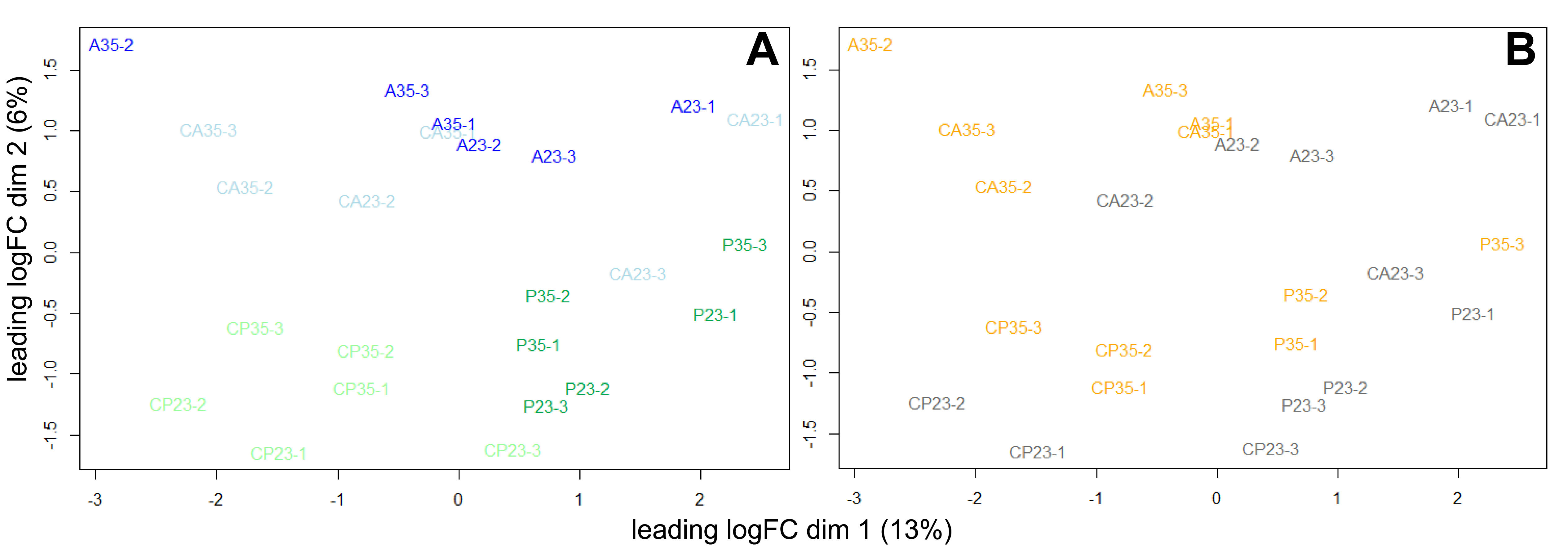

**Figure S7. Multidimensional scaling plot of TagSeq samples.** Individual library replicates of experimental groups are plotted based on the top two explanatory dimensions as Euclidean distances between transcript expression profiles. Color coding corresponds to (A) injury treatment (light blue = CA, blue = A, light green = CP, green = P) or (B) temperature treatment (gray = 23°C, orange = 35°C).

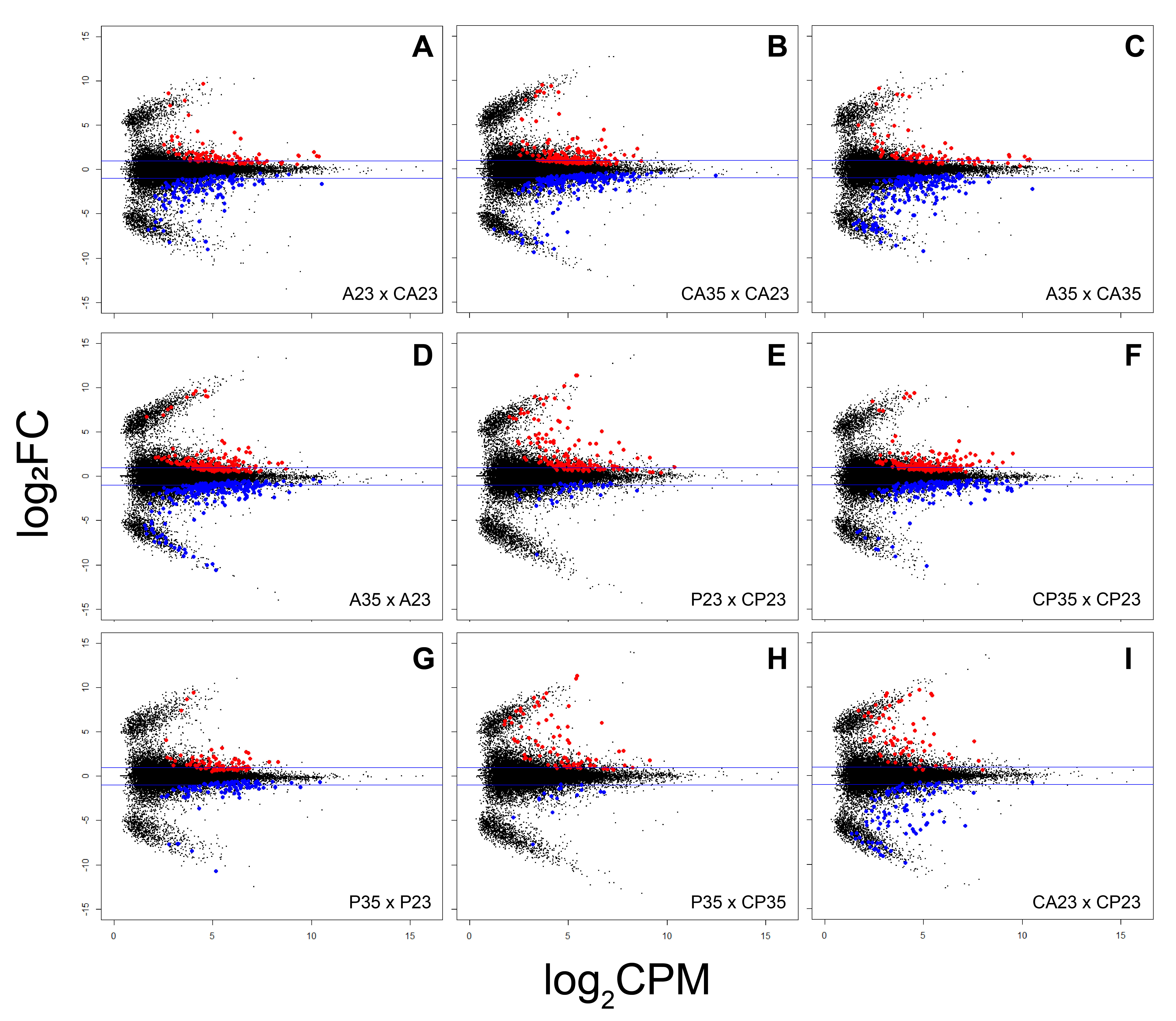

**Figure S8. Mean-difference plots of selected TagSeq comparisons.** Plots show all filtered transcripts (dots) including those differentially expressed (red = upregulated, blue = downregulated; FDR < 0.05) for the pairwise comparisons: A23 x CA23 (A), CA35 x CA23 (B), A35 x CA35 (C), A35 x A23 (D), P23 x CP23 (E), CP35 x CP23 (F), P35 x P23 (G), P35 x CP35 (H), CA23 x CP23 (I). Horizontal blue lines mark ±1 log_2_FC (fold change). CPM = counts per million.

**Table S3.** Selected DE transcripts (FDR < 0.001) associated with anterior injury (A23 x CA23)

| **Transcript ID** | **UniProtKB / Pfam ID** | **Protein** | **E-value** | **Protein coordinates** | **logFC** | **logCPM** | **F** | **P value** | **FDR** |
| --- | --- | --- | --- | --- | --- | --- | --- | --- | --- |
| 98623 |  |  |  |  | -3.13556 | 5.209282 | 162.3944 | 1.83E-10 | 6.28E-06 |
| 98411 | CAH15_MOUSE | Carbonic anhydrase 15 | 1.57E-48 | 219-1187[+] | -2.39307 | 4.529742 | 147.0926 | 4.09E-10 | 7.02E-06 |
| 104706 |  |  |  | 164-487[+] | -4.68026 | 5.598169 | 130.0723 | 1.17E-09 | 1.34E-05 |
| 96233 | PUR6_CHICK | Multifunctional protein ADE2 | 0 | 157-1425[+] | 3.49399 | 6.421227 | 121.0594 | 2.01E-09 | 1.73E-05 |
| 91287 | PUNA_GEOSE | Purine nucleoside phosphorylase 1 | 1.97E-17 | 2-457[+] | -2.83877 | 5.401997 | 103.6322 | 6.57E-09 | 4.52E-05 |
| 111433 | CAT8_MOUSE | Cathepsin 8 | 5.18E-11 | 83-439[+] | -1.65828 | 7.125899 | 97.53961 | 1.05E-08 | 6.01E-05 |
| 91191 | FBP1_STRPU | Fibropellin-1 | 8.25E-37 | 36-1361[+] | -8.21738 | 2.801388 | 125.2985 | 2.18E-08 | 0.000107 |
| 97432 | PUR6_CHICK | Multifunctional protein ADE2 | 0 | 132-1400[+] | 4.19349 | 6.087589 | 82.98669 | 3.86E-08 | 0.000164 |
| 88942 | CKS1_MOUSE | Cyclin-dependent kinases regulatory subunit 1 | 1.42E-29 |  | 1.390159 | 5.273971 | 80.97685 | 4.3E-08 | 0.000164 |
| 65316 | PDE9A_HUMAN | High affinity cGMP-specific 3',5'-cyclic phosphodiesterase 9A | 2.03E-163 | 256-1431[+] | 1.592953 | 4.368625 | 78.40992 | 5.46E-08 | 0.000167 |
| 102787 |  |  |  | 252-1001[+] | 3.772886 | 2.930086 | 78.03995 | 5.66E-08 | 0.000167 |
| 108369 | GRN_DICDI | Granulin | 7.03E-06 |  | 1.475071 | 10.38169 | 77.70205 | 5.84E-08 | 0.000167 |
| 102110 | VPP4_HUMAN | V-type proton ATPase 116 kDa subunit a isoform 4 | 9.56E-108 | 1-1392[+] | -1.76917 | 6.456291 | 74.18268 | 8.23E-08 | 0.000217 |
| 70960 | OSTA_LEUER | Organic solute transporter subunit alpha | 1.63E-13 | 3-1184[+] | -1.83469 | 3.912291 | 70.68732 | 1.17E-07 | 0.000234 |
| 75833 | IIGP5_HUMAN | Interferon-inducible GTPase 5 | 4.42E-53 | 127-1329[+] | -3.14794 | 3.09102 | 70.49707 | 1.19E-07 | 0.000234 |
| 109959 | GRN_DICDI | Granulin | 2.90E-05 |  | 1.567083 | 9.363465 | 70.37634 | 1.21E-07 | 0.000234 |
| 73659 | VPP1_RAT | V-type proton ATPase 116 kDa subunit a1 | 0 | 1-2499[+] | -2.14321 | 5.262253 | 70.34297 | 1.21E-07 | 0.000234 |
| 88157 | ADH_SULAC | NAD-dependent alcohol dehydrogenase | 3.72E-16 | 231-1547[+] | 1.410683 | 5.137089 | 70.25655 | 1.22E-07 | 0.000234 |
| 106647 | EPDR1_HUMAN | Mammalian ependymin-related protein 1 | 5.47E-05 | 99-803[+] | 1.91607 | 4.670539 | 68.93944 | 1.4E-07 | 0.000254 |
| 105510 | DUT_RAT | Deoxyuridine 5'-triphosphate nucleotidohydrolase | 1.57E-72 | 1-606[+] | 1.665216 | 4.365238 | 67.32697 | 1.67E-07 | 0.000286 |
| 8722 | GVIN1_MOUSE | Interferon-induced very large GTPase 1 | 0 | 50-5014[+] | -2.5136 | 4.119865 | 65.60689 | 2.01E-07 | 0.000328 |
| 99366 | STAR5_BOVIN | StAR-related lipid transfer protein 5 | 2.68E-49 | 183-842[+] | 1.447534 | 6.249263 | 56.13154 | 6.04E-07 | 0.000937 |
| 109771 | GRN_DICDI | Granulin | 8.24E-06 |  | 1.5513 | 10.27211 | 55.57083 | 6.57E-07 | 0.000937 |
| 79013 | MIOX_DANRE | Inositol oxygenase | 1.16E-113 | 142-996[+] | 1.714183 | 7.339379 | 54.95092 | 6.99E-07 | 0.000937 |
| 111225 |  |  |  |  | 1.954403 | 10.12021 | 55.02424 | 7.25E-07 | 0.000937 |
| 110698 | PF04103.18 | CD20-like family | 2.70E-07 | 100-858[+] | -2.78856 | 4.671946 | 54.7067 | 7.3E-07 | 0.000937 |
| 109797 | GRN_DICDI | Granulin | 2.26E-05 |  | 1.615888 | 9.313346 | 54.15159 | 7.82E-07 | 0.000937 |
| 79730 | SGMA_DICDI | Sphingomyelin phosphodiesterase A | 1.01E-69 | 114-2189[+] | -1.83826 | 4.416679 | 54.03349 | 7.85E-07 | 0.000937 |
| 110851 | IF5A_DROME | Eukaryotic translation initiation factor 5A | 1.85E-59 | 45-548[+] | 0.62119 | 9.270789 | 53.83296 | 8.06E-07 | 0.000937 |
| 22453 | AAKG2_MOUSE | 5'-AMP-activated protein kinase subunit gamma-2 | 4.26E-70 | 109-3705[+] | 2.072012 | 4.981187 | 53.5303 | 8.37E-07 | 0.000937 |
| 110968 |  |  |  |  | 1.851908 | 4.514405 | 53.45625 | 8.45E-07 | 0.000937 |
| 77962 | NQO1_CAVPO | NAD(P)H dehydrogenase [quinone] 1 | 2.31E-14 | 3-1367[+] | -2.86985 | 5.284109 | 53.28075 | 9.01E-07 | 0.000967 |

**Table S4.** Selected DE transcripts (FDR < 0.001) associated with thermal stress in posterior fragments (CA35 x CA23)

| **Transcript ID** | **UniProtKB / Pfam ID** | **Protein** | **E-value** | **Protein coordinates** | **logFC** | **logCPM** | **F** | **P value** | **FDR** |
| --- | --- | --- | --- | --- | --- | --- | --- | --- | --- |
| 109908 | CRYAB_MACFA | Alpha-crystallin B chain | 1.34E-15 | 129-629[+] | 3.314269 | 6.691767 | 174.7168 | 1E-10 | 2.74E-06 |
| 111384 | CH10_ORYLA | 10 kDa heat shock protein | 4.96E-43 | 116-421[+] | 2.27355 | 8.513635 | 165.1441 | 1.59E-10 | 2.74E-06 |
| 99284 | POSTN_MOUSE | Periostin | 1.81E-17 | 1-903[+] | 1.876368 | 5.974099 | 141.4008 | 5.62E-10 | 6.44E-06 |
| 84988 | TENX_HUMAN | Tenascin-X | 5.16E-29 | 98-2230[+] | 4.477983 | 6.805893 | 124.6449 | 1.67E-09 | 1.15E-05 |
| 77307 | HSP7C_SAGOE | Heat shock cognate 71 kDa protein | 0 | 201-2153[+] | 3.516845 | 5.483423 | 122.2006 | 1.84E-09 | 1.15E-05 |
| 108040 | U2AF1_BOVIN | Splicing factor U2AF 35 kDa subunit | 9.49E-11 | 64-732[+] | 1.116211 | 6.861008 | 120.618 | 2E-09 | 1.15E-05 |
| 108614 | CB076_DANRE | UPF0538 protein C2orf76 homolog | 6.33E-30 | 3-431[+] | 1.551339 | 4.875399 | 114.4255 | 3.03E-09 | 1.49E-05 |
| 86448 | SMCE1_MOUSE | SWI/SNF-related matrix-associated actin-dependent regulator of chromatin subfamily E member 1 | 3.11E-13 | 228-1004[+] | -1.87353 | 6.247321 | 98.34513 | 9.85E-09 | 4.23E-05 |
| 110342 |  |  |  |  | -2.40205 | 5.491794 | 96.57077 | 1.13E-08 | 4.32E-05 |
| 99029 | CDC37_DROVI | Hsp90 co-chaperone Cdc37 | 3.34E-91 | 140-1222[+] | 1.613666 | 7.855046 | 93.13988 | 1.49E-08 | 5.13E-05 |
| 81037 | STIP1_HUMAN | Stress-induced-phosphoprotein 1 | 1.41E-136 | 143-1111[+] | 1.910945 | 6.142955 | 90.15807 | 1.91E-08 | 5.98E-05 |
| 105630 | GHITM_HUMAN | Growth hormone-inducible transmembrane protein | 3.40E-108 | 123-1163[+] | 1.46372 | 6.324875 | 85.80455 | 2.78E-08 | 7.78E-05 |
| 92507 |  |  |  | 174-1259[+] | 1.40887 | 5.319792 | 85.16439 | 2.95E-08 | 7.78E-05 |
| 109078 | SET_HUMAN | SET | 2.40E-46 | 1-483[+] | -1.39141 | 6.8527 | 80.14001 | 4.64E-08 | 0.000114 |
| 96808 | UBE2A_MOUSE | Ubiquitin-conjugating enzyme E2 A | 1.06E-82 | 152-613[+] | 1.011058 | 5.774296 | 76.19659 | 6.75E-08 | 0.000155 |
| 93864 | AHSA1_HUMAN | Activator of 90 kDa heat shock protein ATPase homolog 1 | 2.96E-106 | 101-1174[+] | 2.935567 | 3.976932 | 75.13687 | 7.49E-08 | 0.000161 |
| 48125 | PF13383.9 | Methyltransferase domain | 3.60E-09 | 182-736[+] | -9.03923 | 4.269381 | 76.32621 | 1.83E-07 | 0.000369 |
| 84031 | PRUN1_MOUSE | Exopolyphosphatase PRUNE1 | 2.20E-24 | 1139-1768[+] | -1.67292 | 4.339537 | 62.90863 | 2.71E-07 | 0.000517 |
| 98668 | F10A1_CHICK | Hsc70-interacting protein | 4.14E-52 | 529-897[+] | 1.21532 | 6.527399 | 60.42764 | 3.6E-07 | 0.000635 |
| 103578 | KTR3_YEAST | Probable mannosyltransferase KTR3 | 4.27E-25 | 283-1347[+] | 1.273476 | 5.427458 | 60.20753 | 3.7E-07 | 0.000635 |
| 20983 | MARH6_MOUSE | E3 ubiquitin-protein ligase MARCHF6 | 0 | 167-3673[+] | 0.763059 | 6.503036 | 58.20545 | 4.69E-07 | 0.000736 |
| 105147 | HMG2_DROME | High mobility group protein DSP1 | 3.20E-59 | 119-739[+] | -1.308 | 7.580118 | 58.16186 | 4.71E-07 | 0.000736 |
| 89924 |  |  |  |  | 1.650298 | 4.355143 | 56.68541 | 5.64E-07 | 0.000842 |

**Table S5.** Selected DE transcripts (FDR < 0.001) associated with anterior injury (at 35°C) (A35 x CA35)

| **Transcript ID** | **UniProtKB / Pfam ID** | **Protein** | **E-value** | **Protein coordinates** | **logFC** | **logCPM** | **F** | **P value** | **FDR** |
| --- | --- | --- | --- | --- | --- | --- | --- | --- | --- |
| 104706 |  |  |  | 164-487[+] | -5.10594 | 5.598169 | 125.846 | 1.52E-09 | 5.23E-05 |
| 35544 | RIR1_MOUSE | Ribonucleoside-diphosphate reductase large subunit | 0 | 91-2499[+] | 1.66601 | 4.79965 | 107.609 | 4.9E-09 | 8.42E-05 |
| 98623 |  |  |  |  | -2.33851 | 5.209282 | 93.07389 | 1.5E-08 | 0.00013 |
| 98411 | CAH15_MOUSE | Carbonic anhydrase 15 | 1.57E-48 | 219-1187[+] | -1.87425 | 4.529742 | 92.13523 | 1.62E-08 | 0.00013 |
| 107740 |  |  |  | 85-900[+] | -3.25857 | 6.036317 | 90.77952 | 1.89E-08 | 0.00013 |
| 92647 | CATL2_HUMAN | Cathepsin L2 | 5.76E-74 | 121-1374[+] | -3.53544 | 4.251303 | 86.57239 | 2.6E-08 | 0.00014 |
| 102110 | VPP4_HUMAN | V-type proton ATPase 116 kDa subunit a isoform 4 | 9.56E-108 | 1-1392[+] | -1.9174 | 6.456291 | 85.56251 | 2.84E-08 | 0.00014 |
| 22453 | AAKG2_MOUSE | 5'-AMP-activated protein kinase subunit gamma-2 | 4.26E-70 | 109-3705[+] | 2.578414 | 4.981187 | 82.07861 | 3.89E-08 | 0.000167 |
| 111433 | CAT8_MOUSE | Cathepsin 8 | 5.18E-11 | 83-439[+] | -1.45026 | 7.125899 | 75.14959 | 7.48E-08 | 0.000262 |
| 105249 |  |  |  | 107-1120[+] | -5.35692 | 2.997025 | 74.96622 | 7.61E-08 | 0.000262 |
| 101016 | VA0D1_MOUSE | V-type proton ATPase subunit d 1 | 0 | 165-1211[+] | -1.78877 | 6.02138 | 72.02181 | 1.02E-07 | 0.000316 |
| 110166 | DYLT5_MOUSE | Dynein light chain Tctex-type 5 | 5.03E-11 | 446-1012[+] | -2.11241 | 4.88915 | 71.25593 | 1.1E-07 | 0.000316 |
| 91287 | PUNA_GEOSE | Purine nucleoside phosphorylase 1 | 1.97E-17 | 2-457[+] | -2.31125 | 5.401997 | 69.64589 | 1.3E-07 | 0.000345 |
| 109733 | RLT1_RHIO9 | Mucoricin | 2.85E-06 | 82-876[+] | -1.94821 | 5.6816 | 68.3799 | 1.49E-07 | 0.000366 |
| 33214 | S17B3_XENTR | Transcription factor Sox-17-beta.3 | 1.80E-26 | 383-3178[+] | -6.59461 | 1.927501 | 75.47229 | 1.82E-07 | 0.000405 |
| 73659 | VPP1_RAT | V-type proton ATPase 116 kDa subunit a1 | 0 | 1-2499[+] | -2.10986 | 5.262253 | 66.16949 | 1.89E-07 | 0.000405 |
| 111787 |  |  |  | 3-317[+] | -1.5458 | 7.3398 | 64.17189 | 2.35E-07 | 0.000475 |
| 107646 | CRYL1_PONAB | Lambda-crystallin homolog | 1.48E-67 | 81-1052[+] | -3.18658 | 4.264846 | 63.1788 | 2.63E-07 | 0.000502 |
| 91191 | FBP1_STRPU | Fibropellin-1 | 8.25E-37 | 36-1361[+] | -6.80864 | 2.801388 | 82.07929 | 3.03E-07 | 0.000534 |
| 106794 | HPGDS_CHICK | Hematopoietic prostaglandin D synthase | 3.64E-43 | 39-674[+] | -2.18042 | 5.984745 | 61.24083 | 3.31E-07 | 0.000534 |
| 105197 | APRR2_ARATH | Two-component response regulator-like APRR2 | 0.000139 | 287-745[+] | -1.82309 | 5.465303 | 60.36423 | 3.63E-07 | 0.000534 |
| 87030 |  |  |  | 259-1146[+] | -5.22013 | 4.534726 | 60.5688 | 3.74E-07 | 0.000534 |
| 86237 | CP2F3_CAPHI | Cytochrome P450 2F3 | 6.22E-68 | 71-1654[+] | -6.57703 | 2.06433 | 67.69667 | 3.75E-07 | 0.000534 |
| 110698 |  |  |  | 100-858[+] | -3.10683 | 4.671946 | 60.05287 | 3.81E-07 | 0.000534 |
| 96233 | PUR6_CHICK | Multifunctional protein ADE2 | 0 | 157-1425[+] | 2.356456 | 6.421227 | 59.96159 | 3.89E-07 | 0.000534 |
| 46538 |  |  |  | 360-872[+] | 3.758685 | 3.21735 | 58.67219 | 4.43E-07 | 0.000586 |
| 110851 | IF5A_DROME | Eukaryotic translation initiation factor 5A | 1.85E-59 | 45-548[+] | 0.639883 | 9.270789 | 57.05703 | 5.39E-07 | 0.000685 |
| 109959 | GRN_DICDI | Granulin | 2.90E-05 |  | 1.396759 | 9.363465 | 56.39421 | 5.84E-07 | 0.000717 |
| 32913 | ANKAR_MOUSE | Ankyrin and armadillo repeat-containing protein | 3.37E-34 | 239-3637[+] | -6.11223 | 2.597477 | 62.83714 | 6.1E-07 | 0.000722 |
| 70960 | OSTA_LEUER | Organic solute transporter subunit alpha | 1.63E-13 | 3-1184[+] | -1.59438 | 3.912291 | 53.81446 | 8.07E-07 | 0.000867 |
| 109797 | GRN_DICDI | Granulin | 2.26E-05 |  | 1.609146 | 9.313346 | 53.70235 | 8.28E-07 | 0.000867 |
| 80562 | CEL_BOVIN | Bile salt-activated lipase | 3.44E-65 | 222-2210[+] | -3.0049 | 3.530507 | 53.23576 | 8.7E-07 | 0.000867 |
| 48125 |  |  |  | 182-736[+] | 8.155488 | 4.269381 | 59.9025 | 8.86E-07 | 0.000867 |
| 84165 |  |  |  | 134-826[+] | -5.22758 | 4.490783 | 53.47325 | 8.9E-07 | 0.000867 |
| 110686 | VA0E1_RAT | V-type proton ATPase subunit e 1 | 1.29E-20 |  | -1.39156 | 6.476873 | 53.02653 | 8.93E-07 | 0.000867 |
| 10234 | LGR5_BOVIN | Leucine-rich repeat-containing G-protein coupled receptor 5 | 7.88E-17 | 1033-4074[+] | -6.88101 | 2.444933 | 58.98583 | 9.16E-07 | 0.000867 |
| 111461 | H2AV_XENTR | Histone H2A.V | 4.18E-64 | 55-441[+] | 1.183739 | 7.734511 | 52.68502 | 9.34E-07 | 0.000867 |
| 8722 | GVIN1_MOUSE | Interferon-induced very large GTPase 1 | 0 | 50-5014[+] | -2.19318 | 4.119865 | 52.03099 | 1.02E-06 | 0.000919 |
| 109725 |  |  |  | 140-826[+] | -2.56785 | 4.17308 | 51.26166 | 1.13E-06 | 0.000991 |

**Table S6.** Selected DE transcripts (FDR < 0.001) associated with thermal stress (in anteriorly injured worms) (A35 x A23)

| **Transcript ID** | **UniProtKB / Pfam ID** | **Protein** | **E-value** | **Protein coordinates** | **logFC** | **logCPM** | **F** | **P value** | **FDR** |
| --- | --- | --- | --- | --- | --- | --- | --- | --- | --- |
| 77307 | HSP7C_SAGOE | Heat shock cognate 71 kDa protein | 0 | 201-2153[+] | 4.017426 | 5.483423 | 147.8521 | 4.01E-10 | 1.38E-05 |
| 109908 | CRYAB_MACFA | Alpha-crystallin B chain | 1.34E-15 | 129-629[+] | 2.767965 | 6.691767 | 128.1878 | 1.23E-09 | 2.12E-05 |
| 100280 | MLF2_HUMAN | Myeloid leukemia factor 2 | 1.02E-38 | 138-899[+] | 2.883075 | 4.432257 | 101.8239 | 7.53E-09 | 6.66E-05 |
| 108040 | U2AF4_RAT | Splicing factor U2AF 26 kDa subunit | 1.51E-122 | 64-732[+] | 1.021279 | 6.861008 | 101.4409 | 7.76E-09 | 6.66E-05 |
| 110342 |  |  |  |  | -2.37797 | 5.491794 | 93.4582 | 1.46E-08 | 0.0001 |
| 108614 | CB076_DANRE | UPF0538 protein C2orf76 homolog | 6.33E-30 | 3-431[+] | 1.368476 | 4.875399 | 91.02239 | 1.78E-08 | 0.000102 |
| 99029 | CDC37_DROVI | Hsp90 co-chaperone Cdc37 | 3.34E-91 | 140-1222[+] | 1.573794 | 7.855046 | 88.61795 | 2.18E-08 | 0.000107 |
| 81037 | STIP1_HUMAN | Stress-induced-phosphoprotein 1 | 1.41E-136 | 143-1111[+] | 1.810509 | 6.142955 | 81.56137 | 4.07E-08 | 0.000163 |
| 111384 | CH10_ORYLA | 10 kDa heat shock protein, mitochondrial | 4.96E-43 | 116-421[+] | 1.553039 | 8.513635 | 81.04501 | 4.27E-08 | 0.000163 |
| 99284 | POSTN_MOUSE | Periostin | 1.81E-17 | 1-903[+] | 1.367487 | 5.974099 | 77.01001 | 6.24E-08 | 0.000214 |
| 108481 |  |  |  | 1-798[+] | -1.64577 | 6.2247 | 72.67391 | 9.56E-08 | 0.000299 |
| 102690 | SRCA_CHICK | Sarcalumenin | 5.84E-144 | 3-1508[+] | -1.40179 | 4.806425 | 70.56086 | 1.19E-07 | 0.000315 |
| 98338 | FACR1_DROME | Putative fatty acyl-CoA reductase CG5065 | 3.02E-95 | 72-1322[+] | -3.24524 | 5.618295 | 71.01324 | 1.19E-07 | 0.000315 |
| 84988 | TENX_HUMAN | Tenascin-X | 5.16E-29 | 98-2230[+] | 3.24073 | 6.805893 | 68.07196 | 1.62E-07 | 0.000398 |
| 108060 | Y2624_MYCTU | Universal stress protein Rv2624c | 1.07E-08 | 140-541[+] | 2.032426 | 5.312626 | 66.28725 | 1.86E-07 | 0.000427 |
| 105630 | GHITM_HUMAN | Growth hormone-inducible transmembrane protein | 3.40E-108 | 123-1163[+] | 1.25711 | 6.324875 | 63.94843 | 2.41E-07 | 0.000486 |
| 111193 | RLT1_RHIO9 | Mucoricin | 2.03E-05 | 1-381[+] | -0.98275 | 8.993963 | 63.94387 | 2.41E-07 | 0.000486 |
| 105262 |  |  |  |  | -2.01744 | 4.116542 | 62.86182 | 2.72E-07 | 0.000486 |
| 20983 | MARH6_MOUSE | E3 ubiquitin-protein ligase MARCHF6 | 0 | 167-3673[+] | 0.792511 | 6.503036 | 62.76487 | 2.75E-07 | 0.000486 |
| 100763 | FIBA_APOPA | Fibrinogen-like protein A | 2.02E-26 | 3-1265[+] | -2.22648 | 3.973629 | 62.44863 | 2.85E-07 | 0.000486 |
| 109733 | RLT1_RHIO9 | Mucoricin | 2.85E-06 | 82-876[+] | -1.84738 | 5.6816 | 61.69435 | 3.11E-07 | 0.000486 |
| 109625 |  |  |  | 3-809[+] | -1.74106 | 6.566724 | 61.69197 | 3.11E-07 | 0.000486 |
| 111639 |  |  |  |  | -1.10403 | 6.752587 | 57.5401 | 5.08E-07 | 0.00073 |
| 70519 | PLD3B_MACFA | PRELI domain containing protein 3B | 1.38E-64 | 90-674[+] | 1.432401 | 5.191892 | 57.26595 | 5.25E-07 | 0.00073 |
| 96224 |  |  |  |  | 1.697362 | 4.9415 | 57.175 | 5.31E-07 | 0.00073 |
| 98172 | WIPF1_MOUSE | WAS/WASL-interacting protein family member 1 | 0.000505 | 306-1298[+] | 1.561528 | 6.015115 | 55.68124 | 6.38E-07 | 0.000843 |
| 110596 |  |  |  |  | -1.03506 | 8.150899 | 55.15391 | 6.82E-07 | 0.00085 |
| 33214 | S17B3_XENTR | Transcription factor Sox-17-beta.3 | 1.80E-26 | 383-3178[+] | -5.89091 | 1.927501 | 61.54563 | 6.97E-07 | 0.00085 |
| 79013 | MIOX_DANRE | Inositol oxygenase | 1.16E-113 | 142-996[+] | -1.71088 | 7.339379 | 54.74167 | 7.18E-07 | 0.00085 |
| 110324 |  |  |  |  | -2.82414 | 3.589845 | 53.85541 | 8.03E-07 | 0.000902 |
| 4354 | TBC25_HUMAN | TBC1 domain family member 25 | 7.55E-108 | 3-2330[+] | 1.496825 | 4.076436 | 53.55587 | 8.35E-07 | 0.000902 |
| 107020 | BRK1B_XENLA | Probable protein BRICK1-B | 3.70E-30 |  | -1.65965 | 6.136484 | 53.50619 | 8.4E-07 | 0.000902 |

**Table S7.** Selected DE transcripts (FDR < 0.001, max 50) associated with combined anterior injury and thermal stress (A35 x CA23)

| **Transcript ID** | **UniProtKB / Pfam ID** | **Protein** | **E-value** | **Protein coordinates** | | **logFC** | **logCPM** | **F** | **P value** | **FDR** |
| --- | --- | --- | --- | --- | --- | --- | --- | --- | --- | --- |
| 111384 | CH10_ORYLA | 10 kDa heat shock protein, mitochondrial | 4.96E-43 | 116-421[+] | 2.390464 | | 8.513635 | 181.0038 | 7.51E-11 | 1.55E-06 |
| 98623 |  |  |  |  | -3.3047 | | 5.209282 | 176.2162 | 9.36E-11 | 1.55E-06 |
| 104706 |  |  |  | 164-487[+] | -6.0047 | | 5.598169 | 170.0697 | 1.35E-10 | 1.55E-06 |
| 109908 | CRYAB_MACFA | Alpha-crystallin B chain | 1.34E-15 | 129-629[+] | 3.10327 | | 6.691767 | 155.6504 | 2.58E-10 | 2.06E-06 |
| 106668 |  |  |  |  | -1.58693 | | 8.238701 | 152.817 | 3E-10 | 2.06E-06 |
| 98411 | CAH15_MOUSE | Carbonic anhydrase 15 | 1.57E-48 | 219-1187[+] | -2.35782 | | 4.529742 | 142.7344 | 5.21E-10 | 2.99E-06 |
| 108614 | CB076_DANRE | UPF0538 protein C2orf76 homolog | 6.33E-30 | 3-431[+] | 1.710804 | | 4.875399 | 138.1736 | 6.77E-10 | 3.32E-06 |
| 110598 | HPGDS_CHICK | Hematopoietic prostaglandin D synthase | 5.97E-47 | 65-739[+] | -2.26208 | | 4.627148 | 130.4669 | 1.07E-09 | 4.33E-06 |
| 105197 | APRR2_ARATH | Two-component response regulator-like APRR2 | 0.000139 | 287-745[+] | -2.72131 | | 5.465303 | 129.54 | 1.14E-09 | 4.33E-06 |
| 100763 | FIBA_APOPA | Fibrinogen-like protein A | 2.02E-26 | 3-1265[+] | -3.22164 | | 3.973629 | 126.0655 | 1.41E-09 | 4.84E-06 |
| 108040 | U2AF4_RAT | Splicing factor U2AF 26 kDa subunit | 1.51E-122 | 64-732[+] | 1.125859 | | 6.861008 | 122.6001 | 1.76E-09 | 5.49E-06 |
| 111433 | CAT8_MOUSE | Cathepsin 8 | 5.18E-11 | 83-439[+] | -1.80958 | | 7.125899 | 114.8422 | 2.95E-09 | 8.19E-06 |
| 91287 | PUNA_GEOSE | Purine nucleoside phosphorylase 1 | 1.97E-17 | 2-457[+] | -3.01576 | | 5.401997 | 114.1128 | 3.1E-09 | 8.19E-06 |
| 111787 |  |  |  | 3-317[+] | -2.06869 | | 7.3398 | 111.6382 | 3.68E-09 | 8.84E-06 |
| 97304 | MA2B1_MOUSE | Lysosomal alpha-mannosidase | 5.77E-20 | 2-481[+] | 1.252745 | | 6.902165 | 110.4183 | 4.01E-09 | 8.84E-06 |
| 77307 | HSP7C_SAGOE | Heat shock cognate 71 kDa protein | 0 | 201-2153[+] | 3.322287 | | 5.483423 | 110.3173 | 4.12E-09 | 8.84E-06 |
| 110342 |  |  |  |  | -2.53174 | | 5.491794 | 105.0568 | 5.91E-09 | 1.19E-05 |
| 102110 | VPP4_HUMAN | V-type proton ATPase 116 kDa subunit a isoform 4 | 9.56E-108 | 1-1392[+] | -2.10265 | | 6.456291 | 101.8114 | 7.54E-09 | 1.44E-05 |
| 107740 |  |  |  | 85-900[+] | -3.45183 | | 6.036317 | 100.574 | 8.62E-09 | 1.56E-05 |
| 22453 | AAKG2_MOUSE | 5'-AMP-activated protein kinase subunit gamma-2 | 4.26E-70 | 109-3705[+] | 2.865427 | | 4.981187 | 97.98776 | 1.01E-08 | 1.67E-05 |
| 46538 |  |  |  | 360-872[+] | 5.458466 | | 3.21735 | 97.40878 | 1.06E-08 | 1.67E-05 |
| 73659 | VPP1_RAT | V-type proton ATPase 116 kDa subunit a1 | 0 | 1-2499[+] | -2.58604 | | 5.262253 | 97.31525 | 1.07E-08 | 1.67E-05 |
| 97910 | T23O_ANOGA | Tryptophan 2,3-dioxygenase | 2.20E-148 | 3-1319[+] | -2.08832 | | 6.614242 | 96.7525 | 1.12E-08 | 1.67E-05 |
| 88942 | CKS1_MOUSE | Cyclin-dependent kinases regulatory subunit 1 | 1.42E-29 |  | 1.516922 | | 5.273971 | 95.92392 | 1.19E-08 | 1.71E-05 |
| 110698 | PF04103.18 | CD20-like family | 2.70E-07 | 100-858[+] | -3.862 | | 4.671946 | 90.00811 | 1.97E-08 | 2.62E-05 |
| 111639 |  |  |  |  | -1.38626 | | 6.752587 | 89.73018 | 1.99E-08 | 2.62E-05 |
| 105249 | PF07801.14 | Protein of unknown function (DUF1647) | 2.60E-15 | 107-1120[+] | -5.83592 | | 2.997025 | 88.85598 | 2.14E-08 | 2.72E-05 |
| 110686 | VA0E1_RAT | V-type proton ATPase subunit e 1 | 1.29E-20 |  | -1.81463 | | 6.476873 | 88.42853 | 2.22E-08 | 2.72E-05 |
| 91191 | FBP1_STRPU | Fibropellin-1 | 8.25E-37 | 36-1361[+] | -8.21738 | | 2.801388 | 122.8469 | 2.47E-08 | 2.93E-05 |
| 99284 | POSTN_MOUSE | Periostin | 1.81E-17 | 1-903[+] | 1.454337 | | 5.974099 | 86.4271 | 2.64E-08 | 2.99E-05 |
| 81037 | STIP1_HUMAN | Stress-induced-phosphoprotein 1 | 1.41E-136 | 143-1111[+] | 1.863027 | | 6.142955 | 85.8301 | 2.78E-08 | 2.99E-05 |
| 101016 | VA0D1_MOUSE | V-type proton ATPase subunit d 1 | 0 | 165-1211[+] | -1.9609 | | 6.02138 | 85.80336 | 2.78E-08 | 2.99E-05 |
| 87030 |  |  |  | 259-1146[+] | -6.26439 | | 4.534726 | 86.07145 | 2.91E-08 | 2.99E-05 |
| 98338 | FACR1_DROME | Putative fatty acyl-CoA reductase CG5065 | 3.02E-95 | 72-1322[+] | -3.59895 | | 5.618295 | 85.43692 | 3.05E-08 | 2.99E-05 |
| 107944 | COPT1_PONAB | High affinity copper uptake protein 1 | 2.58E-22 | 54-587[+] | -1.88225 | | 5.829838 | 84.752 | 3.06E-08 | 2.99E-05 |
| 102533 | GELS1_LUMTE | Gelsolin-like protein 1 | 6.55E-136 | 97-1221[+] | -2.33155 | | 6.168723 | 84.47587 | 3.13E-08 | 2.99E-05 |
| 84585 |  |  |  | 192-1685[+] | -1.26196 | | 5.401941 | 83.18803 | 3.51E-08 | 3.18E-05 |
| 109725 |  |  |  | 140-826[+] | -3.29367 | | 4.17308 | 83.17939 | 3.52E-08 | 3.18E-05 |
| 92647 | CATL2_HUMAN | Cathepsin L2 | 5.76E-74 | 121-1374[+] | -3.42745 | | 4.251303 | 81.56344 | 4.07E-08 | 3.59E-05 |
| 107646 | CRYL1_PONAB | Lambda-crystallin homolog | 1.48E-67 | 81-1052[+] | -3.63628 | | 4.264846 | 80.65908 | 4.43E-08 | 3.74E-05 |
| 92163 | ASH2L_HUMAN | Set1/Ash2 histone methyltransferase complex subunit ASH2 | 3.85E-174 | 141-1841[+] | 1.509098 | | 4.428873 | 80.56423 | 4.47E-08 | 3.74E-05 |
| 110166 | DYLT5_MOUSE | Dynein light chain Tctex-type 5 | 5.03E-11 | 446-1012[+] | -2.24911 | | 4.88915 | 80.21822 | 4.61E-08 | 3.77E-05 |
| 111694 |  |  |  | 76-435[+] | -4.32095 | | 5.494206 | 79.02894 | 5.49E-08 | 4.38E-05 |
| 109733 | RLT1_RHIO9 | Mucoricin | 2.85E-06 | 82-876[+] | -2.08769 | | 5.6816 | 77.97066 | 5.7E-08 | 4.38E-05 |
| 102019 | KMO_DANRE | Kynurenine 3-monooxygenase | 4.41E-176 | 101-1531[+] | -1.87691 | | 5.534159 | 77.68384 | 5.85E-08 | 4.38E-05 |
| 98668 | F10A1_CHICK | Hsc70-interacting protein | 4.14E-52 | 529-897[+] | 1.38292 | | 6.527399 | 77.66891 | 5.86E-08 | 4.38E-05 |
| 103035 |  |  |  | 137-895[+] | -5.78803 | | 2.821099 | 77.00844 | 6.24E-08 | 4.56E-05 |
| 31224 | PAR14_HUMAN | Protein mono-ADP-ribosyltransferase PARP14 | 5.29E-101 | 22-3069[+] | -2.35118 | | 3.597091 | 74.42569 | 8.03E-08 | 5.73E-05 |
| 106794 | HPGDS_CHICK | Hematopoietic prostaglandin D synthase | 3.64E-43 | 39-674[+] | -2.41853 | | 5.984745 | 74.34577 | 8.17E-08 | 5.73E-05 |
| 105630 | GHITM_HUMAN | Growth hormone-inducible transmembrane protein | 3.40E-108 | 123-1163[+] | 1.353593 | | 6.324875 | 73.64685 | 8.68E-08 | 5.96E-05 |

**Table S8.** Selected DE transcripts (FDR < 0.001) associated with posterior injury (P23 x CP23)

| **Transcript ID** | **UniProtKB / Pfam ID** | **Protein** | **E-value** | **Protein coordinates** | **logFC** | **logCPM** | **F** | **P value** | **FDR** |
| --- | --- | --- | --- | --- | --- | --- | --- | --- | --- |
| 111464 | PF17064.8 | Sleepless protein | 1.00E-06 | 94-495[+] | 11.41238 | 5.432951 | 379.9085 | 1.49E-11 | 2.59E-07 |
| 110872 |  |  |  | 1-489[+] | 11.42187 | 5.395081 | 379.06 | 1.51E-11 | 2.59E-07 |
| 75151 | DHE3_HUMAN | Glutamate dehydrogenase 1, mitochondrial | 0 | 151-1521[+] | 2.986453 | 7.794221 | 201.3073 | 3.11E-11 | 3.56E-07 |
| 111384 | CH10_ORYLA | 10 kDa heat shock protein, mitochondrial | 4.96E-43 | 116-421[+] | 2.058218 | 8.513635 | 137.0297 | 7.24E-10 | 5.85E-06 |
| 78425 | MOT12_HUMAN | Monocarboxylate transporter 12 | 1.50E-07 | 1-339[+] | 2.279155 | 4.80619 | 134.2967 | 8.51E-10 | 5.85E-06 |
| 96233 | PUR6_CHICK | Multifunctional protein ADE2 | 0 | 157-1425[+] | 3.712185 | 6.421227 | 130.5207 | 1.11E-09 | 6.13E-06 |
| 65815 | GLSK_RAT | Glutaminase kidney isoform, mitochondrial | 0 | 256-2238[+] | 8.868308 | 3.875601 | 194.7835 | 1.25E-09 | 6.13E-06 |
| 83206 | CH60_CHICK | 60 kDa heat shock protein, mitochondrial | 0 | 138-1877[+] | 2.662853 | 7.137562 | 116.7814 | 2.58E-09 | 1.11E-05 |
| 108837 | CHIA_MOUSE | Acidic mammalian chitinase | 1.31E-53 | 1-864[+] | 5.4376 | 4.965106 | 114.8411 | 3.18E-09 | 1.22E-05 |
| 19471 | MOT12_XENTR | Monocarboxylate transporter 12 | 5.31E-46 | 453-2324[+] | 2.546285 | 4.884283 | 108.0846 | 4.74E-09 | 1.63E-05 |
| 98055 | TSAL_GEOSL | L-threonine ammonia-lyase | 1.20E-52 | 59-1408[+] | 8.988341 | 3.287691 | 139.6821 | 1.09E-08 | 3.4E-05 |
| 98411 | CAH15_MOUSE | Carbonic anhydrase 15 | 1.57E-48 | 219-1187[+] | -1.87347 | 4.529742 | 94.87829 | 1.3E-08 | 3.71E-05 |
| 101662 | SFXN1_MOUSE | Sideroflexin-1 | 3.78E-68 | 3-566[+] | 1.821735 | 6.091986 | 91.71967 | 1.68E-08 | 4.44E-05 |
| 99348 |  |  |  | 127-543[+] | 7.748744 | 5.029388 | 86.15183 | 2.92E-08 | 7.16E-05 |
| 110736 | CNFN_XENTR | Cornifelin homolog | 1.79E-12 | 217-651[+] | 3.860537 | 4.081214 | 82.59673 | 3.71E-08 | 8.49E-05 |
| 85010 | HGNAT_MOUSE | Heparan-alpha-glucosaminide N-acetyltransferase | 1.16E-117 | 156-2096[+] | -1.44514 | 5.674129 | 80.8038 | 4.37E-08 | 9.38E-05 |
| 110333 | CAPSL_MOUSE | Calcyphosin-like protein | 2.85E-60 | 222-854[+] | 1.426146 | 5.462118 | 76.68823 | 6.44E-08 | 0.00013 |
| 103759 | GCSH_DROME | Glycine cleavage system H protein, mitochondrial | 1.90E-45 | 98-607[+] | 1.343571 | 6.117554 | 71.97304 | 1.03E-07 | 0.000188 |
| 94763 | TSAL_GEOSL | L-threonine ammonia-lyase | 2.36E-52 | 164-1513[+] | 6.367384 | 3.482888 | 71.96221 | 1.04E-07 | 0.000188 |
| 105717 |  |  |  | 159-1175[+] | 3.787605 | 4.246836 | 70.51955 | 1.21E-07 | 0.000208 |
| 91701 |  |  |  | 103-954[+] | 7.082076 | 2.591428 | 91.95599 | 1.51E-07 | 0.000248 |
| 110871 |  |  |  | 183-620[+] | 5.361959 | 4.138225 | 68.1194 | 1.6E-07 | 0.00025 |
| 105744 | FIBA_APOPA | Fibrinogen-like protein A | 0.001 | 2-1264[+] | -2.67943 | 4.137367 | 66.34689 | 1.85E-07 | 0.000277 |
| 97432 | PUR6_CHICK | Multifunctional protein ADE2 | 0 | 132-1400[+] | 3.681636 | 6.087589 | 66.03441 | 2.04E-07 | 0.000292 |
| 101323 |  |  |  | 206-1474[+] | 4.094049 | 5.521143 | 65.63088 | 2.13E-07 | 0.000293 |
| 35544 | RIR1_MOUSE | Ribonucleoside-diphosphate reductase large subunit | 0 | 91-2499[+] | 1.270267 | 4.79965 | 63.41512 | 2.56E-07 | 0.000338 |
| 79043 | GRP75_PONAB | Stress-70 protein, mitochondrial | 0 | 79-2124[+] | 1.965587 | 5.369608 | 62.22121 | 2.93E-07 | 0.000361 |
| 97174 | ASGL1_HUMAN | Isoaspartyl peptidase/L-asparaginase | 6.19E-86 | 355-1371[+] | 8.090432 | 3.737137 | 82.73233 | 2.94E-07 | 0.000361 |
| 73818 |  |  |  |  | 7.550435 | 2.609089 | 67.57224 | 3.79E-07 | 0.000449 |
| 55949 | TECTA_MOUSE | Alpha-tectorin | 4.01E-33 | 106-2607[+] | -2.30322 | 5.150658 | 55.18488 | 6.83E-07 | 0.000782 |
| 108398 | RET1_RAT | Retinol-binding protein 1 | 9.67E-05 | 122-577[+] | 1.262302 | 6.329141 | 54.19737 | 7.69E-07 | 0.000852 |
| 48789 | SC5A8_MOUSE | Sodium-coupled monocarboxylate transporter 1 | 1.80E-84 | 288-2048[+] | 6.15038 | 4.575293 | 54.08343 | 8.23E-07 | 0.000884 |

**Table S9.** Selected DE transcripts (FDR < 0.001) associated with thermal stress in anterior fragments (CP35 x CP23)

| **Transcript ID** | **UniProtKB / Pfam ID** | **Protein** | **E-value** | **Protein coordinates** | **logFC** | **logCPM** | **F** | **P value** | **FDR** |
| --- | --- | --- | --- | --- | --- | --- | --- | --- | --- |
| 111384 | CH10_ORYLA | 10 kDa heat shock protein, mitochondrial | 4.96E-43 | 116-421[+] | 2.542695 | 8.513635 | 202.0457 | 3.02E-11 | 1.04E-06 |
| 81037 | STIP1_HUMAN | Stress-induced-phosphoprotein 1 | 1.41E-136 | 143-1111[+] | 2.511713 | 6.142955 | 147.8263 | 3.93E-10 | 6.75E-06 |
| 109908 | CRYAB_MACFA | Alpha-crystallin B chain | 1.34E-15 | 129-629[+] | 2.842926 | 6.691767 | 133.6341 | 8.85E-10 | 1.01E-05 |
| 108614 | CB076_DANRE | UPF0538 protein C2orf76 homolog | 6.33E-30 | 3-431[+] | 1.59619 | 4.875399 | 120.6725 | 1.99E-09 | 1.71E-05 |
| 99284 | POSTN_MOUSE | Periostin | 1.81E-17 | 1-903[+] | 1.693887 | 5.974099 | 117.1709 | 2.52E-09 | 1.73E-05 |
| 108040 | U2AF4_RAT | Splicing factor U2AF 26 kDa subunit | 1.51E-122 | 64-732[+] | 1.080995 | 6.861008 | 113.0869 | 3.33E-09 | 1.9E-05 |
| 86448 | SMCE1_MOUSE | SWI/SNF-related matrix-associated actin-dependent regulator of chromatin subfamily E member 1 | 3.11E-13 | 228-1004[+] | -1.94086 | 6.247321 | 104.7184 | 6.06E-09 | 2.97E-05 |
| 84988 | TENX_HUMAN | Tenascin-X | 5.16E-29 | 98-2230[+] | 3.952697 | 6.805893 | 102.3411 | 7.78E-09 | 3.34E-05 |
| 99029 | CDC37_DROVI | Hsp90 co-chaperone Cdc37 | 3.34E-91 | 140-1222[+] | 1.660808 | 7.855046 | 98.20825 | 9.96E-09 | 3.8E-05 |
| 101878 | DDX47_HUMAN | Probable ATP-dependent RNA helicase DDX47 | 0 | 94-1425[+] | 1.199492 | 4.957791 | 95.17739 | 1.27E-08 | 4.35E-05 |
| 103787 | SRSF7_HUMAN | Serine/arginine-rich splicing factor 7 | 1.66E-19 | 89-862[+] | 1.489683 | 5.757852 | 88.38026 | 2.23E-08 | 6.95E-05 |
| 110342 |  |  |  |  | -2.19513 | 5.491794 | 84.14404 | 3.23E-08 | 9.23E-05 |
| 83206 | CH60_CHICK | 60 kDa heat shock protein, mitochondrial | 0 | 138-1877[+] | 2.184468 | 7.137562 | 81.10891 | 4.25E-08 | 0.000104 |
| 105630 | GHITM_HUMAN | Growth hormone-inducible transmembrane protein | 3.40E-108 | 123-1163[+] | 1.409665 | 6.324875 | 79.8691 | 4.76E-08 | 0.000104 |
| 20983 | MARH6_MOUSE | E3 ubiquitin-protein ligase MARCHF6 | 0 | 167-3673[+] | 0.895944 | 6.503036 | 79.81057 | 4.79E-08 | 0.000104 |
| 109078 | SET_HUMAN | Protein SET | 2.40E-46 | 1-483[+] | -1.39047 | 6.8527 | 79.64733 | 4.86E-08 | 0.000104 |
| 77307 | HSP7C_SAGOE | Heat shock cognate 71 kDa protein | 0 | 201-2153[+] | 2.71944 | 5.483423 | 77.49529 | 6.05E-08 | 0.000122 |
| 98668 | F10A1_CHICK | Hsc70-interacting protein | 4.14E-52 | 529-897[+] | 1.336435 | 6.527399 | 72.47677 | 9.76E-08 | 0.000186 |
| 110722 |  |  |  |  | -3.05007 | 7.394428 | 71.95855 | 1.09E-07 | 0.000197 |
| 111639 |  |  |  |  | -1.16444 | 6.752587 | 64.50752 | 2.27E-07 | 0.000389 |
| 110596 |  |  |  |  | -1.09966 | 8.150899 | 62.06127 | 2.98E-07 | 0.000488 |
| 93864 | AHSA1_HUMAN | Activator of 90 kDa heat shock protein ATPase homolog 1 | 2.96E-106 | 101-1174[+] | 2.576564 | 3.976932 | 59.26629 | 4.13E-07 | 0.000645 |
| 103029 |  |  |  | 1-849[+] | 1.696059 | 8.297292 | 58.15529 | 4.72E-07 | 0.000682 |
| 99989 | FKBP4_HUMAN | Peptidyl-prolyl cis-trans isomerase FKBP4 | 1.04E-116 | 132-1595[+] | 2.264978 | 6.557439 | 58.20937 | 4.77E-07 | 0.000682 |

**Table S10.** Selected DE transcripts (FDR < 0.001) associated with thermal stress (in posteriorly injured worms) (P35 x P23)

| **Transcript ID** | **UniProtKB / Pfam ID** | **Protein** | **E-value** | **Protein coordinates** | **logFC** | **logCPM** | **F** | **P value** | **FDR** |
| --- | --- | --- | --- | --- | --- | --- | --- | --- | --- |
| 108614 | CB076_DANRE | UPF0538 protein C2orf76 homolog | 6.33E-30 | 3-431[+] | 1.621396 | 4.875399 | 126.5038 | 1.37E-09 | 2.44E-05 |
| 109908 | CRYAB_MACFA | Alpha-crystallin B chain | 1.34E-15 | 129-629[+] | 2.73551 | 6.691767 | 125.9326 | 1.42E-09 | 2.44E-05 |
| 77307 | HSP7C_SAGOE | Heat shock cognate 71 kDa protein | 0 | 201-2153[+] | 3.216757 | 5.483423 | 108.099 | 4.82E-09 | 5.52E-05 |
| 110342 |  |  |  |  | -2.37823 | 5.491794 | 96.23014 | 1.16E-08 | 8.97E-05 |
| 99284 | POSTN_MOUSE | Periostin | 1.81E-17 | 1-903[+] | 1.517142 | 5.974099 | 94.68573 | 1.32E-08 | 8.97E-05 |
| 99029 | CDC37_DROVI | Hsp90 co-chaperone Cdc37 | 3.34E-91 | 140-1222[+] | 1.607016 | 7.855046 | 92.56229 | 1.57E-08 | 8.97E-05 |
| 81037 | STIP1_HUMAN | Stress-induced-phosphoprotein 1 | 1.41E-136 | 143-1111[+] | 1.893449 | 6.142955 | 90.00852 | 1.94E-08 | 9.51E-05 |
| 111639 |  |  |  |  | -1.33402 | 6.752587 | 83.65943 | 3.37E-08 | 0.000132 |
| 108040 | U2AF4_RAT | Splicing factor U2AF 26 kDa subunit | 1.51E-122 | 64-732[+] | 0.923519 | 6.861008 | 83.35667 | 3.46E-08 | 0.000132 |
| 105630 | GHITM_HUMAN | Growth hormone-inducible transmembrane protein | 3.40E-108 | 123-1163[+] | 1.40584 | 6.324875 | 80.17852 | 4.63E-08 | 0.000159 |
| 110596 |  |  |  |  | -1.24436 | 8.150899 | 79.03222 | 5.15E-08 | 0.000161 |
| 86448 | SMCE1_MOUSE | SWI/SNF-related matrix-associated actin-dependent regulator of chromatin subfamily E member 1 | 3.11E-13 | 228-1004[+] | -1.5876 | 6.247321 | 72.4896 | 9.74E-08 | 0.000279 |
| 109078 | SET_HUMAN | Protein SET | 2.40E-46 | 1-483[+] | -1.30319 | 6.8527 | 70.89898 | 1.15E-07 | 0.000303 |
| 100280 | MLF2_HUMAN | Myeloid leukemia factor 2 | 1.02E-38 | 138-899[+] | 2.277786 | 4.432257 | 69.0865 | 1.38E-07 | 0.000339 |
| 20983 | MARH6_MOUSE | E3 ubiquitin-protein ligase MARCHF6 | 0 | 167-3673[+] | 0.821531 | 6.503036 | 67.58881 | 1.62E-07 | 0.000371 |

**Table S11.** Selected DE transcripts (FDR < 0.001) associated with posterior injury (at 35°C) (P35 x CP35)

| **Transcript ID** | **UniProtKB / Pfam ID** | **Protein** | **E-value** | **Protein coordinates** | **logFC** | **logCPM** | **F** | **P value** | **FDR** |
| --- | --- | --- | --- | --- | --- | --- | --- | --- | --- |
| 111464 | PF17064.8 | Sleepless protein | 1.00E-06 | 94-495[+] | 11.34667 | 5.432951 | 388.3452 | 1.28E-11 | 3.32E-07 |
| 110872 |  |  |  | 1-489[+] | 11.05053 | 5.395081 | 365.4093 | 1.93E-11 | 3.32E-07 |
| 75151 |  |  |  | 151-1521[+] | 2.848323 | 7.794221 | 185.563 | 6.11E-11 | 7E-07 |
| 65815 | GLSK_RAT | Glutaminase kidney isoform, mitochondrial | 0 | 256-2238[+] | 9.402309 | 3.875601 | 230.4827 | 4.1E-10 | 3.52E-06 |
| 108837 | CHIA_MOUSE | Acidic mammalian chitinase | 1.31E-53 | 1-864[+] | 5.592593 | 4.965106 | 116.9142 | 2.77E-09 | 1.66E-05 |
| 78425 | MOT12_HUMAN | Monocarboxylate transporter 12 | 1.50E-07 | 1-339[+] | 2.095513 | 4.80619 | 115.0716 | 2.9E-09 | 1.66E-05 |
| 110871 |  |  |  | 183-620[+] | 6.947139 | 4.138225 | 99.82968 | 9.33E-09 | 4.03E-05 |
| 107185 | CDD_MOUSE | Cytidine deaminase | 3.19E-46 | 123-533[+] | 1.616957 | 4.652353 | 98.96031 | 9.39E-09 | 4.03E-05 |
| 101662 | SFXN1_MOUSE | Sideroflexin-1 | 3.78E-68 | 3-566[+] | 1.824047 | 6.091986 | 92.58425 | 1.56E-08 | 5.97E-05 |
| 105717 |  |  |  | 159-1175[+] | 4.449639 | 4.246836 | 85.37524 | 2.94E-08 | 0.000101 |
| 98055 | TSAL_GEOSL | L-threonine ammonia-lyase | 1.20E-52 | 59-1408[+] | 7.84665 | 3.287691 | 109.5698 | 5.09E-08 | 0.000158 |
| 99348 |  |  |  | 127-543[+] | 7.949248 | 5.029388 | 79.0615 | 5.52E-08 | 0.000158 |
| 97174 | ASGL1_HUMAN | Isoaspartyl peptidase/L-asparaginase | 6.19E-86 | 355-1371[+] | 8.851938 | 3.737137 | 106.0255 | 6.39E-08 | 0.000164 |
| 91701 |  |  |  | 103-954[+] | 7.345529 | 2.591428 | 104.483 | 6.85E-08 | 0.000164 |
| 19471 | MOT12_XENTR | Monocarboxylate transporter 12 | 5.31E-46 | 453-2324[+] | 2.071564 | 4.884283 | 75.581 | 7.17E-08 | 0.000164 |
| 103759 | GCSH_DROME | Glycine cleavage system H protein, mitochondrial | 1.90E-45 | 98-607[+] | 1.273255 | 6.117554 | 65.53787 | 2.02E-07 | 0.000434 |
| 92523 | ALAT2_XENTR | Alanine aminotransferase 2 | 0 | 136-1695[+] | 3.761059 | 5.047545 | 65.07918 | 2.25E-07 | 0.000455 |
| 76615 | COCA1_MOUSE | Collagen alpha-1(XII) chain | 1.28E-22 | 51-1688[+] | 4.080907 | 4.950472 | 61.87903 | 3.23E-07 | 0.000617 |
| 106668 |  |  |  |  | 0.976139 | 8.238701 | 59.8857 | 3.84E-07 | 0.000694 |
| 98411 | CAH15_MOUSE | Carbonic anhydrase 15 | 1.57E-48 | 219-1187[+] | -1.44681 | 4.529742 | 57.76908 | 4.94E-07 | 0.000849 |
| 94763 | TSAL_GEOSL | L-threonine ammonia-lyase | 2.36E-52 | 164-1513[+] | 5.992093 | 3.482888 | 56.67446 | 5.7E-07 | 0.000933 |

**Table S12.** Selected DE transcripts (FDR < 0.001, max 50) associated with combined posterior injury and thermal stress (P35 x CP23)

| **Transcript ID** | **UniProtKB / Pfam ID** | **Protein** | **E-value** | **Protein coordinates** | **logFC** | **logCPM** | **F** | **P value** | **FDR** |
| --- | --- | --- | --- | --- | --- | --- | --- | --- | --- |
| 111384 | CH10_ORYLA | 10 kDa heat shock protein, mitochondrial | 4.96E-43 | 116-421[+] | 2.765223 | 8.513635 | 234.915 | 8.54E-12 | 2.78E-07 |
| 111464 | PF17064.8 | Sleepless protein | 1.00E-06 | 94-495[+] | 11.34667 | 5.432951 | 375.1865 | 1.62E-11 | 2.78E-07 |
| 110872 |  |  |  | 1-489[+] | 11.05053 | 5.395081 | 352.4341 | 2.47E-11 | 2.82E-07 |
| 75151 | DHE3_HUMAN | Glutamate dehydrogenase 1, mitochondrial | 0 | 151-1521[+] | 2.873574 | 7.794221 | 188.1282 | 5.46E-11 | 4.68E-07 |
| 81037 | STIP1_HUMAN | Stress-induced-phosphoprotein 1 | 1.41E-136 | 143-1111[+] | 2.760998 | 6.142955 | 175.8944 | 9.5E-11 | 6.53E-07 |
| 111639 |  |  |  |  | -1.84449 | 6.752587 | 155.7052 | 2.58E-10 | 1.48E-06 |
| 83206 | CH60_CHICK | 60 kDa heat shock protein, mitochondrial | 0 | 138-1877[+] | 3.036281 | 7.137562 | 147.6354 | 3.97E-10 | 1.82E-06 |
| 110342 |  |  |  |  | -2.99423 | 5.491794 | 146.4257 | 4.24E-10 | 1.82E-06 |
| 65815 | GLSK_RAT | Glutaminase kidney isoform, mitochondrial | 0 | 256-2238[+] | 9.402309 | 3.875601 | 221.4316 | 5.35E-10 | 2.04E-06 |
| 109908 | CRYAB_MACFA | Alpha-crystallin B chain | 1.34E-15 | 129-629[+] | 2.813381 | 6.691767 | 131.2453 | 1.02E-09 | 3.51E-06 |
| 85010 | HGNAT_MOUSE | Heparan-alpha-glucosaminide N-acetyltransferase | 1.16E-117 | 156-2096[+] | -1.84557 | 5.674129 | 128.4444 | 1.21E-09 | 3.75E-06 |
| 108614 | CB076_DANRE | UPF0538 protein C2orf76 homolog | 6.33E-30 | 3-431[+] | 1.637456 | 4.875399 | 127.2344 | 1.31E-09 | 3.75E-06 |
| 109733 | RLT1_RHIO9 | Mucoricin | 2.85E-06 | 82-876[+] | -2.64122 | 5.6816 | 123.349 | 1.68E-09 | 4.43E-06 |
| 77307 | HSP7C_SAGOE | Heat shock cognate 71 kDa protein | 0 | 201-2153[+] | 3.375343 | 5.483423 | 114.9464 | 2.98E-09 | 7.32E-06 |
| 101878 | DDX47_HUMAN | Probable ATP-dependent RNA helicase DDX47 | 0 | 94-1425[+] | 1.301876 | 4.957791 | 112.3725 | 3.5E-09 | 8.01E-06 |
| 99029 | CDC37_DROVI | Hsp90 co-chaperone Cdc37 | 3.34E-91 | 140-1222[+] | 1.770842 | 7.855046 | 111.0026 | 3.85E-09 | 8.26E-06 |
| 78425 | MOT12_HUMAN | Monocarboxylate transporter 12 | 1.50E-07 | 1-339[+] | 1.993242 | 4.80619 | 104.2868 | 6.26E-09 | 1.26E-05 |
| 101662 | SFXN1_MOUSE | Sideroflexin-1 | 3.78E-68 | 3-566[+] | 1.92207 | 6.091986 | 101.6599 | 7.63E-09 | 1.41E-05 |
| 98411 | CAH15_MOUSE | Carbonic anhydrase 15 | 1.57E-48 | 219-1187[+] | -1.92848 | 4.529742 | 100.6569 | 8.23E-09 | 1.41E-05 |
| 19471 | MOT14_MOUSE | Monocarboxylate transporter 14 | 9.20E-28 | 453-2324[+] | 2.443171 | 4.884283 | 100.1933 | 8.53E-09 | 1.41E-05 |
| 108837 | CHIA_MOUSE | Acidic mammalian chitinase | 1.31E-53 | 1-864[+] | 5.039694 | 4.965106 | 100.9761 | 8.63E-09 | 1.41E-05 |
| 96403 | FCN2_HUMAN | Ficolin-2 | 1.17E-58 | 121-999[+] | -4.14305 | 3.51571 | 98.30356 | 9.88E-09 | 1.54E-05 |
| 105630 | GHITM_HUMAN | Growth hormone-inducible transmembrane protein | 3.40E-108 | 123-1163[+] | 1.540744 | 6.324875 | 95.01228 | 1.28E-08 | 1.92E-05 |
| 105744 | FIBA_APOPA | Fibrinogen-like protein A | 0.001 | 2-1264[+] | -3.2844 | 4.137367 | 93.92119 | 1.4E-08 | 2.01E-05 |
| 103759 | GCSH_DROME | Glycine cleavage system H protein, mitochondrial | 1.90E-45 | 98-607[+] | 1.529375 | 6.117554 | 92.67653 | 1.55E-08 | 2.13E-05 |
| 110596 |  |  |  |  | -1.33093 | 8.150899 | 90.00443 | 1.94E-08 | 2.49E-05 |
| 108040 | U2AF1_BOVIN | Splicing factor U2AF 35 kDa subunit | 9.49E-113 | 64-732[+] | 0.961449 | 6.861008 | 89.91055 | 1.95E-08 | 2.49E-05 |
| 110871 |  |  |  | 183-620[+] | 6.191528 | 4.138225 | 87.13494 | 2.62E-08 | 3.21E-05 |
| 111705 |  |  |  | 55-423[+] | -1.86433 | 6.969525 | 81.11622 | 4.24E-08 | 5.03E-05 |
| 110333 | CAPSL_MOUSE | Calcyphosin-like protein | 2.85E-60 | 222-854[+] | 1.444778 | 5.462118 | 78.74093 | 5.29E-08 | 6.06E-05 |
| 98055 | TSAL_GEOSL | L-threonine ammonia-lyas | 1.20E-52 | 59-1408[+] | 7.84665 | 3.287691 | 104.2505 | 6.95E-08 | 7.55E-05 |
| 110598 | HPGDS_CHICK | Hematopoietic prostaglandin D synthase | 5.97E-47 | 65-739[+] | -1.6368 | 4.627148 | 75.78369 | 7.03E-08 | 7.55E-05 |
| 109811 |  |  |  | 1-456[+] | 2.111746 | 4.975141 | 75.27661 | 7.39E-08 | 7.69E-05 |
| 110736 | CNFN_XENTR | Cornifelin homolog | 1.79E-12 | 217-651[+] | 3.656831 | 4.081214 | 74.93767 | 7.64E-08 | 7.71E-05 |
| 97174 | ASGL1_HUMAN | Isoaspartyl peptidase/L-asparaginase | 6.19E-86 | 355-1371[+] | 8.851938 | 3.737137 | 101.5208 | 8.37E-08 | 8.21E-05 |
| 91701 |  |  |  | 103-954[+] | 7.345529 | 2.591428 | 99.46209 | 9.31E-08 | 8.69E-05 |
| 111027 | HEMTN_HIRME | Neurohemerythrin | 2.84E-48 | 121-483[+] | -1.97599 | 9.452163 | 73.10526 | 9.36E-08 | 8.69E-05 |
| 100763 | FIBA_APOPA | Fibrinogen-like protein A | 2.02E-26 | 3-1265[+] | -2.2653 | 3.973629 | 72.44551 | 9.79E-08 | 8.85E-05 |
| 103787 | SRSF7_HUMAN | Serine/arginine-rich splicing factor 7 | 1.66E-19 | 89-862[+] | 1.340151 | 5.757852 | 72.10078 | 1.01E-07 | 8.85E-05 |
| 111193 | RLT1_RHIO9 | Mucoricin | 2.03E-05 | 1-381[+] | -1.0399 | 8.993963 | 71.4748 | 1.08E-07 | 8.85E-05 |
| 102867 |  |  |  | 3-743[+] | -3.70894 | 4.422183 | 71.64116 | 1.09E-07 | 8.85E-05 |
| 86448 | SMCE1_MOUSE | SWI/SNF-related matrix-associated actin-dependent regulator of chromatin subfamily E member 1 | 3.11E-13 | 228-1004[+] | -1.57266 | 6.247321 | 71.03724 | 1.13E-07 | 8.85E-05 |
| 111238 | SIAE_MOUSE | Sialate O-acetylesterase | 9.93E-26 | 3-563[+] | -2.50104 | 5.87043 | 71.18651 | 1.13E-07 | 8.85E-05 |
| 98668 | F10A1_CHICK | Hsc70-interacting protein | 4.14E-52 | 529-897[+] | 1.321561 | 6.527399 | 71.0003 | 1.13E-07 | 8.85E-05 |
| 99989 | FKBP4_HUMAN | Peptidyl-prolyl cis-trans isomerase FKBP4 | 1.04E-116 | 132-1595[+] | 2.519116 | 6.557439 | 70.87245 | 1.17E-07 | 8.94E-05 |
| 99348 |  |  |  | 127-543[+] | 6.983743 | 5.029388 | 70.45567 | 1.28E-07 | 9.57E-05 |
| 108398 | RET1_RAT | Retinol-binding protein 1 | 9.67E-05 | 122-577[+] | 1.433201 | 6.329141 | 69.48337 | 1.33E-07 | 9.69E-05 |
| 88942 | CKS1_MOUSE | Cyclin-dependent kinases regulatory subunit 1 | 1.42E-29 |  | 1.28089 | 5.273971 | 68.17036 | 1.52E-07 | 0.000109 |
| 4822 | SPPL3_MOUSE | Signal peptide peptidase-like 3 | 2.25E-120 | 178-1452[+] | 1.471924 | 5.539888 | 67.81241 | 1.58E-07 | 0.000111 |
| 101323 |  |  |  | 206-1474[+] | 4.166293 | 5.521143 | 67.83187 | 1.68E-07 | 0.000116 |

**Table S13.** Selected DE transcripts (FDR < 0.001) associated with posterior versus anterior body fragments (CA23 x CP23)

| **Transcript ID** | **UniProtKB / Pfam ID** | **Protein** | **E-value** | **Protein coordinates** | **logFC** | **logCPM** | **F** | **P value** | **FDR** |
| --- | --- | --- | --- | --- | --- | --- | --- | --- | --- |
| 110872 |  |  |  | 1-489[+] | 9.250092577 | 5.395081 | 227.5491 | 4.62E-10 | 1.11E-05 |
| 111464 | PF17064.8 | Sleepless protein | 1.00E-06 | 94-495[+] | 9.066498742 | 5.432951 | 216.35591 | 6.46E-10 | 1.11E-05 |
| 65815 | GLSK_RAT | Glutaminase kidney isoform, mitochondrial | 0 | 256-2238[+] | 8.500284729 | 3.8756008 | 176.65621 | 2.37E-09 | 2.72E-05 |
| 110778 |  |  |  | 2-394[+] | -3.66767379 | 4.0408414 | 96.382836 | 1.15E-08 | 9.88E-05 |
| 110736 | CNFN_XENTR | Cornifelin homolog | 1.79E-12 | 217-651[+] | 4.126679425 | 4.0812143 | 93.016474 | 1.51E-08 | 0.000104 |
| 29004 | PHP3_CAEEL | Homeobox protein php-3 | 1.35E-20 | 125-1786[+] | 2.809894579 | 3.9573868 | 89.470147 | 2.03E-08 | 0.000105 |
| 111744 |  |  |  | 78-410[+] | -2.76062134 | 4.6909984 | 88.807122 | 2.15E-08 | 0.000105 |
| 91701 |  |  |  | 103-954[+] | 7.998835259 | 2.5914279 | 118.07895 | 3.18E-08 | 0.000128 |
| 101323 |  |  |  | 206-1474[+] | 4.713344201 | 5.5211429 | 84.255602 | 3.45E-08 | 0.000128 |
| 109520 |  |  |  | 70-759[+] | -1.3863853 | 6.7216249 | 82.545751 | 3.72E-08 | 0.000128 |
| 109025 | PI16_MOUSE | Peptidase inhibitor 16 | 9.83E-33 | 120-938[+] | -5.216424 | 6.0503044 | 79.64825 | 5.23E-08 | 0.000163 |
| 98055 | TSAL_GEOSL | L-threonine ammonia-lyase | 1.20E-52 | 59-1408[+] | 7.915674901 | 3.287691 | 105.96541 | 6.28E-08 | 0.00018 |
| 111490 |  |  |  | 58-567[+] | -5.56069267 | 5.1367829 | 72.87501 | 1E-07 | 0.000265 |
| 110035 |  |  |  | 42-812[+] | -6.04559056 | 4.4412115 | 72.02652 | 1.09E-07 | 0.000267 |
| 106397 |  |  |  | 78-554[+] | -9.08368312 | 2.9636069 | 166.15167 | 1.4E-07 | 0.000306 |
| 75151 | DHE3_HUMAN | Glutamate dehydrogenase 1 | 0 | 151-1521[+] | 1.668907666 | 7.7942205 | 68.807849 | 1.42E-07 | 0.000306 |
| 106621 | TX53A_ETHRU | U-scoloptoxin(05)-Er3a | 0.000427 | 74-511[+] | -1.85976779 | 6.1074206 | 68.128027 | 1.53E-07 | 0.000309 |
| 97174 | ASGL1_HUMAN | Isoaspartyl peptidase/L-asparaginase | 6.19E-86 | 355-1371[+] | 8.396546072 | 3.7371368 | 89.956078 | 1.76E-07 | 0.000337 |
| 70009 |  |  |  | 2560-2889[-] | -1.10967967 | 5.2055569 | 63.171398 | 2.63E-07 | 0.000463 |
| 108890 |  |  |  | 158-541[+] | -1.82338368 | 5.5281834 | 62.955038 | 2.7E-07 | 0.000463 |
| 110095 |  |  |  | 63-842[+] | -8.34037953 | 2.497638 | 137.91264 | 3.4E-07 | 0.000546 |
| 108837 | CHIA_MOUSE | Acidic mammalian chitinase | 1.31E-53 | 1-864[+] | 3.821258351 | 4.9651062 | 61.106331 | 3.5E-07 | 0.000546 |
| 99348 |  |  |  | 127-543[+] | 6.461864622 | 5.0293876 | 59.956316 | 4.04E-07 | 0.000583 |
| 45987 | PF14295.9 | PAN domain | 0.011 | 167-2392[+] | -8.53470778 | 2.718586 | 96.946352 | 4.07E-07 | 0.000583 |
| 103851 |  |  |  | 2-1303[+] | -9.8621235 | 4.0764944 | 97.876272 | 4.25E-07 | 0.000584 |
| 110658 |  |  |  | 49-438[+] | -6.13448867 | 4.8424905 | 59.036359 | 4.5E-07 | 0.000594 |
| 103128 |  |  |  | 116-475[+] | -1.62602195 | 5.037624 | 56.242583 | 5.95E-07 | 0.000731 |
| 86767 |  |  |  | 2075-2404[-] | -1.01462859 | 5.3818191 | 56.232557 | 5.96E-07 | 0.000731 |
| 111023 |  |  |  | 79-585[+] | -6.5263034 | 4.6159054 | 54.159956 | 8.15E-07 | 0.000966 |
| 18061 | HUNB_DROOR | Protein hunchback | 4.45E-23 | 74-3595[+] | 7.086475018 | 2.2033096 | 59.453352 | 8.71E-07 | 0.000967 |
| 94763 | TSAL_GEOSL | L-threonine ammonia-lyase | 2.36E-52 | 164-1513[+] | 5.42110532 | 3.4828877 | 53.283988 | 8.73E-07 | 0.000967 |

**Table S14.** Number of upregulated (UR) and downregulated (DR) transcripts (FDR < 0.05) for selected pairwise comparisons

|  | **A23 x CA23** | **CA35 x CA23** | **A35 x CA35** | **A35 x A23** | **A35 x CA23** | **P23 x CP23** | **CP35 x CP23** | **P35 x P23** | **P35 x CP35** | **P35 x CP23** | **CA23 x CP23** |
| --- | --- | --- | --- | --- | --- | --- | --- | --- | --- | --- | --- |
| **UR** | 97 | 195 | 85 | 200 | 625 | 157 | 186 | 87 | 80 | 634 | 65 |
| **DR** | 130 | 208 | 191 | 263 | 1025 | 48 | 176 | 104 | 16 | 662 | 97 |
| **Total** | 227 | 403 | 276 | 463 | 1650 | 205 | 362 | 191 | 96 | 1296 | 162 |

**Table S15.** Number of shared upregulated (UR) and downregulated (DR) transcripts (FDR < 0.05) for a selected subset of overlaps between pairwise comparisons

|  | **A23 x CA23 / CA35 x CA23** | **P23 x CP23 / CP35 x CP23** | **A35 x CA35 / P35 x CP35** | **A23 x CA23 / P23 x CP23** | **A35 x A23 / P35 x P23** | **CA35 x CA23 / CP35 x CP23** | **A35 x CA23 / P35 x CP23** |
| --- | --- | --- | --- | --- | --- | --- | --- |
| **UR** | 11 | 19 | 9 | 29 | 60 | 89 | 141 |
| **DR** | 7 | 9 | 10 | 8 | 59 | 71 | 214 |
| **Total** | 18 | 28 | 19 | 37 | 119 | 160 | 355 |

**Table S16.** DE transcripts (FDR < 0.05) shared between injury-alone and heat-alone comparisons for anterior-less worms (A23 x CA23 / CA35 x CA23)

| **Transcript ID** | **UniProtKB / Pfam ID** | **Protein** | **E-value** | **Direction** |
| --- | --- | --- | --- | --- |
| 92163 | ASH2L_HUMAN | Set1/Ash2 histone methyltransferase complex subunit ASH2 | 3.85E-174 | + |
| 97304 | MA2B1_MOUSE | Lysosomal alpha-mannosidase | 5.77E-20 | + |
| 105690 |  |  |  | + |
| 57865 |  |  |  | + |
| 70447 | AGAL_MOUSE | Alpha-galactosidase A | 1.46E-29 | + |
| 79043 | GRP75_PONAB | Stress-70 protein, mitochondrial | 0 | + |
| 111384 | CH10_ORYLA | 10 kDa heat shock protein, mitochondrial | 4.96E-43 | + |
| 68413 |  |  |  | + |
| 74053 | ANM5_MOUSE | Protein arginine N-methyltransferase 5 | 0 | + |
| 39644 | PPBT_BOVIN | Alkaline phosphatase, tissue-nonspecific isozyme | 1.05E-150 | + |
| 104156 | TEBP_RABIT | Prostaglandin E synthase 3 | 9.91E-26 | + |
| 98623 |  |  |  | - |
| 107944 | COPT1_PONAB | High affinity copper uptake protein 1 | 2.58E-22 | - |
| 106668 |  |  |  | - |
| 107656 | SYWC_BOVIN | Tryptophan--tRNA ligase, cytoplasmic | 1.38E-149 | - |
| 71501 | PANK1_ORYSJ | Pantothenate kinase 1 | 5.59E-18 | - |
| 110598 | HPGDS_CHICK | Hematopoietic prostaglandin D synthase | 5.97E-47 | - |
| 51758 | CHS_MELAT | Chitin synthase | 5.08E-140 | - |

**Table S17.** DE transcripts (FDR < 0.05) shared between injury-alone and heat-alone comparisons for posterior-less worms (P23 x CP23 / CP35 x CP23)

| **Transcript ID** | **UniProtKB / Pfam ID** | **Protein** | **E-value** | **Direction** |
| --- | --- | --- | --- | --- |
| 111384 | CH10_ORYLA | 10 kDa heat shock protein, mitochondrial | 4.96E-43 | + |
| 83206 | CH60_CHICK | 60 kDa heat shock protein, mitochondrial | 0 | + |
| 79043 | GRP75_PONAB | Stress-70 protein, mitochondrial | 0 | + |
| 22946 | SYAC_MOUSE | Alanine--tRNA ligase, cytoplasmic | 0 | + |
| 81398 | DAG1_BOVIN | Dystroglycan 1 | 9.97E-08 | + |
| 88942 | CKS1_MOUSE | Cyclin-dependent kinases regulatory subunit 1 | 1.42E-29 | + |
| 101878 | DDX47_HUMAN | Probable ATP-dependent RNA helicase DDX47 | 0 | + |
| 109067 | PHB2_MOUSE | Prohibitin-2 | 5.66E-156 | + |
| 77071 | GLU2B_HUMAN | Glucosidase 2 subunit beta | 1.28E-111 | + |
| 103977 | IPYR_DROME | Inorganic pyrophosphatase | 1.21E-32 | + |
| 12883 | MTMR2_BOVIN | Myotubularin-related protein 2 | 0 | + |
| 32108 | S2611_HUMAN | Sodium-independent sulfate anion transporter | 1.29E-150 | + |
| 93210 | DCOR_RAT | Ornithine decarboxylase | 4.85E-146 | + |
| 5552 |  |  |  | + |
| 80232 | SYEP_DROME | Bifunctional glutamate/proline--tRNA ligase | 0 | + |
| 1461 | SHOC2_XENLA | Leucine-rich repeat protein SHOC-2 | 3.29E-176 | + |
| 99280 | CALM_ELEEL | Calmodulin | 2.31E-82 | + |
| 98282 | SRCH_RABIT | Sarcoplasmic reticulum histidine-rich calcium-binding protein | 1.26E-19 | + |
| 100967 | FNTA_BOVIN | Protein farnesyltransferase/geranylgeranyltransferase type-1 subunit alpha | 2.34E-123 | + |
| 85010 | HGNAT_MOUSE | Heparan-alpha-glucosaminide N-acetyltransferase | 1.16E-117 | - |
| 106791 | KAD1_CHICK | Adenylate kinase isoenzyme 1 | 3.61E-80 | - |
| 111705 |  |  |  | - |
| 109733 | RLT1_RHIO9 | Mucoricin | 2.85E-06 | - |
| 96403 | FCN2_HUMAN | Ficolin-2 | 1.17E-58 | - |
| 110358 | SM20_SCHMA | 20 kDa calcium-binding protein | 9.31E-19 | - |
| 111157 | LYPL1_HUMAN | Lysophospholipase-like protein 1 | 4.68E-75 | - |
| 98223 | CP4F_SHEEP | Prostaglandin E2 omega-hydroxylase CYP4F21 | 4.18E-138 | - |
| 105387 | COFI_OGAPD | Cofilin | 2.45E-36 | - |

**Table S18.** DE transcripts (FDR < 0.05) shared between injury-alone comparisons for both body fragment groups (A23 x CA23 / P23 x CP23)

| **Transcript ID** | **UniProtKB / Pfam ID** | **Protein** | **E-value** | **Direction** |
| --- | --- | --- | --- | --- |
| 96233 | PUR6_CHICK | Multifunctional protein ADE2 | 0 | + |
| 97432 | PUR6_CHICK | Multifunctional protein ADE2 | 0 | + |
| 88942 | CKS1_MOUSE | Cyclin-dependent kinases regulatory subunit 1 | 1.42E-29 | + |
| 108369 | GRN_DICDI | Granulin | 7.03E-06 | + |
| 88157 | ADH_SULAC | NAD-dependent alcohol dehydrogenase | 3.72E-16 | + |
| 106647 | EPDR1_HUMAN | Mammalian ependymin-related protein 1 | 5.47E-05 | + |
| 79013 | MIOX_DANRE | Inositol oxygenase | 1.16E-113 | + |
| 110851 | IF5A_DROME | Eukaryotic translation initiation factor 5A | 1.85E-59 | + |
| 22453 | AAKG2_HUMAN | 5'-AMP-activated protein kinase subunit gamma-2 | 8.59E-76 | + |
| 110968 |  |  |  | + |
| 104447 | HMG2_DROME | High mobility group protein DSP1 | 1.03E-65 | + |
| 35544 | RIR1_MOUSE | Ribonucleoside-diphosphate reductase large subunit | 0 | + |
| 102670 |  |  |  | + |
| 78465 | PF07679.19 | Immunoglobulin I-set domain | 8.10E-06 | + |
| 111063 | H2AX_HUMAN | Histone H2AX | 1.11E-77 | + |
| 111461 | H2AV_XENTR | Histone H2A.V | 4.18E-64 | + |
| 26720 | NLK_HUMAN | Serine/threonine-protein kinase NLK | 2.01E-159 | + |
| 108398 | RET1_RAT | Retinol-binding protein 1 | 9.67E-05 | + |
| 15612 | FKBP4_SPOFR | 46 kDa FK506-binding nuclear protein | 7.49E-28 | + |
| 93326 | HDHD5_HUMAN | Haloacid dehalogenase-like hydrolase domain-containing 5 | 2.68E-94 | + |
| 107200 | AN32A_DROPS | Acidic leucine-rich nuclear phosphoprotein 32 family member A | 5.03E-47 | + |
| 79043 | GRP75_PONAB | Stress-70 protein, mitochondrial | 0 | + |
| 96623 | ALRF2_MOUSE | Aly/REF export factor 2 | 8.71E-47 | + |
| 106189 | PSB7_MOUSE | Proteasome subunit beta type-7 | 6.72E-129 | + |
| 94138 | TCPE_MACFA | T-complex protein 1 subunit epsilon | 0 | + |
| 111384 | CH10_ORYLA | 10 kDa heat shock protein, mitochondrial | 4.96E-43 | + |
| 66139 | DD19A_BOVIN | ATP-dependent RNA helicase DDX19A | 0 | + |
| 109067 | PHB2_MOUSE | Prohibitin-2 | 5.66E-156 | + |
| 92574 | SERA_HUMAN | D-3-phosphoglycerate dehydrogenase | 2.62E-145 | + |
| 98411 | CAH15_MOUSE | Carbonic anhydrase 15 | 1.57E-48 | - |
| 91287 | PUNA_GEOSE | Purine nucleoside phosphorylase 1 | 1.97E-17 | - |
| 80562 | CEL_BOVIN | Bile salt-activated lipase | 3.44E-65 | - |
| 110575 | COPT1_PONAB | High affinity copper uptake protein 1 | 2.71E-19 | - |
| 94480 |  |  |  | - |
| 82046 | S22A3_MOUSE | Solute carrier family 22 member 3 | 7.25E-58 | - |
| 81549 | AQP9_HUMAN | Aquaporin-9 | 1.94E-52 | - |
| 96403 | FCN2_HUMAN | Ficolin-2 | 1.17E-58 | - |

**Table S19.** DE transcripts (FDR < 0.05; max 50 shown) shared between heat-alone comparisons for both body fragment groups (CA35 x CA23 / CP35 x CP23)

| **Transcript ID** | **UniProtKB / Pfam ID** | **Protein** | **E-value** | **Direction** |
| --- | --- | --- | --- | --- |
| 109908 | CRYAB_MACFA | Alpha-crystallin B chain | 1.34E-15 | + |
| 111384 | CH10_ORYLA | 10 kDa heat shock protein, mitochondrial | 4.96E-43 | + |
| 99284 | POSTN_MOUSE | Periostin | 1.81E-17 | + |
| 84988 | TENX_HUMAN | Tenascin-X | 5.16E-29 | + |
| 77307 | HSP7C_SAGOE | Heat shock cognate 71 kDa protein | 0 | + |
| 108040 | U2AF4_RAT | Splicing factor U2AF 26 kDa subunit | 1.51E-122 | + |
| 108614 | CB076_DANRE | UPF0538 protein C2orf76 homolog | 6.33E-30 | + |
| 99029 | CDC37_DROVI | Hsp90 co-chaperone Cdc37 | 3.34E-91 | + |
| 81037 | STIP1_BOVIN | Stress-induced-phosphoprotein 1 | 2.22E-144 | + |
| 105630 | GHITM_HUMAN | Growth hormone-inducible transmembrane protein | 3.40E-108 | + |
| 92507 |  |  |  | + |
| 96808 | UBE2A_MOUSE | Ubiquitin-conjugating enzyme E2 A | 1.06E-82 | + |
| 93864 | AHSA1_HUMAN | Activator of 90 kDa heat shock protein ATPase homolog 1 | 2.96E-106 | + |
| 98668 | F10A1_CHICK | Hsc70-interacting protein | 4.14E-52 | + |
| 103578 | KTR3_YEAST | Probable mannosyltransferase KTR3 | 4.27E-25 | + |
| 20983 | MARH6_MOUSE | E3 ubiquitin-protein ligase MARCHF6 | 0 | + |
| 89924 |  |  |  | + |
| 108657 | CREB_HYDVD | Cyclic AMP-responsive element-binding protein | 6.30E-11 | + |
| 71909 | AHSA1_HUMAN | Activator of 90 kDa heat shock protein ATPase homolog 1 | 2.68E-103 | + |
| 99989 | FKBP4_HUMAN | Peptidyl-prolyl cis-trans isomerase FKBP4 | 1.04E-116 | + |
| 110191 |  |  |  | + |
| 103787 | SRSF7_HUMAN | Serine/arginine-rich splicing factor 7 | 1.66E-19 | + |
| 104156 | TEBP_RABIT | Prostaglandin E synthase 3 | 9.91E-26 | + |
| 77997 | SUV3_DANRE | ATP-dependent RNA helicase SUPV3L1, mitochondrial | 0 | + |
| 82094 | GDIA_MACFA | Rab GDP dissociation inhibitor alpha | 0 | + |
| 86448 | SMCE1_MOUSE | SWI/SNF-related matrix-associated actin-dependent regulator of chromatin subfamily E member 1 | 3.11E-13 | - |
| 110342 |  |  |  | - |
| 109078 | SET_HUMAN | Protein SET | 2.40E-46 | - |
| 84031 | PRUN1_MOUSE | Exopolyphosphatase PRUNE1 | 2.20E-24 | - |
| 105147 | HMG2_DROME | High mobility group protein DSP1 | 3.20E-59 | - |
| 85106 | HSP83_DROMI | Heat shock protein 83 | 7.41E-45 | - |
| 110541 | ATPO_PIG | ATP synthase subunit O, mitochondrial | 1.82E-70 | - |
| 111993 |  |  |  | - |
| 110722 |  |  |  | - |
| 110084 | PDLI7_HUMAN | PDZ and LIM domain protein 7 | 3.36E-11 | - |
| 106037 | PF17064.8 | Sleepless protein | 3.60E-07 | - |
| 106668 |  |  |  | - |
| 99527 | BASI_CHICK | Basigin | 1.31E-17 | - |
| 100763 | FIBA_APOPA | Fibrinogen-like protein A | 2.02E-26 | - |
| 110596 |  |  |  | - |
| 110324 |  |  |  | - |
| 101142 | PANK1_ORYSJ | Pantothenate kinase 1 | 2.50E-11 | - |
| 111639 |  |  |  | - |
| 111542 |  |  |  | - |
| 109885 | SM16_SCHMA | 16 kDa calcium-binding protein | 1.01E-20 | - |
| 109879 |  |  |  | - |
| 69592 |  |  |  | - |
| 49152 | ESN_DROME | Protein espinas | 1.17E-05 | - |
| 75628 | YJX4_SCHPO | CRAL-TRIO domain-containing protein C23B6.04c | 5.82E-12 | - |
| 109244 | QCR1_MOUSE | Cytochrome b-c1 complex subunit 1, mitochondrial | 1.09E-99 | - |

**Table S20.** Oligonucleotide primer sequences used for qPCR

| **Transcript accession** | **Predicted protein** | **qPCR forward primer** | **qPCR reverse primer** |
| --- | --- | --- | --- |
| 79043 | mortalin (stress-70 protein, mitochondrial) | CTGCCATCAGAAGAGTGCGA | CGTGGCCTTTCGTATCGAGT |
| 111384 | hsp10 (10 kDa heat shock protein, mitochondrial) | GTTAGGCGTTAGCAGTCGGT | CGTTCCACAAGAATGCGGTC |
| 103489 | GAPDH | CTTCACCAGATGCGCCAATG | GCCAATGGAGCTAGGCAGTT |
